# Independent evolution of ipecac alkaloid biosynthesis

**DOI:** 10.1101/2024.09.23.614470

**Authors:** Maite Colinas, Clara Morweiser, Olivia Dittberner, Bianca Chioca, Ryan Alam, Helena Leucke, Yoko Nakamura, Delia Ayled Serna Guerrero, Sarah Heinicke, Maritta Kunert, Jens Wurlitzer, Kerstin Ploss, Benke Hong, Veit Grabe, Adriana A. Lopes, Sarah E. O’Connor

## Abstract

Ipecac alkaloids are medicinal monoterpenoid-derived tetrahydroisoquinoline alkaloids found in two distantly related plants: *Carapichea ipecacuanha* (Gentianales) and *Alangium salviifolium* (Cornales). We have elucidated ipecac alkaloid biosynthesis in both species, conclusively demonstrating that biosynthesis of the structurally complex ipecac alkaloid protoemetine has evolved independently. We show that although protoemetine biosynthesis proceeds via the same chemical logic in both species, each plant uses a distinct monoterpene precursor. Moreover, we provide evidence that both plants initiate ipecac biosynthesis by a non-enzymatic Pictet-Spengler reaction, and we elucidate the biosynthetic fate of both the 1*R* and 1*S* stereoisomers that are produced in this non-stereoselective reaction. Phylogenetic analyses clearly show independent pathway evolution through parallel and convergently evolved enzymes. This work provides insight into how nature can capitalize on highly reactive starting substrates, the manner in which multi-step pathways can arise, and also lays the foundation for metabolic engineering of these important medicinal compounds.

## Introduction

Plants produce an enormous diversity of natural products or specialized metabolites. Although natural product pathways are typically lineage-specific, in some cases evolutionarily distant plants independently evolved pathways to synthesize the same molecule (Negin & Jander, 2023; Weng, 2014). Examples include glucosinolates (Rodman *et al*, 1998), benzoxazinoids (Florean *et al*, 2023), caffeine (Huang *et al*, 2016; O’Donnell *et al*, 2021; Vignale *et al*, 2024), cannabinoids (Berman *et al*, 2023) and cardenolides (Kunert *et al*, 2023; Younkin *et al*, 2024; Zust *et al*, 2020). Ipecac alkaloids are monoterpenoid-derived tetrahydroisoquinoline alkaloids that occur in *Carapichea ipecacuanha* (Gentianales order) and *Alangium salviifolium* (Cornales order). Both species are known medicinal plants: ipecac syrup made from *C. ipecacuanha* rhizomes has been used as a vomit inducing medicine in cases of intoxication, while *A. salviifolium*, also known as Ankol(a), is used both as an emetic and in traditional ayurvedic medicine to treat a variety of diseases (Hu *et al*, 2020; Seger & Muelenbelt, 2004). The active emetic ingredients are the tetrahydroisoquinoline alkaloid pathway products, cephaeline and emetine, both derived from protoemetine (Fig. 1). Additionally, anti-cancer and anti-malarial activities have been described for other protoemetine-derived alkaloids such as tubulosine (Kim *et al*, 2020; Sauvain *et al*, 1996). Since *Cornales* and *Gentianales* are estimated to have diverged ca. 150 million years ago (Zuntini *et al*, 2024), it is likely that the protoemetine pathway has evolved independently. Thus, these plants serve as excellent models to study the evolution of a complex medicinal alkaloid biosynthesis pathway.

**Fig. 1.**
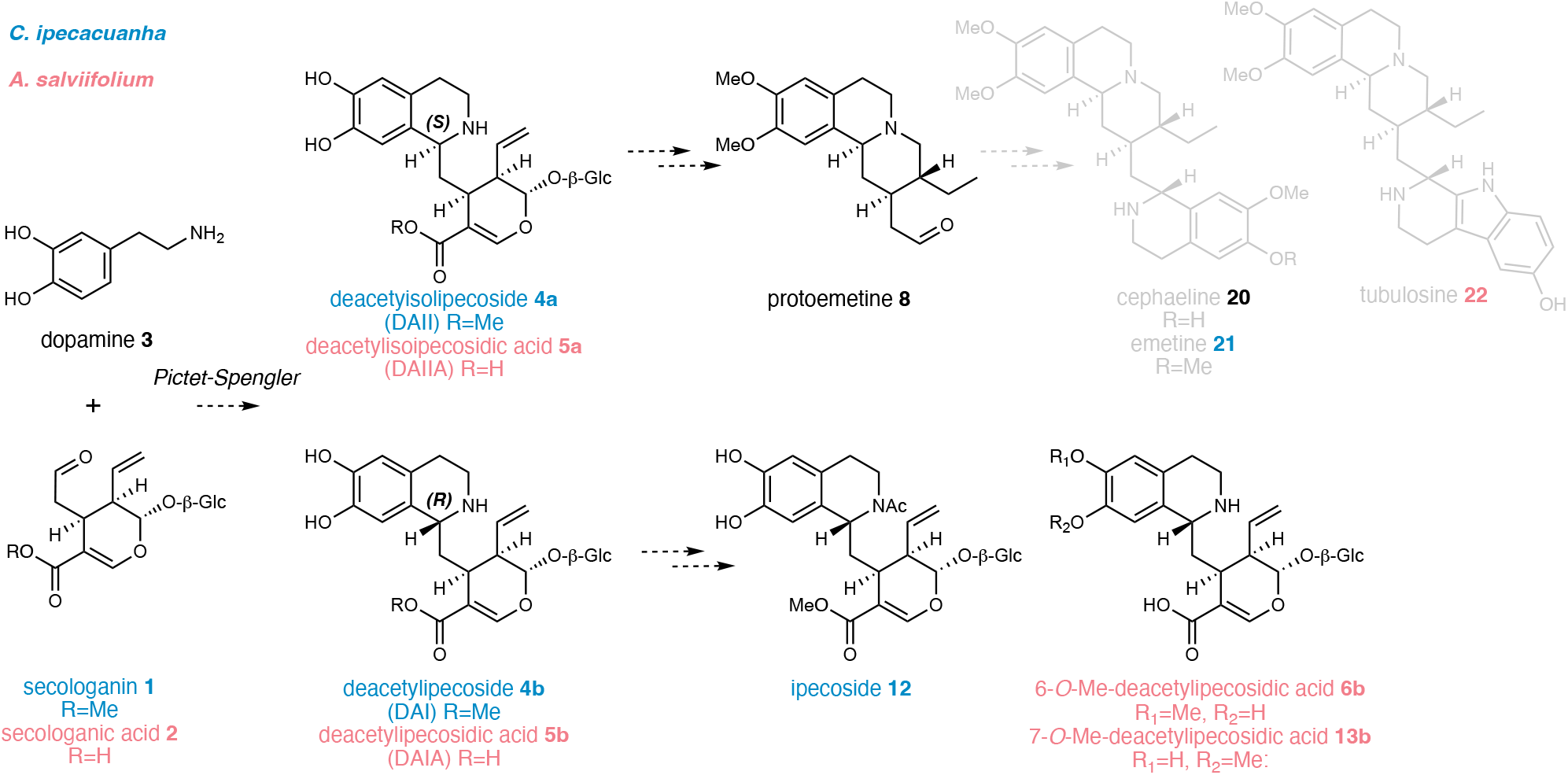
Ipecac alkaloid biosynthesis in *C. ipecacuanha* and *A. salviifolium* based on the literature. Either secologanin **1** or secologanic acid **2** are coupled with dopamine **3** in a Pictet-Spengler reaction to yield deacetylisoipecoside DAII (*S*-epimer) **4a** and deacetylipecoside DAI (*R*-epimer) **4b** or the respective acids deacetylisoipecosidic acid DAIIA (*S*-epimer) **5a** and deacetylipecosidic acid DAIA **5b**. Derivatized forms of the *R*-epimer are *N*-acetylated ipecoside found in *C. ipecacuanha* and 6– or 7-*O*-Me-deacetylipecosidic acid found in *A. salviifolium*. The *S*-epimer undergoes a series of reactions including methylations, deglycosylation, reduction, and in the case of deacetylisoipecoside, deesterification to form protoemetine **8**, which is then derivatized to form downstream alkaloids in both plants as shown. Compounds specific for *C. ipecacuanha* are shown in blue; compounds specific for *A. salviifolium* are shown in magenta.

In *C. ipecacuanha*, ipecac alkaloid biosynthesis begins with a Pictet-Spengler reaction that couples monoterpenoid secologanin **1** with dopamine to generate the initial tetrahydroisoquinoline scaffold. Conflicting literature suggests that either secologanin **1** or secologanic acid **2** could be similarly conjugated with dopamine **3** in *A. salviifolium* (DeEknamkul *et al*, 1997; Itoh *et al*, 2001; Jain *et al*, 2002; Nagakura *et al*, 1978), though secologanic acid has been shown to be the monoterpene precursor in indole alkaloid biosynthesis in *Camptotheca acuminata*, also in the Cornales lineage (Kang *et al*, 2021; Sadre *et al*, 2016). Although all previously identified enzymes that catalyze the Pictet-Spengler reaction are stereoselective, unusually, both *S* and *R* stereoisomers of the initial tetrahydroisoquinoline Pictet-Spengler product are observed in *C. ipecacuanha* (deacetylisoipecoside DAII **4a** [*S* epimer] and deacetylipecoside DAI **4b** [*R* epimer]) (Battersby *et al*, 1978) (Fig. 1). Similarly, methylated forms of both *S* and *R* stereoisomers of the corresponding acids (deacetylisoipecosidic acid, DAIIA, **5a** (*S* epimer) and deacetylipecosidic acid DAIA, **5b,** (*R* epimer)) are observed in *A. salviifolium* (Itoh *et al*., 2001; Tanahashi *et al*, 2000). Early work suggested that a Pictet-Spenglerase enzyme is present in *A. salviifolium* but a gene encoding this enzyme has never been identified (De-Eknamkul *et al*, 2000; DeEknamkul *et al*., 1997). In both *C. ipecacuanha* and *A. salviifolium*, the *S*-epimers are converted via *O*-methylation, deglycosylation, reduction and decarboxylation to protoemetine **8**, an ipecac alkaloid common to both species (Battersby *et al*, 1959; Battersby & Harper, 1959; Itoh *et al*, 2000). In *C. ipecacuanha*, protoemetine is converted into cephaeline and emetine (Battersby *et al*., 1959), while in *A. salviifolium*, proteoemetine is converted to cephaeline, alangimarckine and tubulosine (Albright *et al*, 1965; Battersby *et al*, 1965). In *C. ipecacuanha* the *R*-epimer is *N*-acetylated to ipecoside **12** (Battersby *et al*, 1967), while in *A. salviifolium* the *R* epimer is converted to 6– and 7-*O*-Me-DAIA **6b** and **13b** (Tanahashi *et al*., 2000) (Fig. 1).

A few ipecac alkaloid biosynthetic genes from *C. ipecacuanha* encoding glucosidases and *O*-methyltransferases (*OMT*) have been reported (Cheong *et al*, 2011; Nomura & Kutchan, 2010; Nomura *et al*, 2008), but ipecac biosynthesis remains largely unknown. Here, we report the complete discovery of protoemetine biosynthesis in both *C. ipecacuanha* and *A. salviifolium*, which we show proceeds via an unexpected order of enzymatic reactions. We provide indirect evidence that the Pictet-Spengler reaction initiating the pathway most likely occurs spontaneously in the vacuole, which explains the presence of both 1*R* and 1*S* stereoisomers. While protoemetine is derived from the 1*S* epimer, we also identify biosynthetic genes that derivatize the 1*R* epimer in *C. ipecacuanha* and *A. salviifolium*. A phylogenetic comparison of the respective pathway enzymes indicates that the pathways evolved independently through means of parallel and convergent evolution. This collection of metabolic pathways provides a striking example of independent evolution of complex, medicinally important compounds in plants.

## Results

### Metabolite profiling of *C. ipecacuanha* and *A. salviifolium* tissues

We isolated *C. ipecacuanha* and *A. salviifolium* tissues for RNAseq data and for metabolomic profiling. Protoemetine (**8**) (1*S* stereoisomer) derived alkaloids were found in C*. ipecacuanha* rhizomes (cephaeline and emetine) and in *A. salviifolium* root (cephaeline) (Fig. 2a). The 1*R* derived products were also found in C*. ipecacuanha* rhizomes (ipecoside, **12**), and *A. salviifolium* root (6-*O*-Me-DAIA, **6b**) (Fig. 2a, d). Tissue specific metabolite profiling (Fig. 2 f, g, Supplementary Fig. 1-2, comparisons to standards shown in Supplementary Fig. 3-7) revealed that *C. ipecacuanha* accumulates similar amounts of ipecac alkaloids in both young leaves and rhizome (Fig. 2f). In contrast, *A. salviifolium* pathway intermediates up to protoemetine **8** are detected in high levels in leaf buds, but cephaeline only accumulates in roots and bark of older stems (Fig. 2g). We used these metabolite profiles to guide the search for protoemetine gene candidates in the corresponding RNAseq datasets.

**Fig. 2.**
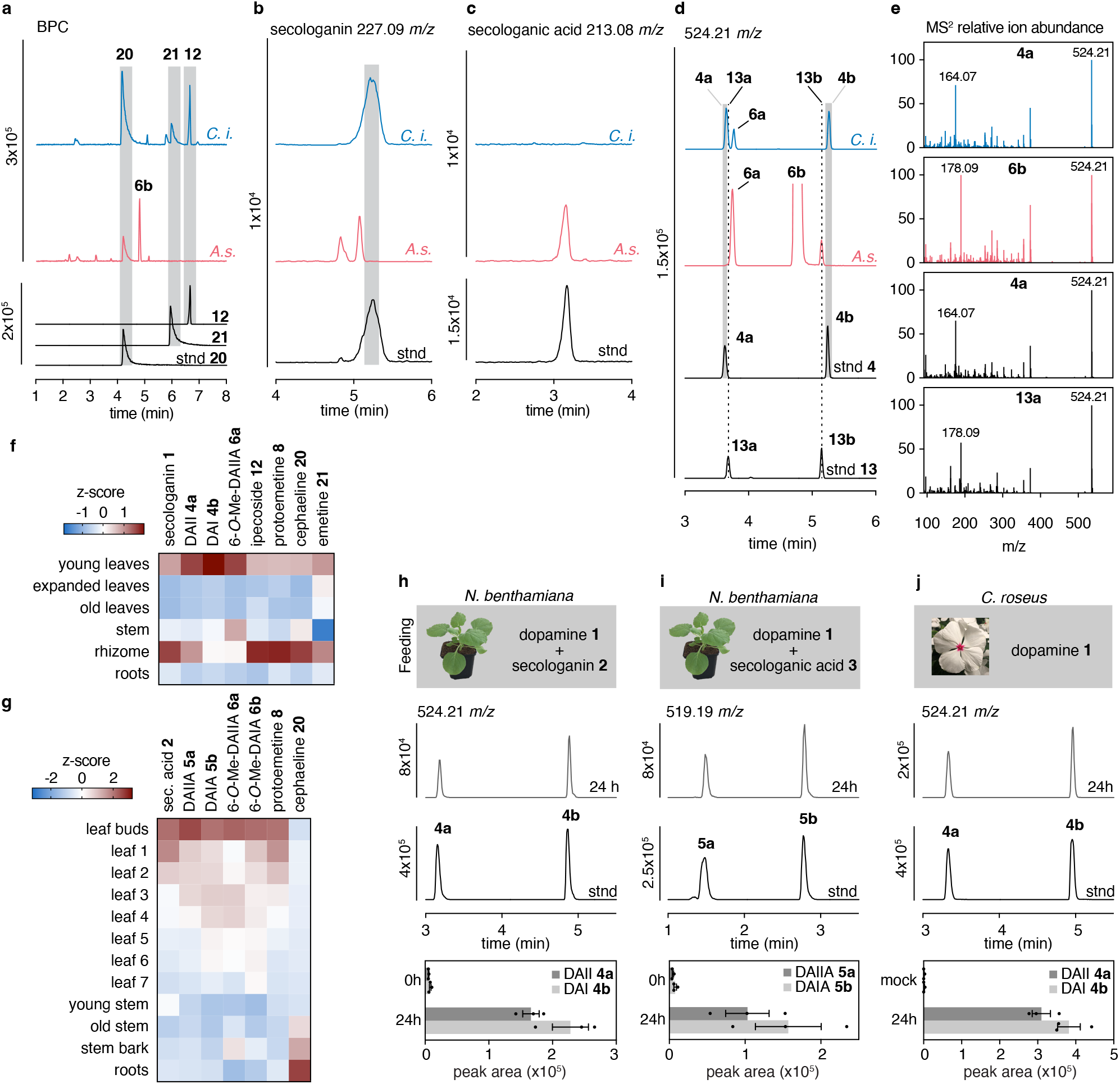
Metabolite profiling and evidence for a non-enzymatic Pictet-Spengler reaction in *C. ipecacuanha* and *A. salviifolium*. **a-d**, LC-MS analysis of *C. ipecacuanha* rhizome extracts (blue) and *A. salviifolium* root extracts (magenta). **a,** Base Peak Chromatograms (BPC) of extracts at 2 mg FW/ml. Major ipecac alkaloids are indicated. Authentic standards for cephaeline, emetine and ipecoside were run alongside as indicated. See Supplementary Fig. 3 for full characterization. **b-d,** Extracted Ion chromatograms (EICs) of extracts at 10 mg FW/ml. **b**, EIC of secologanin ([M-glucose+H]^+^ = 227.09 *m/*z) indicates that it is only found in *C. ipeacuanha*. **c,** EIC of secologanic acid ([M-glucose+H]^+^ = 213.08 *m/*z) indicates that it accumulates only in *A. salviifolium*. See Supplementary Fig. 4 for full characterization. **d**, **e**, DAI/I (**4a**,**b**) and *O*-Me-DAI/IA have identical *m/z* [M+H]^+^ = 524.21 but are distinguished by different retention times (**d**) and MS^2^ fragmentation (**e**). DAI/I (**4a/b**) is exclusively found in *C. ipecacuanha* and not *in A. salviifolium* where 6-*O*-Me-, and 7-*O*-Me-DAIA (1*R*) (**6b** and **13b**) accumulate to large amounts. A putative intermediate on route to protoemetine biosynthesis, 6-*O*-Me-DAIIA **6a** (1*S*) is found in both species. See Supplementary Fig. 5 for full characterization. Assignment of **4a** as 1*S* and **4b** as 1*R* was confirmed by nuclear magnetic resonance (NMR) spectroscopy (Supplementary information). **f-g**, Tissue specific relative distribution of selected ipecac alkaloids in *C. ipecacuanha* (**f**) and *A. salviifolium* (**g**) showing that in addition to roots or rhizome, young leaf tissues of both species also appear to be locations of high ipecac alkaloid accumulation. Heatmaps depict Z-scores of average peak areas of three replicates for each metabolite (Supplementary Fig. 1-2). **h-i,** Reaction products DAI/I or DAI/IA of coupling reaction of dopamine with secologanin (**h**) or secologanic acid (**i**) are observed after within 24 hours after infiltration into *N. benthamiana* leaves. **j**, Infiltration of dopamine to flower petals of the natural secologanin producer *Catharanthus roseus* also leads to appearance of DAI/I also within 24 hours. Stnd, standard.

### Ipecac alkaloid biosynthesis is initiated with a Pictet-Spengler reaction

In the metabolite profiling experiments described above, we also noted that secologanin (**1)** is only observed in *C. ipecacuanha* while secologanic acid (**2)** is observed only in *A. salviifolium* (Fig. 2 b,c,). This suggests that *C. ipecacuanha* uses secologanin in ipecac biosynthesis, while *A. salviifolium* uses the corresponding acid. Consistent with this, DAII **4a** and DAI **4b** are detected in *C. ipecacuanha* but not in *A. salviifolium* (Fig. 2 d,e). To our surprise, we noticed that when we infiltrated the starting substrates– secologanin or secologanic acid together with dopamine– into *Nicotiana benthamiana* leaves, the corresponding Pictet-Spengler products DA(I)I or DAI(I)A (both 1-*S* and 1-*R* stereoisomers) were formed within 24 hours (Fig. 2h, i). Moreover, when *Catharanthus roseus* flower petals, which contain endogenous secologanin **1** but do not produce ipecac alkaloids, were infiltrated with dopamine, accumulation of DAI **4b** (1*R*) and DAII **4a** (1*S*) were also observed after 24 hours (Fig. 2j), clearly demonstrating that this non-enzymatic reaction can take place *in planta*. Dopamine is a highly activated substrate for the Pictet-Spengler reaction, reacting rapidly in a non-stereo-selective manner under mild acidic conditions (Menendez-Perdomo & Facchini, 2023). Although enzymes that catalyze the Pictet-Spengler reaction with dopamine are known, these enzymes are always stereoselective (Lee & Facchini, 2010). Since *C. ipecacuanha* and *A. salviifolium* produce products derived from both stereoisomers, and since we demonstrated that this reaction can occur in two different plants without a dedicated enzyme, it seems reasonable to assume that this Pictet-Spengler coupling occurs non-enzymatically and non-stereoselectively in these plants.

### Identification of ipecac alkaloid biosynthetic gene candidates

After the Pictet-Spengler reaction, protoemetine is generated from DAII (**4a)** or DAIIA (**5a)** (1*S* epimers) by *O*-methylation, deglycosylation, reduction and, in the case of *C. ipecacuanha*, de-esterification (Fig. 1). The 1*R*-epimers (DAIIA **5a** or DAIA **5b**) are either acetylated to form ipecoside (**12)** in *C. ipecacuanha* or *O*-methylated to form 6-OMe-DAIA **6b** and 7-OMe-DAIA **13b** in *A. salviifolium*. We searched for genes in our generated transcriptomes that encode enzymes that could catalyze these reactions. First, we identified the previously published *C. ipecacuanha OMT* genes (renamed *CiDOMT1, CiDOMT2, CiDPOMT*) and glucosidase (*CiDGD*) (Cheong *et al*., 2011; Nomura & Kutchan, 2010; Nomura *et al*., 2008). Expression levels of the three *OMT* genes in our *C. ipecacuanha* transcriptome was highest in young leaves and rhizome (Supplementary Fig. 8a), consistent with the metabolite profile of ipecac alkaloid accumulation (Fig. 2f). Using the previously discovered *CiDOMT1* gene as bait for coexpression analysis, we identified two reductase and three esterase transcripts. Surprisingly, the previously reported glucosidase gene, *CiDGD* did not coexpress with the *CiOMT* genes (–0.49 Pearson correlation with *CiDOMT1*), but among the highly co-expressed contigs a new glucosidase, named *CiS6DGD* (71.4 % amino acid identity to CiDGD), was found. A gene annotated as an acetyl transferase was also identified.

Additionally, we noticed that orthologs of known precursor generating enzymes secologanin synthase (*SLS*)– which catalyzes the final step of secologanin biosynthesis– and tyrosine decarboxylase (*TyrDC*)– predicted to be involved in dopamine biosynthesis– were also tightly coexpressed with *CiDOMT1*.

No pathway genes from *A. salviifolium* have been reported, and the *A. salviifolium* transcriptome did not contain orthologs of the previously published *CiOMT* genes. Therefore, we mined the *A. salviifolium* transcriptome for orthologs of putative precursor genes *TyrDC* and secologanic acid synthase (*SLAS*), which catalyzes the last step of secologanic acid biosynthesis (Yang *et al*, 2019). The expression profile of these two identified orthologs were highest in leaf buds and roots (Supplementary Fig. 8b), consistent with metabolite data (Fig. 2g). Using *TyrDC* as a bait for coexpression analysis, we identified coexpressed genes that had functional annotations consistent with *O*-methyltransferase, dehydrogenase and glycosyl hydrolase activity. Since metabolite data indicated that *A. salviifolium* utilizes secologanic acid to form the initial Pictet-Spengler product DAIIA, we predicted that an esterase would not be required for protoemetine biosynthesis in this plant.

### Comparative discovery of protoemetine 8 biosynthesis

Transient expression of pathway gene candidates along with infiltration of the starting substrate in *N. benthamiana* enabled us to successfully deconvolute the complete protoemetine (**8**) biosynthetic pathway from both *C. ipecacuanha* and *A. salviifolium* (Fig. 3 and Supplementary Fig. 9-12). Combinatorial expression of the identified gene candidates was followed by assay of individual genes. Based on previous proposals and also the chemical logic established for other secologanin-derived natural products such as corynantheal (*Chinchona pubescens*) (Nomura & Kutchan, 2010; Nomura *et al*., 2008; Trenti *et al*, 2021), we initially hypothesized that in *C. ipecacuanha*, DAII (**4a**) would be deglycosylated, reduced and de-esterified, which would in turn lead to spontaneous decarboxylation to form protoemetine. However, since we observed DAIIA (**5a**) and 6-*O*-Me-DAIIA (**6a**) in *C. ipecacuanha* (Fig. 2f, Supplementary Fig. 2), we speculated that deglycosylation may happen after de-esterification. Indeed, the identified esterase (CiDE) de-esterifies the glucosylated intermediate DAII (**4a**) to yield DAIIA (**5a**) (Fig. 3, Supplementary Fig. 10). In *A. salviifolium* the pathway directly starts with de-esterified DAIIA, derived from the Pictet-Spengler condensation of secologanic acid (**2**) with dopamine. Thus, both *C. ipecacuanha* and *A. salviifolium* utilize DAIIA (**5a**) as a protoemetine (**8**) intermediate.

**Fig. 3.**
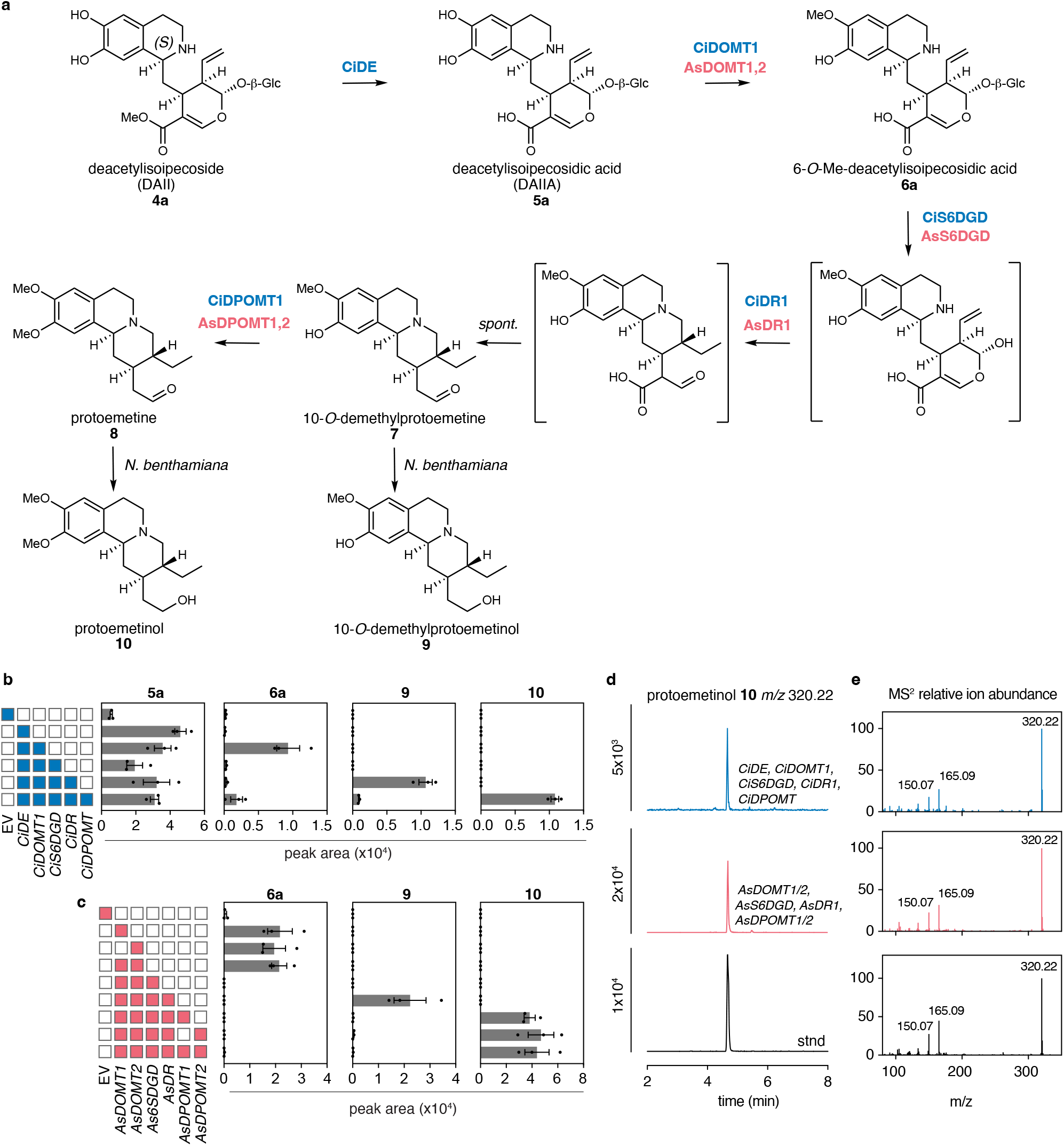
Discovery of *C. ipecacuanha and A. salviifolium* protoemetine biosynthetic genes. **a**, The complete pathway leading to protoemetine (**8**). Molecules in parentheses are hypothesized unstable intermediates that were not detected by LC-MS. Aldehydes **7** and **8** are only detected in traces by LC-MS; instead the corresponding alcohols **9** and **10** are detected. Peak identities were confirmed by comparing to standards (see Supplementary information for full MS characterization and NMR characterization of protoemetine standard), except 10-*O*-demethyl-protoemetinol, which was identified based on MS^2^ fragmentation (Supplementary Fig. 15). **b,** LC-MS peak areas of products in *N. benthamiana* upon expression of indicated *C. ipecacuanha* pathway genes and infiltration of a synthetically generated mixture of DAII (**4a**) and DAI (**4b**). Data are mean ± SEM of *n* = 3 biological replicates. **c**, LC-MS peak areas of products in *N. benthamiana* upon expression of indicated *A. salviifolium* pathway genes and infiltration of synthetically generated mixture of DAIIA (**5a**) and DAIA (**5b**). Data are mean ± SEM of *n* = 3 biological replicates. **d**, EIC for protoemetinol upon expression of indicated *C. ipecacuanha* pathway genes (blue), or *A. salviifolium* pathway genes (magenta) and authentic standard (stnd, black). **e**, MS^2^ fragmentation for the corresponding peaks shown in **d** confirming the peak as protoemetinol. See Supplementary material for synthesis of the protoemetinol standard. Results of additional gene combinations are shown in Supplementary Fig. 9-12).

DAIIA (**5a**) is then 6-*O*-methylated by CiDOMT1 (previously identified by Nomura) or by AsDOMT1,2 (Fig. 3). Although an authentic standard is not available, the resulting product has an *m/z* value and MS^2^ consistent with 6-*O*-Me-DAIIA (**6b**), and this product is also detected in extracts of both species (Fig. 2d,e, Supplementary Fig. 5). We observed that CiS6DGD/AsS6DGD deglycosylated 6-*O*-Me-DAIIA, whereas the pathway could not be reconstituted with the previously reported *C. ipecacuanha* glucosidase, *CiDGD*, (Fig. 3, Supplementary Fig. 13). The aglycone generated by CiS6DGD/AsS6DGD is subjected to a two-step reduction, catalyzed in both species by a medium chain reductase, CiDR1 or AsDR1 (Fig. 3, Supplementary Fig. 14). Spontaneous decarboxylation is then triggered, yielding 10-*O*-demethylprotoemetine, which is further methylated by CiDPOMT or AsDPOMT1,2. Notably, we could only detect trace amounts of the aldehydes 10-*O*-demethylprotoemetine and protoemetine in *N. benthamiana* (Supplementary Fig. 10,12), but instead detected the reduced forms 10-*O*-demethylprotoemetinol (identified by *m/z* MS^2^ fragmentation Supplementary Fig. 15) and protoemetinol (confirmed by comparison with an authentic standard), which is expected due to endogenous *N. benthamiana* aldehyde reductases (Torrens-Spence *et al*, 2018; Zhang *et al*, 2011). Paralogs of DR (CiDR2 and AsDR2) are also active, but result in formation of lower amounts of protoemetinol (Supplementary Fig. 16,17). Although CiDOMT2 can methylate the 7-hydoxy group of DAI, it does so with low efficiency (Supplementary Fig. 10,18).

### Discovery of species-specific *R*-epimer pathways

Our metabolite profiling suggested that the *R*-epimer DAI (**4b**) is acetylated to form ipecoside (**12**) in *C. ipecacuanha*, while DAIA (**5b**) is methylated on the 6-hydroxy group (and to a lesser extent on the 7-hydroxy group) in *A. salviifolium* to form 6 (or 7)-*O*-Me-DAIA (**6b**, **13b**) (Fig. 2d,e, Supplementary Fig. 2b). A gene annotated as a BAHD type acetyltransferase in the *C. ipecacuanha* transcriptome that was highly co-expressed with *CiDOMT1* (Supplementary Fig. 8a) led to formation of ipecoside (as evidenced by comparison with an authentic standard) when expressed in *N. benthamiana* along with DAI (**4b**) (Fig. 4a-c). This gene was thus named *ipecoside synthase* (*CiIpS*).

**Fig. 4.**
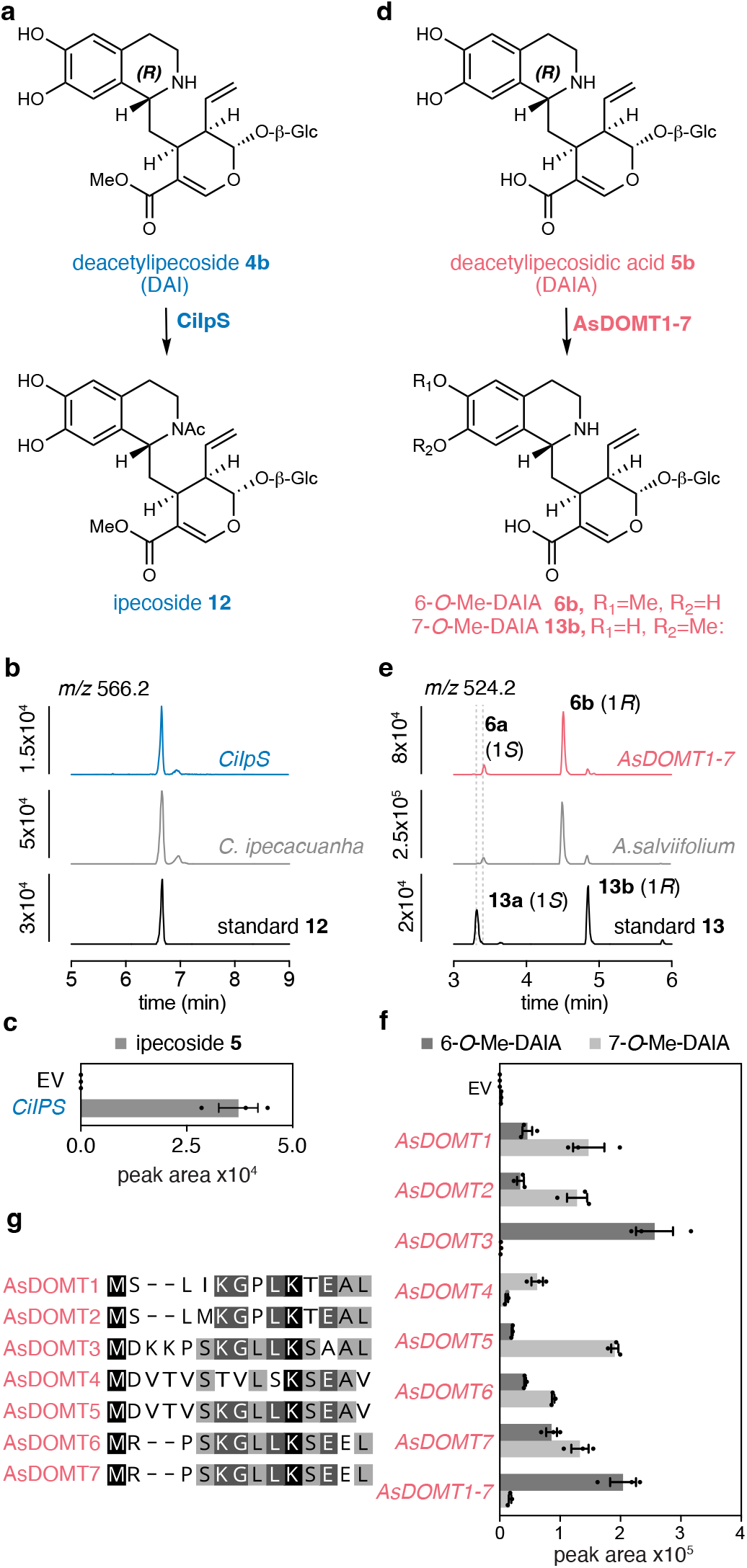
Reconstitution of *C. ipecacuanha* and *A. salviifolium* specific *R*-epimer pathways in *N. benthamiana*. **a**, *C. ipecacuanha* CiIPS *N*-acetylates DAI (**4b**) to yield ipecoside (**12**). **b,** EICs of ipecoside (**12**) (*m/z* 566.22) obtained from *N. benthamaiana* expressing CiIpS and infiltrated with a synthetic mixture of DAI (**4b**) and DAII (**4a**) (blue, top), native *C. ipecacuanha* extract (grey, middle) and authentic standard (black, bottom). **c,** LC-MS peak areas of *N. benthamiana* upon expression of CiIpS and infiltration of DAI (**4b**) compared to empty vector (EV) control. Bars show means of three biological replicates, error bars are standard error of the mean (SEM). **d,** In *A. salviifolium* DAIA (**5b**) is *O*-methylated at 6 or 7 position leading to 6-*O*-Me-DAIA (**6b**) or 7-*O*-Me-DAIA (**13b**) by the closely related enzymes AsDOMT1-7. **e,** EICs from *N. benthamaiana* expressing *AsDOMT1-7* and infiltrated with a synthetic mixture DAIA (**5b**) and DAIIA (**5a**) (magenta, top), native *A. salviifolium* extract (grey) and mixture of synthetic 7-*O*-Me-DAI/IA **13a** and **13b** standards (black, bottom). **f,** peak areas of products obtained from *N. benthamiana* transformed with EV (control) compared to *AsDOMT* expression constructs plus infiltration with a synthetic mixture of DAIA (**5b**) and DAIIA (**5a)**. Bars show mean from three biological replicates is shown, error bars are SEM. AsDOMTs enzymes exhibit different product specificities. **g**, Alignment of the *N*-termini of AsDOMT1-7. Product profiles shown in **f** appear to be largely consistent with the level of sequence identity at the *N*-terminus. Full alignment is shown in Supplementary Fig. 19.

In the *A. salviifolium* transcriptome, seven closely related class I *OMT* genes that showed promising expression profiles were tested (Supplementary Fig. 8b). AsDOMT1 and 2 methylate DAIIA (**5a**) on the 6-hydroxy group on route to protoemetine as described above, but also catalyze 6-, and 7-*O*-methylation of the *R*-epimer DAIA (**5b**) (Fig. 4d-f). An authentic standard of 7-OMe-DAIA was generated through an *in vitro* Pictet Spengler reaction of secologanic acid with 4-*O*-Me-dopamine; the identity of 6-*O*-Me-DAIA was established by comparison with this authentic standard and also appears to be identical with a highly accumulating compound in *A. salviifolium* (Supplementary Fig. 5). AsDOMT3-7 are selective for the *R*-epimer, DAIA (**5b**), but produce different *O*-methylated product profiles (Fig. 4f). A sequence alignment of these seven OMT enzymes reveals a high level of overall sequence identity (83.7 %) (Supplementary Fig. 19) but only 55 % identity at the *N*-terminus (Fig. 4g), a region known to confer substrate and product specificity (Lukacin *et al*, 2004). Indeed, the similarities in substrate and product profiles correlate with the level of identity at the *N*-terminus (Fig. 4f,g). Surprisingly, when all *AsDOMT* genes are expressed together in *N. benthamiana,* their product profile differs from the sum profile expected from expression of single genes; specifically, much higher levels of 6-*O*-Me-DAIA compared to 7-*O*-Me-DAIA are observed, a profile similar to what is detected in the native plants (Fig. 4e). It is possible that upon expression in combination, enzymatic activities influence each other, leading to changed product profiles as previously described (Park *et al*, 2018).

### A vacuolar exporter from *C. roseus* enhances protoemetine biosynthesis from DAII

In reconstitution experiments, the majority of infiltrated DAII (**4a**) starting substrate was not converted into downstream products (Supplementary Fig. 10). In the related monoterpene indole alkaloid pathway, the Pictet-Spengler product produced from secologanin and tryptamine is produced enzymatically by a vacuolar enzyme and then exported into the cytosol by a transporter (CrNPF2.9) (Payne *et al*, 2017). We hypothesized that exogenously supplied DAI/I **4b/4a** may be imported into the vacuole by *N. benthamiana* transporters (de Brito Francisco & Martinoia, 2018; Martinoia *et al*, 2007), rendering these starting materials inaccessible to the downstream cytosolic pathway enzymes. To test whether CrNPF2.9 could export the protoemetine precursor DAII (**4a**), we expressed CrNPF2.9 with *C. ipecacuanha* protoemetine biosynthetic genes in *N. benthamiana*. Gratifyingly, we observed an almost 12-fold relative increase in protoemetinol levels, suggesting that this vacuolar exporter could transport DAII (**4a**) out of the vacuole (Supplementary Fig. 10, 20a). No increase was observed when CrNPF2.9 was expressed in combination with *A. salviifolium* protoemetine pathway genes (Supplementary Fig. 12), suggesting that this transporter does not recognize DAIIA (**5a**). Expression of CrNPF2.9 with CiIpS did not lead to an increase of the *R*-stereoisomer ipecoside, suggesting that this transporter is specific for the *S*-stereoisomer (Supplementary Fig. 20b). Furthermore, expression of the protoemetine biosynthetic genes with uncoupled secologanin and dopamine (as opposed to DAII **4a**) only yielded protoemetinol when CrNPF2.9 was included (Supplementary Fig. 20c). We hypothesize that the glycosylated secologanin would get imported into the vacuole, where the mildly acidic conditions would facilitate the non-enzymatic formation of DAI/I (Supplementary Fig. 20d). Although we tested numerous candidates, the native vacuolar exporters of *C. ipecacuanha* and *A. salviifolium* remain to be discovered.

### Nuclear localized glucosidases deglycosylate *R*-epimer derived products

Glucosidase CiDGD deglycosylates the 1*R*-derived stereoisomer ipecoside (**12**) (Nomura *et al*., 2008) (Fig. 5a,b). In contrast, CiS6DGD, which we showed to be involved in protoemetine biosynthesis, did not turnover ipecoside (Fig. 5b). Assays of CiDGD and Ci6SDGD with substrates DAI/I (**4b**/**4a**), 7-O-Me-DAI/I (**19b/19a**), DAI/IA (**5b/5a**), 6-*O*-Me-DAIIA (**6a**) suggest that CiDGD has broad substrate specificity, whereas the more selective Ci6SDGD most efficiently deglycosylates 6-*O*-Me-DAIIA **6a**, which is on pathway to protoemetine **8** (Supplementary Fig. 21a). Analogously, the *A. salviifolium* glucosidases AsDGD1,2, or AsS6DGD were similarly assayed with the *A. salviifolium* substrates DAI/IA **5a**/**5b,** 7-*O*-Me-DAI/IA **13b/13a**, 6-*O*-Me-DAIIA **6a,** 6-O-Me-DAIA **6b**. AsDGD1 and AsDGD2 consumed 7– and 6-*O*-Me-DAIA entirely (Fig. 5d, Supplementary Fig. 20b), while AsS6DGD only consumed the protoemetine pathway intermediate 6-*O*-Me-DAIIA (Supplementary Fig. 21b). Thus, both *C. ipecacuanha* and *A. salviifolium* seem to have a protoemetine pathway specific glucosidase along with a glucosidase with broader substrate specificity. The substrate specificity of CiS6DGD and AsS6DGD require that DAIIA is subjected to 6-*O*-methylation prior to deglycosylation. In contrast, CiDGD and AsDGD1,2 deglycosylate DAII(A) directly, which prevents formation of protoemetine (Supplementary Fig. 13, 21).

**Fig. 5.**
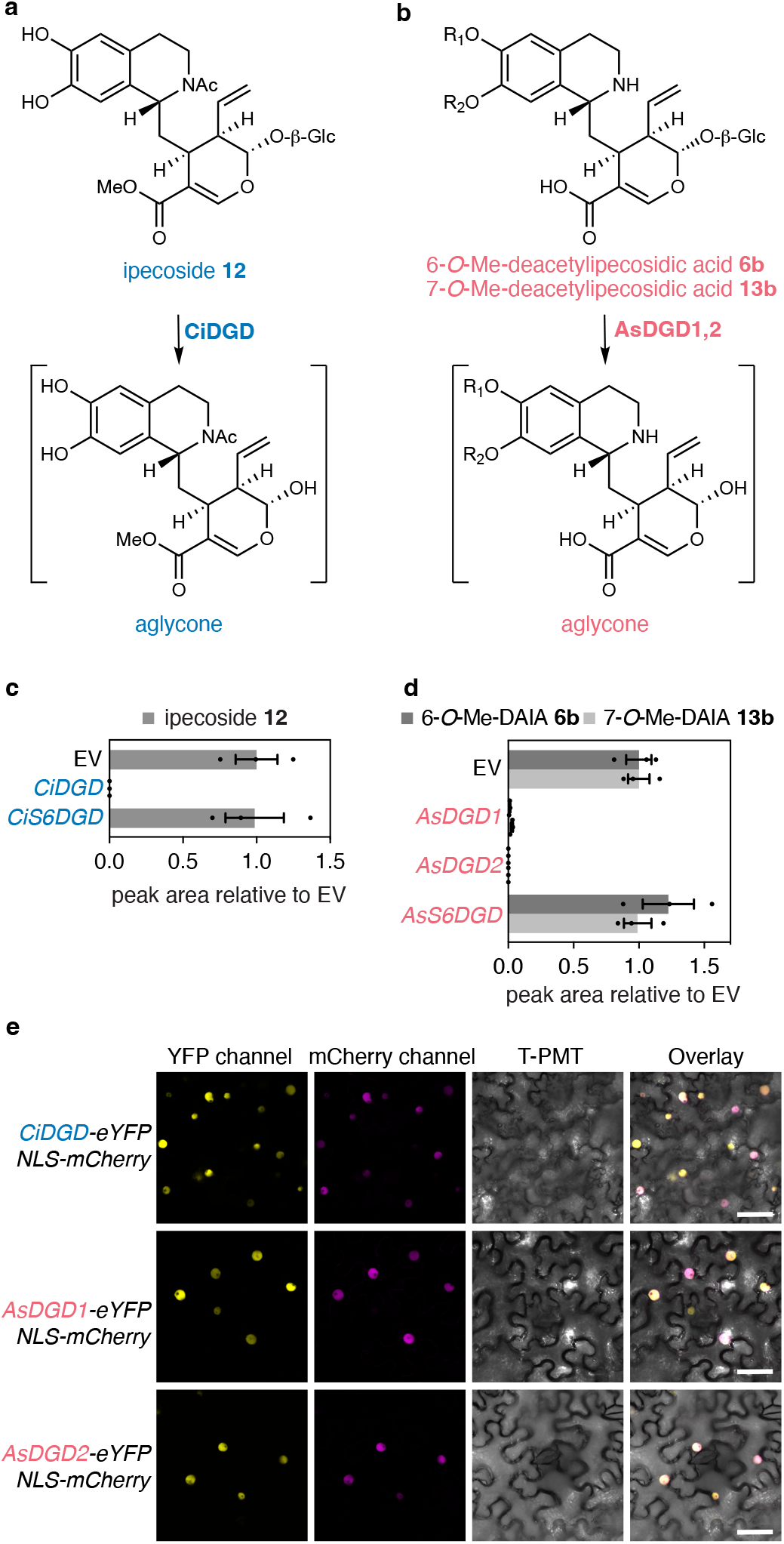
Deglycosylation of *R* epimer derived products by nuclear localized glucosidases. **a**, CiDGD deglycosylates ipecoside. **b**, *N. benthamiana* leaf disks expressing *CiDGD* or *CiS6DGD* incubated with ipecoside (**12**) for 24 hours. Ipecac is consumed when *CiDGD* is expressed but not with *CiS6DGD*. results of incubation with additional substrates are shown in Supplementary Fig. 21a. Bars show the mean peak areas normalized to peak areas in EV control of three biological replicates, error bars are standard error of the mean (SEM). **c**, AsDGD1,2 deglycosylate *O*-methylated DAIA *R*-epimers. **d**, Incubation of 6-*O*-Me-DAIA (**6b**, light grey, produced *in vitro* by recombinant AsDOMT3, see methods for details) or 7-*O*-Me-DAIA (**13b**) (in mix with 7-*O*-Me-DAIIA, **13a**) to *N. benthamiana* leaf disks expressing *AsDGD1* or *2* or *As6SDGD*. AsDGD1,2 deglycosylate these substrates but AsS6DGD does not. Bars show the mean peak areas normalized to peak areas in EV control of three biological replicates, error bars are SEM. Results of incubation with additional DAIA derivatives are shown in Supplementary Fig. 21b. **e**, Confocal laser scanning microscopy of *N. benthamiana* leaves co-expressing *C*-terminal eYFP fusions of CiDGD, AsDGD1 or AsDGD2 along with mcherry fused to a Nuclear Localization Sequence (NLS). The data clearly show nuclear localization of these enzymes. Scale bars indicate 50 µm. Additional subcellular localization data are shown in Supplementary Fig. 23.

CiDGD, and AsDGD1,2 each contain a *C*-terminal bipartite nucleus localization sequence (NLS) (as predicted with DeepLoc2.0) (Thumuluri *et al*, 2022) (Supplementary Fig. 22). When these genes were fused to eYFP, each co-localized with an mcherry-NLS marker when expressed in *N. benthamiana* leaves (Ivanov & Harrison, 2014) (Fig. 5e and Supplementary Fig. 23). *C*-terminal tagged AsDGD1,2 showed localization across the entire nucleus whereas the *N*-terminal tagged fusions were localized to a smaller compartment within the nucleus, which could be aggregates, as has been previously proposed for the glucosidase involved in monoterpene indole alkaloid biosynthesis (Guirimand *et al*, 2010) (Supplementary Fig. 23). Taken together these results indicate that both species derivatize *R*-epimers in a species dependent manner and contain highly active nuclear localized glucosidases with relatively broad substrate specificity. The protoemetine specific glucosidase CiS6DGD, which is highly similar to CiDGD, also showed nuclear localization (Fig. 5e). The protoemetine specific glucosidase from *A. salviifolium* lacked the NLS and was localized to both nucleus and cytosol (Supplementary Fig. 23).

### Ipecac alkaloid biosynthetic genes evolved independently in the Gentianales and the Cornales

Having elucidated ipecac alkaloid biosynthesis in these two distantly related plants, we performed phylogenetic comparisons of the identified enzymes (Fig. 6, Supplementary Fig. 24-26). These analyses clearly demonstrated that *A. salviifolium* (Cornales) and *C. ipecacuanha* (Gentianales) enzymes evolved independently. AsDOMT enzymes belong to the class I Mg^2+^ dependent Caffeoyl CoA 3-O-methyltransferase family (Liu *et al*, 2022) (Fig. 6a). AsDOMT1-7 form a separate subclade suggesting that the different substrate and product profiles observed among these enzymes (Fig. 4) likely arose through tandem gene duplication and subfunctionalization. All other identified OMT enzymes are class II Mg^2+^ independent, which form two well separated clades, in which one clade contains AsDPOMTs and the other contains CiDOMTs and CiDPOMT. All three *C. ipecacuanha* OMTs are closely related suggesting that these genes arose through tandem duplications and subsequent neofunctionalization to catalyze *O*-methylation of either DAIIA or 10-*O*-demethylprotoemetine.

**Fig. 6.**
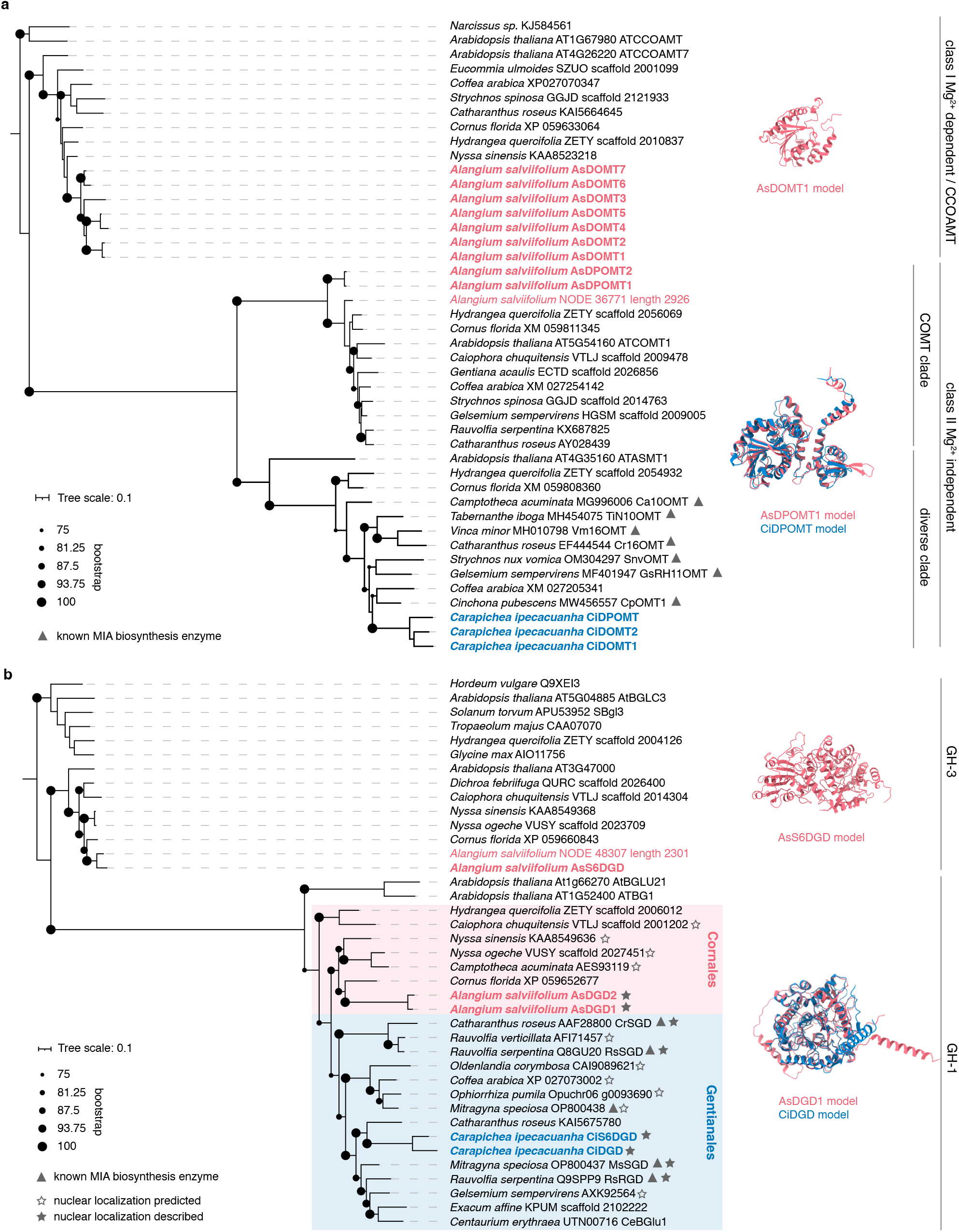
Ipecac alkaloid biosynthesis arose independently in *C. ipecacuanha* and *A. salviifolium*. Maximum likelihood phylogenetic trees of amino acid sequences of pathway enzymes and homologs from other Cornales and Gentianales species. Alphafold3 models of representative pathway enzymes are shown alongside the different enzyme classes. **a,** *O*-methyltransferases. Analyzed sequences cluster with different known classes and subclades of OMTs. AsDOMTs form part of the clade of Caffeoyl CoA 3-*O*-methyltransferases (CCoAOMTs) which are Mg^2+^ dependent class I OMTs well known to be involved in lignin biosynthesis. All other OMTs of this study are class II Mg^2+^ independent class II OMTs. Both classes share a common domain with the same fold but class II OMTs contain an additional domain. Class II OMT sequences form two clades: a clade containing OMTs with high sequence similarity and predicted to have caffeic acid *O*-methylation (COMT) activity and a clade containing OMTs with less sequence similarity and diverse functions (separation of clades has been previously observed for example (Salim *et al*, 2018). AsDPOMTs are found in the COMT clade while CiDOMTs and CiDPOMT are part of the diverse clade. A tree containing all bootstrap values is shown in Supplementary Fig. 25. **b**, Glucosidases. The protoemetine pathway specific AsS6DGD is a member of the GH-3 family and thus has a different protein fold than the other glucosidases characterized in this study, which all of the GH-1 type. The GH-1 *A. salviifolium* and *C. ipecacuanha* GH-1 sequences cluster with sequences from the Cornales or the Gentianales order, respectively, suggesting parallel evolution. A tree containing all bootstrap values is shown in Supplementary Fig. 26. Enzyme names were are shown if available. ATASAMT, *A. thaliana* N-acetylserotonin O-methyltransferase, Ca10OMT, *C. acuminata* 10-hydroxycamptothecin *O*-methyltransferase, TiN10OMT, *T. iboga* noribogaine 10-O-methyltransferase, Vm16OMT, *V. minor* 16-hydroxyvincadifformine 16-O-methyltransferase, Cr16OMT, *C. roseus* 16-hydroxytabersonine O-methyltransferase, SnvOMT, *S. nux-vomica* strychnine O-methyltransferase, GsRH11OMT, *G. sempervirens* rankinidine/humatenine-11-O-methyltransferase, CpOMT1, *C. pubescens* quinine O-methyltransferase, AtBGCL3, *A. thaliana* beta-glucosidase 3, SBgl3, *S. torvum* furastonol glycoside glucosidase, AtBGLU21, *A. thaliana* beta-glucosidase 21, AtBG1, *A. thaliana* beta glucosidase 1, SGD, strictosidine glucosidase, RsRGD, *R. serpentina* raucaffricine glucosidase, CeBGlu1, *C. erythraea* beta-glucosidase. All sequences included in the tree are listed in Supplementary Dataset 1.

The protoemetine specific CiS6DGD belongs to the GH-1 family of glucosidases (Fig. 6b), while AsS6DGD is a member of the GH-3 family (Fig. 6b). Thus, CiS6DGD and AsS6DGD represent a case of convergently evolved enzymes (Weng, 2014; Weng & Noel, 2013). CiDGD and CiS6DGD are closely related, suggesting that these enzymes evolved through tandem-gene duplications and sub-functionalization. The broad specificity glucosidases CiDGD and AsDGD1,2 both belong to the GH-1 family of glucosidases, though phylogenetic analysis suggests that these enzymes also evolved independently (parallel evolution). Finally, the medium chain alcohol dehydrogenases DRs share the same protein fold and have relatively high sequence identity, but phylogenetic analysis also indicates that these enzymes evolved independently through means of parallel evolution (Supplementary Fig. 24).

## Discussion

Here we report the discovery of the protoemetine pathway in two distantly related plants, *C. ipecacuanha* and *A. salviifolium.* We show that *C. ipecacuanha* uses the monoterpenoid precursor secologanin (**1)**, while *A. salviifolium* uses secologanic acid (**2)**. This is consistent with monoterpenoid indole alkaloid biosynthesis in the Gentianales and Cornales clades, where tryptamine is condensed with secologanin or secologanic acid, respectively, to generate strictosidine or strictosidinic acid (Miller & Schuler, 2022; Sadre *et al*., 2016).

Strictosidine (3*S*) is stereo-selectively synthesized from secologanin and tryptamine by a well-characterized vacuolar localized Pictet-Spenglerase enzyme (strictosidine synthase) (Guirimand *et al*., 2010). The 3*R* isomer of strictosidine (vincoside) is not present in these plants. In contrast, all ipecac alkaloid producing plants contain products derived from both 1*S* and 1*R* stereoisomers of the initial Pictet-Spengler product (DAI **4b** (1*R*), DAII **4a** (1*S*) in *C. ipecacuanha* and DAIA **5b** (1*R*), DAIIA **5a** (1*S*) in *A. salviifolium*). Therefore, these plants must generate both 1-*R* and 1-*S* Pictet-Spengler products. We observed that non-stereoselective formation of DAI/I/(A) occurs when (i) dopamine and secologanin/secologanic acid are incubated in aqueous buffer at a pH value consistent with the environment of the vacuole, (ii) dopamine and secologanin/secologanic acid are infiltrated into *N. benthamiana* leaves, and (iii) dopamine is infiltrated into *C. roseus* flowers. The known plant Pictet-Spenglerases strictosidine synthase and norcoclaurine synthase are localized in the vacuole (Lee & Facchini, 2010; McKnight *et al*, 1991), though since these enzymes have optimal catalytic efficiency at neutral pH, the acidic environment of the vacuole is not required for the enzymatic reaction (Maresh *et al*, 2008; Samanani & Facchini, 2002). However, for a non-enzymatic reaction, the slightly acidic environment of the vacuole could be crucial, even for the highly activated, electron-rich dopamine substrate. We further showed that co-expression of the *C. roseus* vacuolar strictosidine exporter CrNPF2.9 along with *C. ipecacuanha* protoemetine biosynthetic genes greatly enhanced the levels of the protoemetine product, and facilitated the formation of the protoemetine derived product from starting substrates secologanin and dopamine (Supplementary Fig. 20). Collectively these observations support the notion that DAI/I(A) formation occurs non-enzymatically and within the plant vacuole in these pathways.

The ipecac pathway is characterized by the presence of both *R* and *S* epimers of the early tetrahydroisoquinoline intermediate. Both *C. ipecacuanha* and *A. salviifolium* evolved nearly identical chemistry that converts the *S* epimer to protoemetine. In contrast, the *R* epimers are derivatized in a chemically different and species dependent manner through a simple and shorter “shunt” pathway. Although protoemetine biosynthesis represents the same chemical outcome, there are striking species-specific evolutionary strategies by which enzymes were recruited to the protoemetine pathways. In *C. ipecacuanha*, all involved OMT enzymes (CiDOMT1,2 and CiDPOMT) are closely related and likely arose through gene duplication and subsequent neofunctionalization. Conversely, in *A. salviifolium*, enzymes from different OMT classes appear to have been independently recruited to ipecac alkaloid biosynthesis (AsDOMT is a class I OMT; AsDPOMT is a class II OMT). Additionally, the AsDOMT enzymes have promiscuous substrate specificity, also *O*-methylating the *R* epimer. In contrast, CiDOMT enzymes are substrate specific. For derivatization of the *R*-epimer, *C. ipecacuanha* instead recruited an acetyl transferase (BAHD) to synthesize ipecoside.

Both *C. ipecacuanha* and *A. salviifolium* evolved glucosidases with broad substrate specificity (DGDs) as well as glucosidases that appear to be specific to the intermediate on pathway to protoemetine biosynthesis (S6DGDs). While DGDs from both species are GH-1 type glucosidases, CiS6DGD and AsS6DGD are GH-1 and GH-3 type glucosidases, respectively. Thus, in *C. ipecacuanha* CiDGD and CiS6DGD likely evolved through tandem gene duplication and subfuctionalization. However, in *A. salviifolium* AsS6DGD convergently evolved this function upon recruitment from the GH3 family, a group of glucosidases with a different fold not commonly associated with specialized metabolism (Arthan *et al*, 2006; Suthangkornkul *et al*, 2016) but rather with cell wall biosynthesis (Hrmova *et al*, 1996; Kim *et al*, 2000b; Sampedro *et al*, 2017; Takahashi *et al*, 2018). The specificity of CiS6DGD and AsS6DGD ensures that 6-*O*-Me-DAIIA rather than DAI/I(A) are deglycosylated by these dedicated protoemetine glucosidaes. The evolution of glucosidases and OMTs in *A. salviifolium* could thus be described by the “patchwork hypothesis” which describes independent recruitment of promiscuous enzymes to a pathway while in *C. ipecacuanha* it could be rather described by the “forward hypothesis” which describes that a pathway evolves from the first biosynthetic enzyme onwards through means of tandem gene duplication and neofunctionalization (reviewed by (Noda-Garcia *et al*, 2018; Scossa & Fernie, 2020). Within the Gentianales, ipecac alkaloid pathway evolution in *C. ipecacuanha* appears have evolved independently from the related monoterpene indole alkaloid pathway evolution. Although some of the biosynthetic steps of these two pathways have similar chemistry (*e.g.* corynantheal from *Cinchona spp.* (Nomura & Kutchan, 2010; Nomura *et al*., 2008; Trenti *et al*., 2021), the enzymes that catalyze the respective steps (DR and OMT) do not cluster together with known monoterpene indole alkaloid biosynthesis enzymes but instead form sister clades (Fig. 6a, Supplementary Fig. 24).

It is likely that the *R*-epimers ipecoside (*C. ipecacuanha*) and 6/7-*O*-Me-DAIA (*A. salviifolium*) accumulate in the vacuole, since this is the typical storage location in plants for glycosylated specialized metabolites (Shitan & Yazaki, 2020). These products would therefore be separated from the nuclear localized glucosidases (DGDs) that act upon these substrates. Deglycosylation of ipecoside and 6/7-*O*-Me-DAIA leads to a dialdehyde moiety that is highly reactive (Guirimand *et al*., 2010; Kim *et al*, 2000a; Konno *et al*, 1999). The *R*-epimer-derived ipecoside and 6/7-*O*-Me-DAIA could be released from the vacuole upon tissue damage, only then coming into contact with the nuclear localized glucosidases to generate a reactive– and therefore toxic– defense molecule. Despite the species-specific chemical derivatization– *O*-methylation vs. acetylation– the *R*-epimers may serve similar ecological functions as defensive agents. Mechanisms in which the enzyme is spatially separated from its substrates to avoid the constant accumulation of potentially toxic compounds were first described for glucosinolates (reviewed by (Blazevic *et al*, 2020; Kissen *et al*, 2008). Instances in which a glucosidase is located in the nucleus while the substrate is stored in the vacuole have also been described for monoterpenoid indole alkaloids, saponins and for secoiridoids (Guirimand *et al*., 2010; Koudounas *et al*, 2017; Lacchini *et al*, 2023). Specialized metabolites stored in inactive form that are activated upon tissue damage are recently also being referred to as phytoavengins (Kliebenstein & Kvitko, 2023). Overall, this comparative pathway elucidation of ipecac alkaloid biosynthesis highlights the diversity in evolutionary strategies to evolve chemically complex molecule and provides a foundation for metabolic engineering of these biologically active molecules.

## Acknowledgements

We thank the Ghent University Botanical Garden and Chantal Dugardin for providing *A. salviifolium* living specimens and Néstor J. Hernández Lozada and Mohamed O. Kamileen for transport of these specimens; the gardeners Eva Rothe and Elke Goschala for growing and maintaining *A. salviifolium* and *C. ipecacuanha* plants as well as Franz Kaltofen for growing *N. benthamiana* plants, and Hendrik Tilger for installing tools on the in-house Galaxy server. We thank Dagny Grzech for providing pUPD constructs and CrNPF2.9 construct, Prashant Sonawane for providing the modified 3Ω1 vector and Mohamed O. Kamileen for providing *A. tumefaciens* strains harboring mcherry markers. We thank Matilde Florean for advice on phylogenetic analyses. We are grateful to the Max Planck Society for funding and the European Research Council (788301) and the Leibniz Prize, Deutsche Forschungsgemeinschaft (DFG, German Research Foundation) – 505457618 awarded to Sarah E. O’Connor.

## Author contributions

M.C. designed all experimental work, performed all pathway discovery experiments (generation and analysis of RNA-seq and metabolite data, screening and reconstitution in *N. benthamiana*), analyzed all data except NMR, and supervised the work of C.M., O.D., H.L. and B.C. C.M. performed subcellular localization experiments assisted by V.G. C.M. and M.C generated phylogenetic trees. O.D. assisted with enzyme screening and experiments on spontaneous coupling. R.A. synthesized protoemetine. B.C. purified protoemetine from *A. salviifolium*. H.L. purified DAI and DAII. B.H. assisted with secologanic acid synthesis. Y.N. performed NMR and analyzed NMR data. D.S., S.H. and M.K. developed chromatography and mass spectrometry methods. J.W. assisted with cloning. A.A.L. provided *C. ipecacuanha* living specimens and assisted with growth. K.W. cultured and regenerated *C. ipecacuanha in vitro*. M.C. and S.E.O. designed the study and wrote the manuscript.

