## Supplementary Information for "Independent evolution of ipecac alkaloid biosynthesis"

**Supplementary Dataset 1: separate excel file**

**RNA-seq data is accessible under *TBA*.**

#### Material and Methods

##### Plant material and sampling

*C. ipecacuanha* plantlets were grown *in vitro* on propagation medium (1x Murashige and Skoog medium including vitamins (MS incl. vit.), 3 % sucrose, 3 mg/L 6-Benzylaminopurine, 10 µg/L 1-Naphthaleneacetic acid, 8 g/L agar, pH 5.7). Upon arrival plantlets were transferred to root induction medium (0.75 x MS incl. vit., 3 % sucrose, 0.5 mg/L 1-Naphthaleneacetic acid, 8 g/L agar, pH 5.7). After approximately six weeks, roots formed and the regenerated plants were transferred to soil. For tissue specific metabolite profiling and RNA-seq, plants were harvested four months after transfer to soil. *A. salviifolium* plants were received as cuttings from the Botanical Garden of Ghent University, Belgium, and rooted on rockwool. After six weeks, rooted plants were transferred to soil. For tissue specific profiling, plants were harvested 14 months after transfer to soil. Both species were grown in a temperature and light controlled greenhouse with the following conditions: 12/12 hours light/dark 28-30°C/24-26°C, 70-80% humidity. For tissue specific analyses, three plants from each species were macro-dissected as shown in Supplemental Fig. 1. Tissues were immediately flash frozen into liquid nitrogen and ground in liquid nitrogen using an IKA A11 basic analytical mill or mortar and pestle. The frozen finely ground powder was kept at –80°C until further processing.

##### Metabolite extraction for tissue specific metabolite profiling

Finely ground material was extracted with 100 % MeOH supplemented with 10 mg/L caffeine as internal standard. The volume of MeOH was normalized to fresh tissue weight (1:10 = mg:µl). Samples were vortexed for 1 min and sonicated for 5 min at room temperature. After 1 hr of incubation at room temperature, samples were centrifuged for 15 min at 18000 *g* and supernatants were filtered through Fisher polytetrafluoroethylene (PTFE) syringe filters (0.22 µm). Filtered extracts were diluted in 80% MeOH containing 0.1% formic acid and 1:10 and 1:50 dilutions were analyzed by untargeted UPLC-MS. *C. ipecacuanha* samples were analyzed using UPLC-MS method 1 and *A. salviifolium* samples were with UPLC-MS method 2. For comparable traces shown in main text Fig. 2, representative samples of each plant were analyzed in parallel with UPLC-MS method 3.

##### UPLC-MS/MS methods

Method 1 was used for *C. ipecacuanha* tissue specific metabolite profiling. An Elute LC system (Bruker Daltonics, Massachusetts, USA) was coupled to an Impact II high resolution Quadrupole Time-Of-Flight (Q-TOF) mass spectrometer (Bruker Daltonics, Massachusetts, USA). An ACQUITY UPLC BEH C18 (2.1 x 50 mm, 1.7 µm; 130 Å) column (Waters, Ireland) was set at 40°C and 0.6 ml/min flow rate and 2 µl of samples were injected. The mobile phase was A:B where A was water with 0.1% formic acid and B acetonitrile with 0.1% formic acid. The gradient was as follows: 5% B at 1 min to 8% B at 3 min, to 13% B at 5 min and to 30% at 8 min. Then the column was flushed at 100% B until 9.8 min and re-equilibrated to 5% B until 12 min. Method 2 was used for *A. salviifolium* tissue specific profiling and was identical to Method 1 with the exception of the column temperature, which was set to 35°C. Method 3, used for all other experiments was identical to Method 2 but utilized an UltiMate 3000 Ultra-High Performance Liquid Chromatography (UHPLC) system (Thermo Fisher Scientific, Massachusetts, USA). For all methods, ionization was done via pneumatic assisted electrospray ionization (ESI+) with 4500 V (Method 1 and 2) or 3500 V (Method 3) capillary voltage and 500 V end plate offset; a nebulizer pressure of 2.5 bar, with nitrogen at 250°C and a flow of 11 L/min as the drying gas. Acquisition was done at 12 Hz following a mass range from 80 – 1000 *m/z* with data dependent MS/MS and an active exclusion window of 0.2 min, a reconsideration threshold of 1.8-fold change. Fragmentation was triggered on an absolute threshold of 400 and limited to a total cycle time range of 0.5 s. For collision energy, the stepping option model (from 20 to 50 eV) was used. At the start of each run the *m/z* values were re-calibrated using the expected cluster ion *m/z* values of a direct source infusion of sodium formate-isopropanol solution. To prevent the contamination of the MS by injection peaks and salt, the first minute of each run was run isocratically at 5% B and redirected to waste.

**MS data analysis**

Data was converted to mzml or mzxml format and imported to MZmine (versions 3-4) (Schmid *et al*, 2023). Extracted ion chromatogram (EIC) traces and $MS^2$ data of compounds of interest was exported from MZmine. Peak areas were calculated using the MZmine Processing Wizard and exported. Peak areas of compounds of interest were normalised to the internal standard caffeine and converted to intensity per second. Further data analysis and construction of graphs was done in GraphPad Prism version 10 for Mac OS X. For tentative identification of molecular structures as well as analysis of $MS^2$ fragmentation pattern SIRIUS was used (Bocker & Duhrkop, 2016; Duhrkop *et al*, 2019; Duhrkop *et al*, 2015).

**Non-enzymatic Pictet-Spengler reaction in *N. benthamiana* and *C. roseus***

For coupling in *N. benthamiana*, 1 mM of dopamine hydrochloride **3** and 1 mM of secologanin **1** or secologanic acid **2**, respectively, were infiltrated into leaves of 4-week-old plants (grown as described below). A leaf sample was taken immediately after infiltration for a control and snap frozen. For coupling in *C. roseus*, petals of freshly opened flowers of 4-month-old plants (grown in a controlled growth chamber at 16/8 hrs light/dark, 23°C, 40-50% humidity) were infiltrated with 1 mM of dopamine hydrochloride **3**. Samples were harvested after 24 hours. Control samples were taken after 24 hours from flower petals infiltrated with water. Metabolite extraction and analysis was performed as described below for *N. benthamiana*.

**RNA extraction and sequencing**

Total RNA was extracted using the RNeasy Plant Mini Kit (Qiagen) from replicate 1 of the same tissue material that was used for metabolite profiling according to the manufacturer's instructions, including on column DNAse digest. RNA concentrations and purity were determined with a Nanophotometer N60 (Implen). Samples of sufficient concentration and purity were shipped to Novogene where mRNA library preparation and sequencing were performed according to the company's standard protocol for mRNA sequencing. RNA integrity and quantitation were assessed using the RNA Nano 6000 Assay Kit of the Bioanalyzer 2100 system (Agilent Technologies, CA, USA). All samples were above the required minimum RIN value. Sequencing was performed on an Illumina NovaSeq 6000 PE150 platform with a data output target of 9 G of raw data. Adapter cleaved raw data was provided as fastq files.

**Transcriptome analysis and candidate selection**

All data processing was performed in-house. Reads were quality-checked with FastQC and trimmed using Trimmomatic (Bolger *et al*, 2014). Transcriptomes were assembled by rnaSPAdes using the trimmed reads of all tissues combined as an input (Bushmanova *et al*, 2019) with default settings, except k-mer size was adjusted to 55, 77, 99 to discriminate isoforms with high sequence identity. The resulting assemblies were assessed with Busco for the Eudicots lineage and were 95.6% complete for *C. ipecacuanha* and 97.2 % for *A. salviifolium* (Manni *et al*, 2021). All above tools (FastQC,

Trimmomatic, rnaSPAdes, BUSCO) were run on an in-house Galaxy server (Afgan *et al*, 2018). Functional annotation of transcripts was performed on OmicsBox (Biobam) using the SwissProt 2021 database (blasting parameters: E-Value, 1.0E-3; number of Blast Hits, 10; word size, 6; low complexity filter, on; number of threads, 40; HSP length cutoff, 33) and additionally with eggNOG-mapper (Gotz *et al*, 2008; Huerta-Cepas *et al*, 2016). Reads were mapped to transcriptomes using CLC Genomics workbench 21 (Qiagen) with these parameters: mismatch cost, 2; insertion cost, 3; deletion cost, 3; length fraction, 0.8; similarity fraction 0.85; auto-detect paired distances, on; maximum number of hits for a read, 10. TMM normalized CPM values were used for downstream analyses.

To identify known *C. ipecacuanha* pathway genes, the published sequences (NCBI AB455576, AB527082, AB527083, AB527084, AB576187) were blasted (blastn) against our transcriptome (Cheong *et al*, 2011; Nomura & Kutchan, 2010; Nomura *et al*, 2008). The level of pairwise sequence identity at the nucleotide level between the published sequences and the amplified sequences was: 96.8% for *CiDGD* (previously called *IpeGlu1*), 97.2 % for *CiDOMT1* (previously called *IpeOMT1*), 98.7 % for *CiDOMT2* (previously *IpeOMT2*) and 98.5 % for *CiDPOMT* (previously *IpeOMT3)*. The closest homologous sequence to another published OMT (Cheong *et al.*, 2011) was also *CiDOMT2* (98.8 % pairwise identity). These observed small differences in sequence identity between published and newly cloned sequences can be attributed to small nucleotide polymorphisms (SNPs) of different plant sources. The expression profile of *CiOMT1* was used as a bait for Pearson correlation (Supplementary Figure8).

To identify candidates for the missing esterase and reductase, *C. pubescens CpDCE* (MW456556.1) and *CpDCS* (MW456554.1) (Trenti *et al*, 2021) were blasted (tblastn) against the transcriptome which resulted in 9 contigs with 38-44 % identity and 15 contigs with 55.4-62% identity, respectively. The Blast hits were then filtered for high correlation with *CiDOMT1* tissue specific expression profile (Pearson correlation > 0.85) and high absolute expression level in young leaf and rhizome (> 50 CPM). Further mining the list of co-expressed and highly expressed contigs for relevant functional annotations revealed *Ci6SDGD* and *CiIpS*. For *A. salviifolium* no pathway genes were previously published and blasting *CiDOMTs* against the transcriptome did not yield any orthologs with high identity. We therefore identified a *TyrDC* ortholog whose expression pattern matched the tissue specific metabolite profiling and used it as bait for co-expression analysis. Among the highly co-expressed candidates (Pearson >0.75, CPM in leaf buds and/or roots >50) we selected those with functional annotations consistent with O-methyltransferases, dehydrogenases and glycosyl hydrolases for screening. After positive screening results for *AsDOMT*s, three additional highly homologous sequences with higher root specific expression (*AsDOMT2*, *6*, *7*) were included in pathway reconstitution experiments.

**Gene cloning**

cDNA was prepared from total RNA of *A. salviifolium* leaf buds and roots and *C. ipecacuanha* young leaves and rhizome (extracted as described above) using the RevertAid First Strand cDNA Synthesis Kit (Thermo Scientific) according to manufacturer's instructions. *AsDOMT5* and *AsDPOMT2* could not be amplified and were therefore obtained as synthetic sequences from Twist Biosciences. *CDS* sequences were amplified with the Q5 High-Fidelity 2X Master Mix (New England Biolabs) using cDNA or synthetic genes as templates and gene specific primers containing overhangs for In-Phusion cloning (Supplementary table 1). Amplified sequences were gel-purified using the Zymoclean Gel DNA Recovery Kit (Zymo Research) and cloned using the 5x In-Fusion Snap Assembly Master Mix (TaKaRa Bio). For expression in *N. benthamiana*, coding regions were inserted into a modified 3Ω1 vector (contains *UBQ10* promoter and terminator from *Solanum lycopersicum* (Cardenas et al, 2019)) previously digested with BsaI-HF v2 (New England Biolabs). For expression in *Escherichia coli CDS'* were cloned into pOPINF (Berrow *et al*, 2007) previously digested with KpnI-HF and HindIII-HF (New England Biolabs). Heat shock competent *E. coli* TOP10 were transformed and grown over night in a 37°C incubator on LB Agar plates containing the respective antibiotics. Plasmids were isolated from over- night cultures of single colonies using the Wizard Plus SV Minipreps DNA Purification System kit (Promega). Inserted sequences were confirmed by Sanger sequencing.

*DGD* sequences for subcellular localization were amplified and purified as described above but using the previously cloned 3Ω1 constructs as PCR templates (see above) and gene-specific primers with overhangs compatible with Golden Gate cloning for N- or C-terminal eYFP fusion proteins (Supplementary Table 1). Constructs (level 1) were assembled using *BsaI* (New England Biolabs) and T4 DNA ligase (New England Biolabs), the pDGB3_ɑ1 vector (Sarrion-Perdigones *et al*, 2011), pUPD_pSlUbq10 and pUPD_TeSlUbq10 (Cardenas *et al.*, 2019; Grzech *et al*, 2023) and the gel purified PCR products. *As6SDGD* contained a *BsaI* restriction site which was removed by introducing a silent mutation in an overlapping PCR prior to assembly. Assembling was done by 50 cycles of 5 min at 37°C followed by 5 min at 16°C and stopped by incubation at 65°C for 10 min. Additionally, *CrNPF2.9* was cloned through the same Golden Gate cloning procedure as above into pDGB3_ɑ1 (without eYFP tag) whereas the silencing repressor gene *p19* was obtained at pUPD_p19 (addgene GB0038) and then cloned as above into pDGB3_ɑ1.

##### *A. tumefaciens* mediated transient expression in *N. benthamiana*

In all cases the *A. tumefaciens* GV3101 strain was used and cultured at 28°C in YEB medium containing rifampicin and gentamycin and the appropriate antibiotic for plasmid selection. *N. benthamiana* plants used for infiltration were 3-4 weeks old (grown in a greenhouse with 16h/8h light/dark and 23-26°C /16-22°C, 40-70% humidity). Cells were transformed through electroporation, recovered in YEB without antibiotics and incubated for two days on YEB plates containing antibiotics. Single colonies were confirmed by colony PCR, and grown in liquid YEB for 24 hours. From these cultures glycerol stocks were prepared and stored at –80°C. For agroinfiltration, cells were grown on YEB plates similar to a previously described method with modifications (Zhang *et al*, 2020): Cells from glycerol stocks were spread on YEB plates containing antibiotics and 100 µM acetosyringone and grown for 24 hours until a visible layer of bacteria appeared. The bacteria were transferred to 1-2 ml of infiltration medium (10 mM MES, 10 mM $MgCl_2$, 100 µM acetosyringone, pH 5.7), gently resuspended and the $OD_{600}$ measured in 1:10 dilutions using an Implen OD600 DiluPhotometer.

For all pathway reconstitution experiments, strains were mixed and diluted in infiltration buffer to $OD_{600}$=0.1 per strain. A strain harboring a construct with the *p19* gene was co-infiltrated in all cases. The culture mixtures were infiltrated into *N. benthamiana* leaves and plants were kept for 16 hours in the dark and subsequently grown under grow lights (16h/8h light/dark). Replicates are from three individual plants. Indicated substrates were infiltrated as 500 µM aqueous solutions of epimeric mixtures (see below) three days post infiltration. Leaf material was harvested 24 hours after substrate infiltration by flash freezing in tubes containing metal beads. In the case of infiltration of uncoupled secologanin and dopamine substrates, leaf material was harvested 48 hours after substrate infiltration to allow time for coupling.

For analysis of *N. benthamiana* leaves by confocal laser scanning microscopy, *A. tumefaciens* culturing and infiltration was performed as above but at $OD_{600}$=0.3 per strain. To confirm subcellular localization, each strain harboring an eYFP fusion construct was co-infiltrated with a strain harboring a construct with free mcherry or with NLS-mcherry as fluorescent markers for cytosol or nuclear localization, respectively (Ivanov & Harrison, 2014), along with a strain for expression of *p19*. Leaf tissue was analyzed 2-3 days post infiltration.

**Leaf disk assays**

To test glucosidase activity towards different substrates, a leaf disk assay was employed as previously described with modifications (Hong *et al*, 2022; Kamileen *et al*, 2022; Kamileen *et al*, 2024). Growth and infiltration of *A. tumefaciens* was done as described above and replicates are from three individual plants. Three days post agroinfiltration, 1 cm leaf disks were cut using a leaf puncher and incubated with 200 µl of substrate solution in 50 mM HEPES buffer pH 7.5 in 48-well plates. Substrate solutions were prepared as Master mixes so that concentrations were identical in each well. Substrate concentrations were 400 µM total, corresponding to 200 µM per epimer for chemically produced epimeric substrate mixtures DA(I)I, DAI(I)A, 7-*O*-Me-DAI(I), 7-*O*-Me-DAI(I)A. Commercially available ipecoside was used at 200 µM. The reactions of enzymatically produced substrates (see below) were monitored by UPLC-MS and concentrations were estimated at 250 µM. Plates were sealed with parafilm to avoid evaporation and incubated for 24 hours under grow lights 16/8 light/dark.

**Metabolite extraction from *N. benthamiana***

Leaf material was ground using two 4-mm metal beads and a TissueLyser (Qiagen) with pre-cooled adapters, and extracted with 100 % MeOH containing 0.1 % formic acid and 2 mg/L caffeine as internal standard. For substrate infiltrated leaves, 150 µl per 100 mg leaf material was used. In the case of leaf disk assays, 50 µl was used per single leaf disk. Samples were sonicated for 10 min, incubated on a rotator for 20 min and centrifuged at 18000 *g* for 15 min. The supernatants were mixed 1:2 with MQ H2O to improve shape of early eluting peaks and filtered through a 0.45 µm low binding hydrophilic PTFE filter plate (MultiScreen Solvinert 96, Merck-Millipore) into a 96-well Microtiter Plate (SureSTART WebSeal, Thermo Scientific) according to manufacturer's instructions. Plates were sealed with Rapid Slit Seal (BioChromato) and immediately analyzed with UPLC-MS method 3.

**Confocal laser scanning microscopy**

For confocal laser scanning microscopy, the leaf discs were put on a glass slide, mounted with water, and covered with a coverslip. Fluorescence was observed and imaged with a W Plan-Apochromat 40x/1.0 DIC M27 water objective on a cLSM 880 (both Zeiss, Oberkochen, Germany) equipped with two lasers for excitation of the two different fluorophores. mCherry was excited at 543 nm with a helium-neon laser and emission was filtered between 600 and 651 nm. eYFP was excited at 514 nm with an argon laser and emission was filtered between 525 and 561 nm. The micrographs were taken sequentially in two tracks for each image. In the first track, mCherry emissions were captured as well as the transmitted light image. In the second track, eYFP emission was captured. ZEN black 2.1 V.14.0.18.201 was used as software (Zeiss, Oberkochen, Germany). Images were adjusted and processed with ImageJ software (Schindelin *et al*, 2012).

**Recombinant protein production and purification**

Expression and purification of AsDOMT3, CiDOMT1 and CiDE was performed as previously described with modifications (Stavrinides *et al*, 2016). Briefly, *E. coli* SoluBL2 (DE3) were transformed by heat shock with pOPINF constructs. Pre-cultures were inoculated from single colonies, grown over night at 37°C and used to inoculate 100 ml 2 x YT medium. Cultures were grown at 37°C until $OD_{600}$ 0.5-0.6 was reached, cooled to room temperature and induced with 0.2 mM Isopropyl β- D-1-thiogalactopyranoside (IPTG). After induction the cultures were grown at 18°C over-night and harvested by centrifugation the next day. Cell pellets were lysed using B-PER™ Complete Bacterial Protein Extraction Reagent (Thermo Scientific) supplemented with EDTA and subsequently centrifuged according to manufacturer's instructions. Supernatants were incubated with gentle shaking in falcon tubes with 250 µl Ni-NTA Agarose (Qiagen) for one hour at 4°C to allow binding of His-tagged proteins. Slurry was pelleted gently by centrifugation at 1000 g for 30 s. The supernatant was removed and the slurry was washed three times with ice cold wash buffer (50 mM Tris-HCL pH8, 50 mM glycine, 5% glycerol, 500 mM NaCl, 20 mM imidazole) by inversion, centrifugation and removal of supernatant. Proteins were eluted using elution buffer (as wash buffer but containing 500 mM imidazole). Elution fractions were concentrated and buffer was exchanged to storage buffer (20 mM HEPES, 150 mM NaCl, pH 7.5) using Amicon Ultra Centrifugal Filters (Millipore) with the appropriate exclusion size according to manufacturer's instructions. Purity was assessed through SDS-PAGE and concentration was determined using the extinction coefficient and measuring the absorbance at 280 nm. Proteins were flash frozen in small aliquots in liquid nitrogen and stored at –80°C for storage.

**Phylogenetic analyses**

Sequences were obtained through BLAST searches against the publicly available NCBI and 1KP (One Thousand Plant Transcriptomes, 2019) databases or from *C. ipecacuanha* and *A. salviifolium* transcriptomes generated in the frame of this study. Sequences and accession numbers are provided in Supplementary dataset 1. Full-length amino acid sequences were aligned with webPRANK (https://www.ebi.ac.uk/goldman-srv/webprank/) (Löytynoja & Goldman, 2010) and Maximum Likelihood phylogenetic trees were build using the IQ-TREE web server (http://iqtree.cibiv.univie.ac.at/) (Trifinopoulos *et al*, 2016) (automatic substitution model; bootstrap value 1000). iTOL was used to visualize and graphically edit trees (https://itol.embl.de/) (Letunic & Bork, 2024).

**Protein models**

Protein models were predicted with AlphaFold3 through the AlphaFold Server (https://alphafoldserver.com/) (Abramson *et al*, 2024). Models were visualized using ChimeraX version 1.8 for Mac (Meng *et al*, 2023).

**Commercially available chemicals and standards**

Secologanin (50741), dopamine hydrochloride (H3502) and 4-*O*-Me-dopamine hydrochloride (H3132), cephaeline dihydrochloride (PHL85887, phyproof Reference Substance), emetine dihydrochloride (PHL89489, phyproof Reference Substance) and ipecoside (TA9H93CFC242, TargetMol) were purchased from Sigma.

**Chemically produced standards and substrates**

Secologanic acid was produced through alkaline hydrolysis of secologanin by incubating secologanin with 0.1 M NaOH (40 µl per 1 mg secologanin) for five hours similar to a previously described protocol (Chapelle, 1976). The solution was neutralized and completeness of the reaction was confirmed through LC-MS analysis. Secologanic acid was stored in solution at –25°C.

Epimeric mixtures of DAI(I), DAI(I)A, 7-*O*-Me-DA(I)I and 7-*O*-Me-DAI(I)A were obtained using a previously described protocol with modifications (Nomura *et al.*, 2008): 10 mM secologanin or secologanic acid were incubated together with 10 mM dopamine hydrochloride or 4-*O*-Me-dopamine hydrochloride, respectively, in 0.1 M citrate/ 0.2 M phosphate buffer pH 5.5 for 24 hours at room temperature. This was sufficient for coupling as confirmed by UPLC-MS analysis. 6-*O*-Me-DA(I)A cannot be produced using this protocol and was produced enzymatically (see below).
Protoemetinol was produced from protoemetine (see below) by reduction with $NaBH_4$. An aliquot of 50 µg of protoemetine was incubated in MeOH with 0.5 g/L $NaBH_4$ for 30 minutes at room temperature. The reaction was quenched by adding an equal volume of acetone followed by a 15 minutes incubation at room temperature. The reaction was centrifuged at 18000 g for 15 minutes, diluted with MeOH and analyzed with UPLC-MS method 3 which indicated full conversion to a compound consistent with *m/z* $MS^2$ of protoemetinol (Supplementary Fig. 7).

**Purification of epimer pure DAII and DAI**

An epimeric mixture of DAII and DAI was produced as described above but with the following modifications: a 1.2 fold excess of dopamine hydrochloride was used, the reaction was bubbled with argon and then incubated for three days. Preparative isolation was performed using an Xbridge BEH C18 OBD Prep Column 130A, 5 µm (Waters) on an Agilent 1260 HPLC system (with G7161A Prep Bin pump, G9328A column organizer, G7165A detector) and an Agilent 1290 G7159B fraction collector. The mobile phase was A:B where A was water with 0.1 % formic acid and B acetonitrile with 0.1 % formic acid. The separation method was as follows: 5 % B from 1 min to 25 % B at 24 min, then the column was flushed at 100 % B until 29 min and re-equilibrated to 5 % B until 35 min. The flow rate was 8 mL $min^{-1}$, and detection was performed at $\lambda$ = 254 nm. Aliquots of fractions of each of the two main peaks were analyzed by UPLC-MS method 3 confirming that *m/z* and retention times corresponded to either of the two peaks typically observed in epimeric mixtures (Supplementary Fig. 5). Collected fractions of each peak were combined, dried using a freeze dryer (Labconco) and subjected to NMR.
Deacetylisoipecoside (DAII, **4a**) was unstable during 2D NMR measurements in MeOH-$d_3$, and was therefore measured in acidic MeOH-$d_3$ (MeOH-$d_3$ + 0.1% formic acid), to prevent decomposition during long-term measurements. Deacetylipecoside (DAI **4b**) was lactamized even in acidic MeOH-$d_3$. Thus, the structure of **4b** was confirmed as demethylalangiside after complete lactamization. This observation was consistent with previously reported fact that the lactamization was much faster in the 1*R* than the 1*S* (Beke et al, 2001).

**NMR analysis**

NMR measurements were carried out on a 500 MHz Bruker Avance III HD spectrometer (Bruker Biospin GmbH, Rheinstetten, Germany), equipped with a TCI cryoprobe using standard pulse sequences as implemented in Bruker Topspin ver. 3.6.1. (Bruker Biospin GmbH, Rheinstetten, Germany) or a 400 MHz Bruker Avance III HD spectrometer (Bruker Biospin GmbH, Rheinstetten, Germany). Chemical shifts were referenced to the residual solvent signals of $CDCl_3$ ($\delta_H$ 7.26/$\delta_C$ 77.16) or MeOH-$d_3$ ($\delta_H$ 3.31/$\delta_C$ 49.0), respectively, and coupling constants (*J*) are expressed in Hertz (Hz), in the following format; chemical shift value (multiplicity, coupling constant, integration). $^1$H NMR spectral data are described, using the following abbreviations; brs (broad singlet), s (singlet), d (doublet), t (triplet), q (quartet), dd (doublet of doublets), ddd (doublet of doublets of doublets), dt, (doublet of triplets), td (triplet of doublets), appd (apparent doublet), and m (multiplet).

Based on the structure determined from NMR analysis, a molecular model was created in GaussView ver.6 (Semichem Inc., Shawnee, Kansas, USA) and optimized using the semi-empirical method PM6 in Gaussian ver.16 (Gaussian Inc., Wallingford, Connecticut, USA). The optimized conformation from PM6 was used for ROESY analysis of protoemetine. For deacetylisoipecoside and demethylalangiside, the resulting structures were used for conformer variation with the GMMX processor of the Gaussian program package. Resulting structures were DFT-optimized with Gaussian ver.16 (B3LYP/6-31G(d), gas phase). The lowest energy conformer from the DFT calculations was used for the ROESY analysis.

**Enzymatically produced standards and substrates**

6-*O*-Me-DAIIA and 6-*O*-Me-DAIA were produced using recombinant CiDOMT1 and AsDOMT3, respectively. The reaction mix contained 50 mM HEPES pH 7.5, 1 mM total epimeric DAI(I)A (ca. 500 µM per epimer) as substrate, 1 mM of potential metal cofactors $MgCl_2$ and $MnCl_2$, 1 mM *S*-(5′-Adenosyl)-L-methionine (SAM) chloride dihydrochloride and 5 µM CiDOMT1 or 10 µM AsDOMT3. Reactions were incubated for 24 hours at 30°C and stopped by incubation at 98°C for 10 min to inactivate enzymes. Precipitates were pelleted by centrifugation at 18000*g* for 30 min. Supernatants were diluted 1:2 in H2O and used as substrates in leaf disk assays. Aliquots of the reactions were analyzed by UPLC-MS method 3 to observe expected products (Supplementary Fig. 5).

Confirmation of DAIIA and DAIA peak identity was done through deesterification of purified DAII and DAI, respectively, using recombinant CiDE. Reaction mixes contained 50 mM HEPES pH 7.5, 3 µM recombinant CiDE and 200 µM purified DAII or DAI, respectively. Reactions were incubated overnight at 30°C and stopped by the addition of 50 µl MeOH supplemented with 0.1% FA. Eventual precipitates were pelleted by centrifugation at 18000g for 30 min and supernatants were analyzed by UPLC-MS method 3 (Supplementary Fig. 5).

**Isolation of protoemetine from *A. salviifolium* and production of protoemetinol standard**

Protoemetine was isolated in several batches from a total of 6 g of freshly harvested leaf buds and young leaves. The material was flash frozen and ground with liquid nitrogen and 100% MeOH was added at a ratio of 200 µl per 100 mg. Samples were sonicated in an ultrasonic bath for 10 min, incubated on a rotator for 30 min and centrifuged at 18000 g for 15 min. The supernatants were filtered through 0.22 µm syringe filters, diluted 1:10 in 100% MeOH and subjected to semi-preparative HPLC. Semi-preparative isolation was done using an XBridge BEH C18 column of 100 mm x 4.6 mm, 2.5 µm (Waters) on an Agilent 1260 Infinity HPLC high-performance liquid chromatography system (with G1311B Quat pumps, G1315C diode array detectors, G1316A oven, G1329B autosampler, and G1364F fraction collector). The mobile phase was A:B, where A was 0.1% formic acid in water, and B was acetonitrile. The separation method was as follows: 8% B at 1 min to 13% B at 4 min, to 30% at 12 min. Then the column was flushed at 100% B until 15 min and re-equilibrated to 8% B until 18 min. The flow rate was 2.4 mL $min^{-1}$, and detection was performed at $\lambda$ = 250 nm. Aliquots of collected fractions were analyzed using UPLC-MS method 3 and fractions containing the putative protoemetine peak, based on *m/z* and typical aldehyde peak shape, were combined and freeze dried using a freeze dryer (Labconco). Large freeze-dried fractions were solubilized in small volumes of 100 % MeOH, combined (around 350 µg) and dried under a nitrogen line. NMR was conducted on a Bruker Avance III HD 700 MHz spectrometer (Bruker Biospin GmbH, Rheinstetten, Germany), equipped with a TCI cryoprobe using standard pulse sequences implemented in Bruker Topspin version 3.6.1 (Bruker Biospin GmbH, Rheinstetten, Germany). Chemical shifts were referenced to $CDCl_3$ residual solvent signals ($J_H$ 7.26/$J_C$ 77.16). NMR spectra of the sample were: $^1$H-NMR (700 MHz,$CDCl_3$): $\delta$ ppm: 0.93 (t, $J$ = 7.5 Hz, 3H), 1.13 (m, 1H), 1.29 (m, 1H), 1.48 (m, 1H), 1.58 (m, 1H), 1.95 (m, 1H), 2.08 (dd, $J$ = 11.3 Hz, 1H), 2.33 (m, 2H), 2.50 (ddd, $J$ = 11.9, 11.4, 4.5 Hz, 1H), 2.63 (m, 1H), 2.72 (dd, $J$ = 16.4, 3.3 Hz, 1H), 2.98 (m, 1H), 3.10 (m, 2H), 3.13 (m, 1H), 3.83 (s, 3H), 3.83 (s, 3H), 6.57 (s, 1H), 6.63 (s, 1H), 9.88 (bs, 1H); 13C-NMR (175 MHz, $CDCl_3$): $\delta$ ppm: 10.8, 23.6, 29.1, 35.7, 38.1, 41.3, 48.1, 52.3, 55.8, 55.8, 61.2, 62.6, 108.0, 111.4, 126.2, 129.4, 147.0, 147.3, 202.4. This spectrum is in agreement with a previously published NMR spectrum of protoemetine (Lin *et al*, 2011) and with that of chemically synthesized protoemetine in this study (Supplementary Fig. 24 ). Isolated and synthesized protoemetine were thus identical.

**Chemical synthesis of (–)-protoemetine**

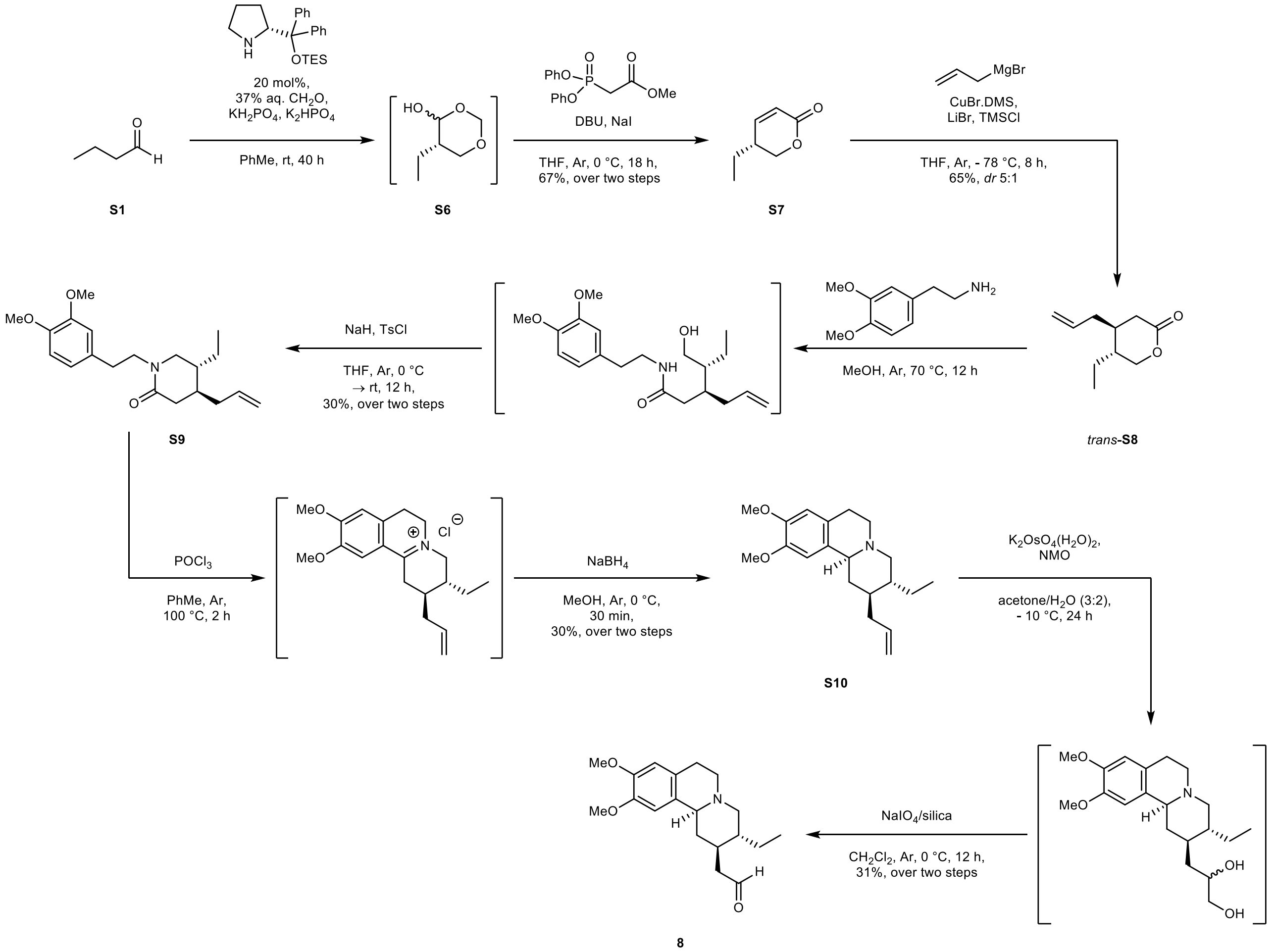

**Scheme 1** Total synthesis of (−)-protoemetine (**8**) (Xie *et al*, 2017)

#### General experimental note

All reagents were obtained from commercial sources and used without any further purification. Thin layer chromatography (TLC) was carried out on aluminum-backed silica gel 60 $F_{254}$ plates (Merck) that were visualized with Hanessian's stain or using $UV_{254\ nm}$ light detection. Wet flash column chromatography was performed using Merck Kieselgel 60 (particle size 0.040–0.063 mm, density 0.8 g/cm$^3$).

**(*R*)-2-{Diphenyl[(triethylsilyl)oxy]methyl }pyrrolidine (S2)**

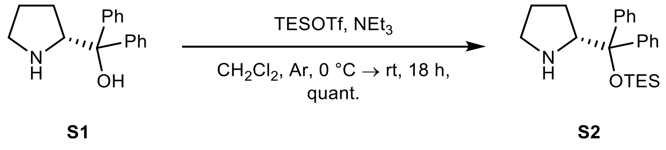

Briefly, following a modified procedure reported previously (Marigo *et al*, 2005), to a dry 50 mL round bottom flask under an Ar atmosphere, containing a mixture of (*R*)-diphenyl(pyrrolidin-2-yl)methanol (**S1**,1 g, 3.95 mmol) and $NEt_3$ (0.72 mL, 5.14 mmol) in anhydrous $CH_2Cl_2$ (26 mL) at 0 °C, was slowly added TESOTf (1.16 mL, 5.14 mmol). The reaction mass was allowed to warm to room temperature and stir for 18 h. To the resulting colorless solution was added $H_2O$ (30 mL). The mixture was extracted with $CH_2Cl_2$ (10 mL x 3) and the combined organic layers were dried over anhydrous $Na_2SO_4$. Following concentration under reduced pressure, the colorless residue obtained was purified by gradient flash column chromatography (0–10% MeOH in $CH_2Cl_2$) to afford title compound **S2** as a colorless oil (1.450 g, quant.): $^1$H NMR ($CDCl_3$, 400 MHz) $\delta$ 7.48–7.45 (m, 2H), 7.37–7.34 (m, 2H), 7.30–7.22 (m, 6H, overlapping with residual $CHCl_3$), 4.05 (t, $J$ = 7.4 Hz, 1H), 2.87–2.81 (m, 1H), 2.71–2.65 (m, 1H), 1.95 (brs, 1H), 1.64–1.52 (m, 3H), 1.29–1.22 (m, 1H), 0.85 (t, $J$ = 7.9 Hz, 9H), 0.35 (q, $J$ = 7.9 Hz, 6H). Spectral and physical data agreed with those reported previously (Hayashi *et al*, 2005).

**Methyl 2-(diphenoxyphosphoryl)acetate (S4)**

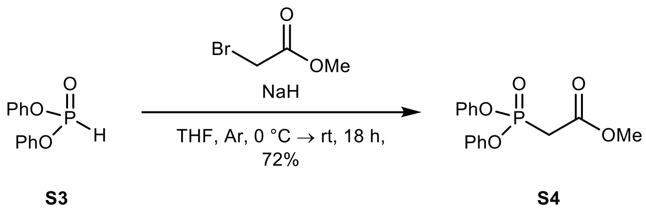

Following a previously reported procedure (Akwaboah *et al*, 2017), to a dry 250 mL three-neck round bottom flask equipped with a dropping funnel under an Ar atmosphere was added anhydrous THF (31 mL) and NaH (60% dispersion in mineral oil, 3.2 g, 80 mmol). Upon cooling to 0 °C, a solution of diphenyl phosphite (**S3**, 15.36 mL, 80 mmol) in anhydrous THF (10.3 mL) was slowly added to the mixture. After stirring at 0 °C for 1 h, a solution of methyl bromoacetate (7.55 mL, 80 mmol) in anhydrous THF (20 mL) was then added dropwise over 2 h to the reaction mass that was then subsequently allowed to warm to room temperature and stir for a further 18 h. The resulting colorless suspension was cooled to 0 °C and quenched with sat. aq. $NH_4Cl$ (40 mL) and $H_2O$ (50 mL). The mixture was extracted with $Et_2O$ (100 mL x 3) and the combined organic layers dried over $Na_2SO_4$. Concentration under reduced pressure afforded an opaque residue that was further purified by gradient flash column chromatography (0–50% EtOAc in hexane) to yield title compound **S4** as a colorless oil (17.736 g, 72%): $^1$H NMR ($CDCl_3$, 400 MHz) $\delta$ 7.36–7.30 (m, 4H), 7.24–7.16 (m, 6H), 3.77 (s, 3H), 3.28 (d, $J_{PH}$ = 21.6 Hz, 2H). Spectral and physical data agreed with those reported previously (Bodenschatz *et al*, 2022).

**(*R*)-5-Ethyl-5,6-dihydro-2*H*-pyran-2-one (S7)**

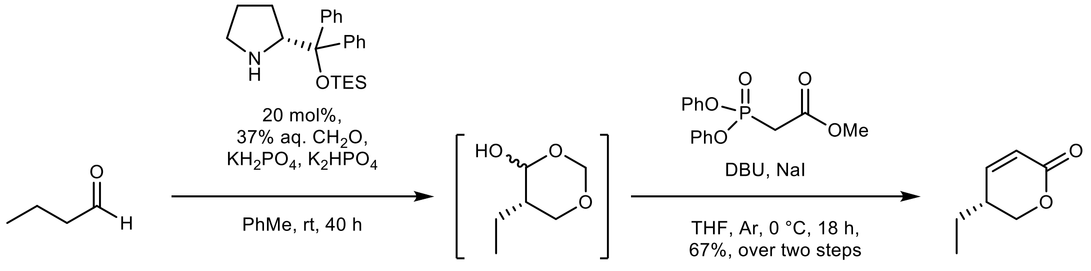

**S5** **S6** **S7**

Following a previously reported procedure (Xie *et al.*, 2017), to a dry 100 mL round bottom flask under an Ar atmosphere containing the silyl ether **S2** shown above (1.452 g, 3.95 mmol), $KH_2PO_4$ (618 mg, 4.54 mmol), $K_2HPO_4$ (1.099 g, 6.31 mmol), in toluene (9.9 mL), was added 37% aq. $CH_2O$ (5.89 mL, 72.57 mmol). After stirring at room temperature for 10 min, the mixture was cooled to 0 °C and charged with butyraldehyde (**S5**, 1.78 mL, 19.72 mmol). The reaction mixture was allowed to warm to room temperature and then rapidly stir for a further 40 h. The resulting biphasic mixture was extracted with toluene (5 mL x 6) and the combined organic layers were washed with brine (10 mL) and dried over anhydrous $Na_2SO_4$. The organic phase was carefully dried *in vacuo* (40 mbar at 30 °C) to afford the hemiacetal **S6** which was then immediately carried forward without any further purification to the next reaction step. To a dry three-neck round bottom flask fitted with a dropping funnel under an Ar atmosphere, containing anhydrous THF (143 mL), was added diphenyl phosphonate **S4** (17.514 g, 57.19 mmol) and NaI (9.458 g, 63.10 mmol). The resulting solution was cooled to 0 °C and then charged with DBU (7.03 mL, 47.13 mmol). After stirring at 0 °C for 30 min, to the reaction mass was then slowly added a solution of crude hemiacetal **S6** in anhydrous THF (33 mL). After stirring at 0 °C for 24 h, the colorless suspension was charged with 1 M aq. HCl (50 mL) and then further diluted with $H_2O$ (100 mL). The mixture was then extracted with EtOAc (100 mL x 3) and the combined organic layers were washed with brine (100 mL) and dried over anhydrous $Na_2SO_4$. Concentration under reduced pressure gave an amber residue that was further purified by gradient flash column chromatography (0–6.25% EtOAc in petroleum ether [b.p. 40–60 °C]) to afford pentenolide **S7** as a light yellow oil (1.673 g, 67% [over two steps]): $^1H$ NMR ($CDCl_3$, 400 MHz) $\delta$ 6.86 (dd, $J$ = 9.8, 3.8 Hz, 1H), 5.99 (dd, $J$ = 9.8, 1.8 Hz, 1H), 4.42 (dd, $J$ = 11.1, 5.0 Hz, 1H), 4.15 (dd, $J$ = 11.1, 7.4 Hz, 1H), 2.47–2.39 (m, 1H), 1.59–1.47 (m, 2H), 1.02 (t, $J$ = 7.5 Hz, 3H). Spectral and physical data agreed with those reported previously (Xie *et al.*, 2017).

**(4*S*,5*R*)-4-Allyl-5-ethyltetrahydro-2*H*-pyran-2-one (*trans*-S8)**

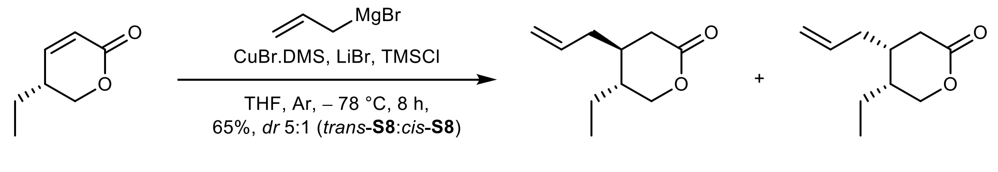

**S7** *trans*-**S8** *cis*-**S8**

Following a previously reported procedure (Xie *et al.*, 2017), to a dry 250 mL three-neck round bottom flask fitted with a dropping funnel under an Ar atmosphere, was added CuBr.DMS (5.452 g, 26.52 mmol), LiBr (2.303 g, 26.52 mmol), and anhydrous THF (88 mL). Upon cooling to – 78 °C, the resulting suspension was slowly charged with a 0.7 M solution of allylmagnesium bromide in THF (39.79 mL, 27.85 mmol) over 30 min and then stirred at – 78 °C for a further 10 min, followed by the addition of trimethylsilyl chloride (6.33 mL, 49.86 mL). After stirring for 10 min at – 78 °C, a solution of pentenolide **S7** (1.673 g, 13.26 mmol) in anhydrous THF (10 mL) was added dropwise to the reaction mixture. Upon stirring for 8 h at – 78 °C, the dark brown reaction mass was partitioned between a mixture of $CH_2Cl_2$ (100 mL) and $H_2O$ (100 mL) and then filtered through a short pad of Celite®. The solids were washed with $CH_2Cl_2$ (10 mL x 3). The combined filtrate was then separated and the resulting aqueous layer extracted with $CH_2Cl_2$ (100 mL x 3). The combined organic layers were washed with brine (200 mL) and dried over anhydrous $Na_2SO_4$. Concentration under reduced pressure afforded an amber oil that was subsequently purified by gradient flash column chromatography (0–6.25% EtOAc in petroleum ether [b.p. 40–60 °C]) to yield a mixture of *trans*-**S8** and *cis*-**S8** (*dr* 5:1, respectively) as a colorless oil (1.457 g, 65% [over two steps]): $^1$H NMR ($CDCl_3$, 400 MHz) *trans*-**S8*** $\delta$ 5.77–5.65 (m, 1H), 5.11–5.06 (m, 2H), 4.29 (dd, $J$ = 11.5, 4.4 Hz, 1H), 4.00 (dd, $J$ = 11.4, 7.5 Hz, 1H), 2.58 (dd, $J$ = 16.3, 6.8 Hz, 1H), 2.28 (overlapping dd, $J$ = 16.3, 7.7 Hz, 1H), 2.27–2.20 (overlapping m, 1H), 1.84–1.75 (m, 1H), 1.60–1.48 (m, 2H), 1.42–1.29 (m, 1H), 0.94 (t, $J$ = 7.4 Hz, 3H). Spectral and physical data agreed with those reported previously (Xie *et al.*, 2017). *Isolation of *trans* was achieved by iterative gradient flash column chromatography (0–6.25% EtOAc in petroleum ether [b.p. 40–60 °C]).

**(4*S*,5*R*)-4-Allyl-1-(3,4-dimethoxyphenethyl)-5-ethylpiperidin-2-one (S9)**

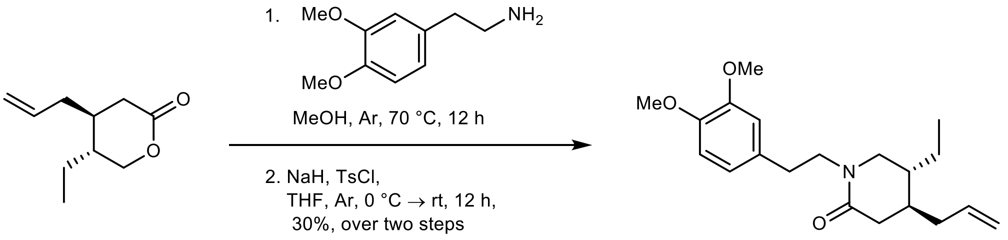

*trans*-**S8** **S9**

To a dry 15 mL pressure tube under an Ar atmosphere was added *trans*-**S8** (500 mg, 2.97 mmol) anhydrous MeOH (1.43 mL), and homoveratrylamine (0.75 mL, 4.46 mmol). The mixture was heated to 70 °C and allowed to stir for 12 h after which thin-layer chromatography (33% EtOAc in in petroleum ether [b.p. 40–60 °C]) indicated the complete consumption of starting material *trans*-**S8**. Concentration of the reaction mass under reduced pressure gave a residue that was then transferred to a dry 100 mL round bottom flask under an Ar atmosphere and dissolved in anhydrous THF (20 mL). Upon cooling to 0 °C, the solution was charged with 60% (w/w) NaH in mineral oil (549 mg, 13.72 mmol). After stirring for 30 min at 0 °C, a solution of TsCl (1.416 g, 7.43 mmol) in anhydrous THF (7.4 mL) was then slowly added to the resulting suspension and the reaction mass allowed to warm to room temperature. After stirring for 12 h, $H_2O$ (10 mL) was added. The mixture was extracted with EtOAc (10 mL x 5) and the combined organic layers were washed with brine (20 mL) and dried over anhydrous $Na_2SO_4$. Concentration *in vacuo* gave a residue that was further purified by gradient flash column chromatography (50% EtOAc in petroleum ether [b.p. 40–60 °C]) to yield lactam **S9** as a yellow oil (293 mg, 30% [over two steps from *trans*-**S8**]): $^1$H NMR ($CDCl_3$, 400 MHz) $\delta$ 6.81–6.76 (m, 3H), 5.75–5.65 (m, 1H), 5.07–4.99 (m, 2H), 3.87 (s, 3H), 3.85 (s, 3H), 3.64–3.65 (m, 1H), 3.52–3.45 (m, 1H), 3.13 (dd, $J$ = 12.3, 5.0 Hz, 1H), 2.88–2.82 (m, 3H), 2.47 (dd, $J$ = 17.5, 5.6 Hz, 1H), 2.22–2.16 (m, 1H), 2.10 (dd, $J$ = 17.5, 8.8 Hz, 1H), 1.99–1.92 (m, 1H), 1.60–1.50 (m, 1H), 1.48–1.40 (m, 1H), 1.25–1.17 (m, 1H), 0.83 (t, $J$ = 7.4 Hz, 3H). Spectral and physical data agreed with those reported previously (Xie *et al.*, 2017).

**(2*S*,3*R*,11b*S*)-2-Allyl-3-ethyl-9,10-dimethoxy-1,3,4,6,7,11b-hexahydro-2*H*-pyrido[2,1-*a*]isoquinoline (S10)**

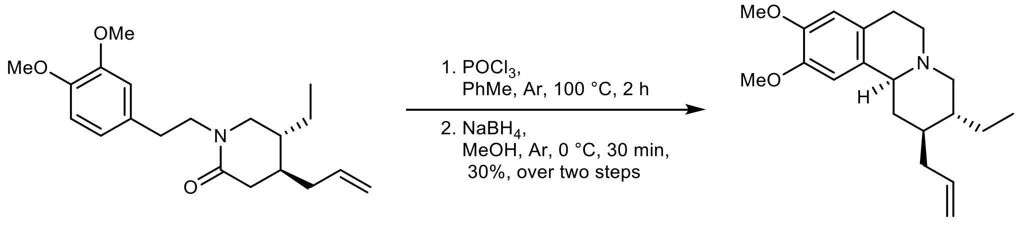

**S9** **S10**

Briefly following a previously reported procedure (Xie *et al.*, 2017), to a dry 50 mL round bottom flask under an Ar atmosphere, containing lactam **S9** (293 mg, 0.88 mmol) in anhydrous PhMe (10 mL), was added $POCl_3$ (0.33 mL, 3.52 mmol). The mixture was heated to 100 °C for 2 h and then concentrated *in vacuo* to afford a brown residue that was subsequently dissolved in anhydrous MeOH (10 mL). Upon cooling to 0 °C, the solution was charged with $NaBH_4$ (67 mg, 1.76 mmol) and then allowed to stir for a further 30 min. The resulting bright yellow solution was concentrated under reduced pressure to afford a residue that was partitioned between EtOAc (10 mL) and $H_2O$ (30 mL). The aqueous layer was extracted with EtOAc (10 mL x 2) and the resulting organic layers were combined, dried over anhydrous $MgSO_4$, and concentrated *in vacuo* to give a residue that was purified by gradient flash column chromatography (10–50% EtOAc in petroleum ether [b.p. 40–60 °C]) to afford title compound **S10** as a yellow oil (88 mg, 30% [over two steps from lactam **S9**]): $^1$H NMR ($CDCl_3$, 400 MHz) $\delta$ 6.68 (s, 1H), 6.56 (s, 1H), 5.89–5.78 (m, 1H), 5.06–5.02 (m, 2H), 3.84 (s, 3H), 3.83 (s, 3H), 3.14–3.03 (m, 3H), 2.97 (ddd, *J* = 11.2, 5.9, 1.6 Hz, 1H), 2.61 (d, *J* = 16.0 Hz, 1H), 2.46 (td, *J* = 11.5, 4.1 Hz, 1H), 2.42–2.37 (m, 1H), 2.26 (dt, *J* = 12.9, 3.1 Hz, 1H), 2.04–1.95 (m, 2H), 1.72–1.64 (m, 1H), 1.54–1.30 (m, 3H), 1.27–1.20 (m, 1H), 1.18–1.09 (m, 1H), 0.91 (t, *J* = 7.5 Hz, 3H). Spectral and physical data agreed with those reported previously (Xie *et al.*, 2017).

**2-[(2*R*,3*R*,11b*S*)-3-Ethyl-9,10-dimethoxy-1,3,4,6,7,11b-hexahydro-2*H*-pyrido[2,1-*a*]isoquinolin-2-yl]acetaldehyde (8)**

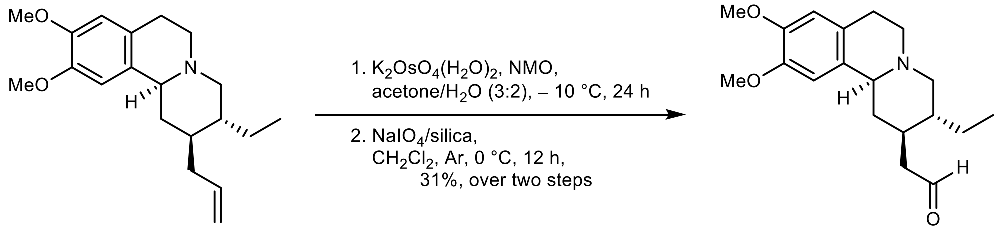

**S10** **8**

Following a previously reported procedure (Xie *et al.*, 2017), to a 50 mL round bottom flask, containing alkene **S10** (81 mg, 0.26 mmol) and *N*-methylmorpoline *N*-oxide (137 mg, 1.17 mmol) in acetone/$H_2O$ (3:2, 13 mL) at – 10 °C, was added $K_2OsO_4(H_2O)_2$ (5.75 mg, 0.016 mmol). After stirring for 12 h, a second portion of $K_2OsO_4(H_2O)_2$ was added to the reaction mass and the mixture was further stirred at – 10 °C. After 12 h, the resulting solution was diluted with *n*-BuOH (10 mL) and charged with 0.4 M aq. $Na_2SO_3$ (10 mL). Repeated extraction of the aforementioned mixture with *n*-BuOH (10 mL x 3) gave a combined organic phase that was subsequently washed with sat. aq. $NaHCO_3$ (10 mL) and brine (10 mL) and then dried over anhydrous $Na_2SO_4$. Concentration under reduced pressure afforded a residue that was dissolved in anhydrous $CH_2Cl_2$ (6.5 mL) and cooled to

0 °C under Ar. The solution was charged with freshly prepared silica-supported $NaIO_4$ (Zhong & Shing, 1997) (819 mg, 0.26 mmol) and stirred at 0 °C for 12 h. Filtration of the reaction mixture through a short pad of Celite® gave a filtrate that was concentrated *in vacuo* to afford a residue that was further purified by gradient flash column chromatography (0–6.25% MeOH in EtOAc) to furnish (−)-protoemetine (**8**) as a light yellow oil (25 mg, 31% [over two steps from alkene **S10**]): NMR data is described in Supplementary Fig. 27. HRMS (ESI/Q-TOF) *m/z*: $[M+H]^+$ Calcd. for $C_{19}H_{28}NO_3$ 318.2064; Found 318.2059 (− 1.6 ppm). Spectral and physical data agreed with those reported previously (Lin *et al.*, 2011).

#### Supplementary Figures

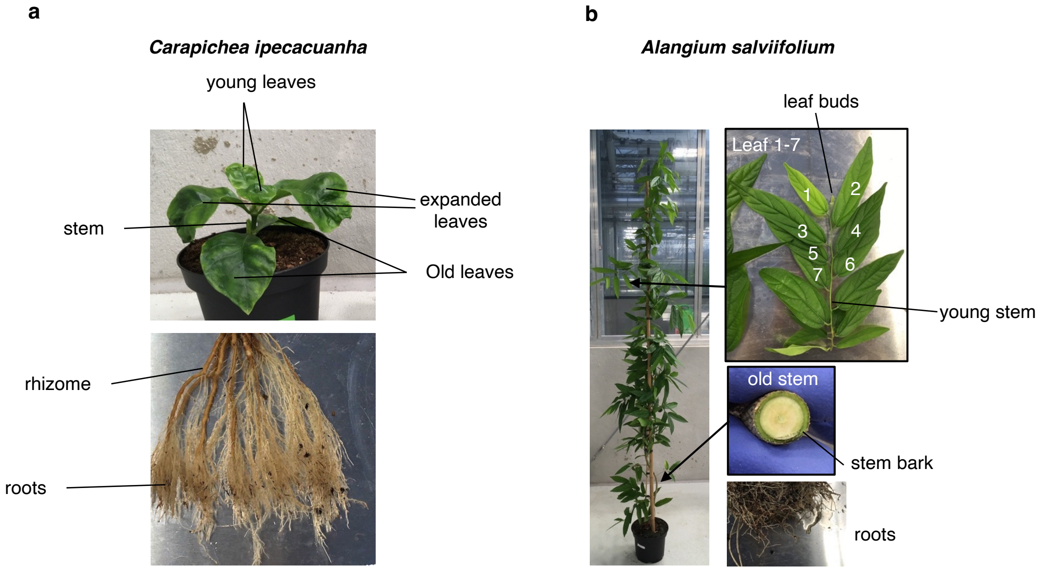

**Supplementary Fig. 1. Photos of plants sampled for metabolite analysis and RNA-seq. a**, *C. ipecacuanha* plants were regenerated from *in vitro* culture and grown on soil for four months. **b**, *A. salviifolium* plants were grown from cuttings harvested after 14 months. Plants were grown under controlled conditions in a greenhouse (12/12 hours light/dark 28-30/ 24-26°C°). Indicated tissues were used for RNA-seq and metabolomics analysis. Metabolomics was performed using three individual plants of the same age, RNA-seq was performed using material from one individual plant (shown on the picture).

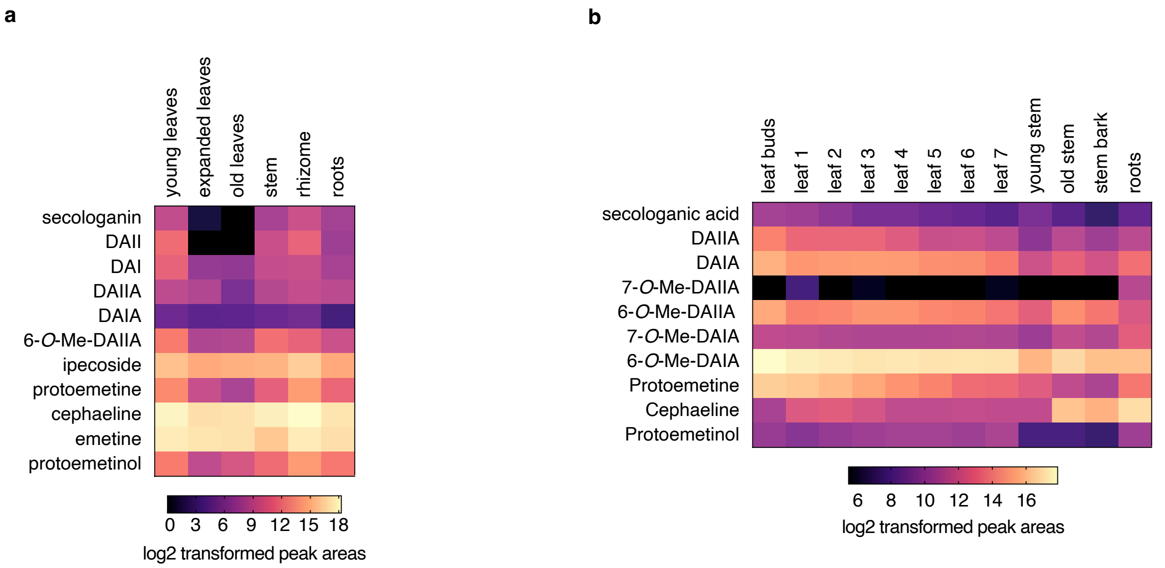

**Supplementary Fig. 2. Extended metabolite content of *C. ipecacuanha* and *A. salviifolium* tissues.** Heatmap depicts log2 transformed mean of LC-MS peak areas from three biological replicates. Data is from the same experiment shown in main text Fig. 2 f-g.

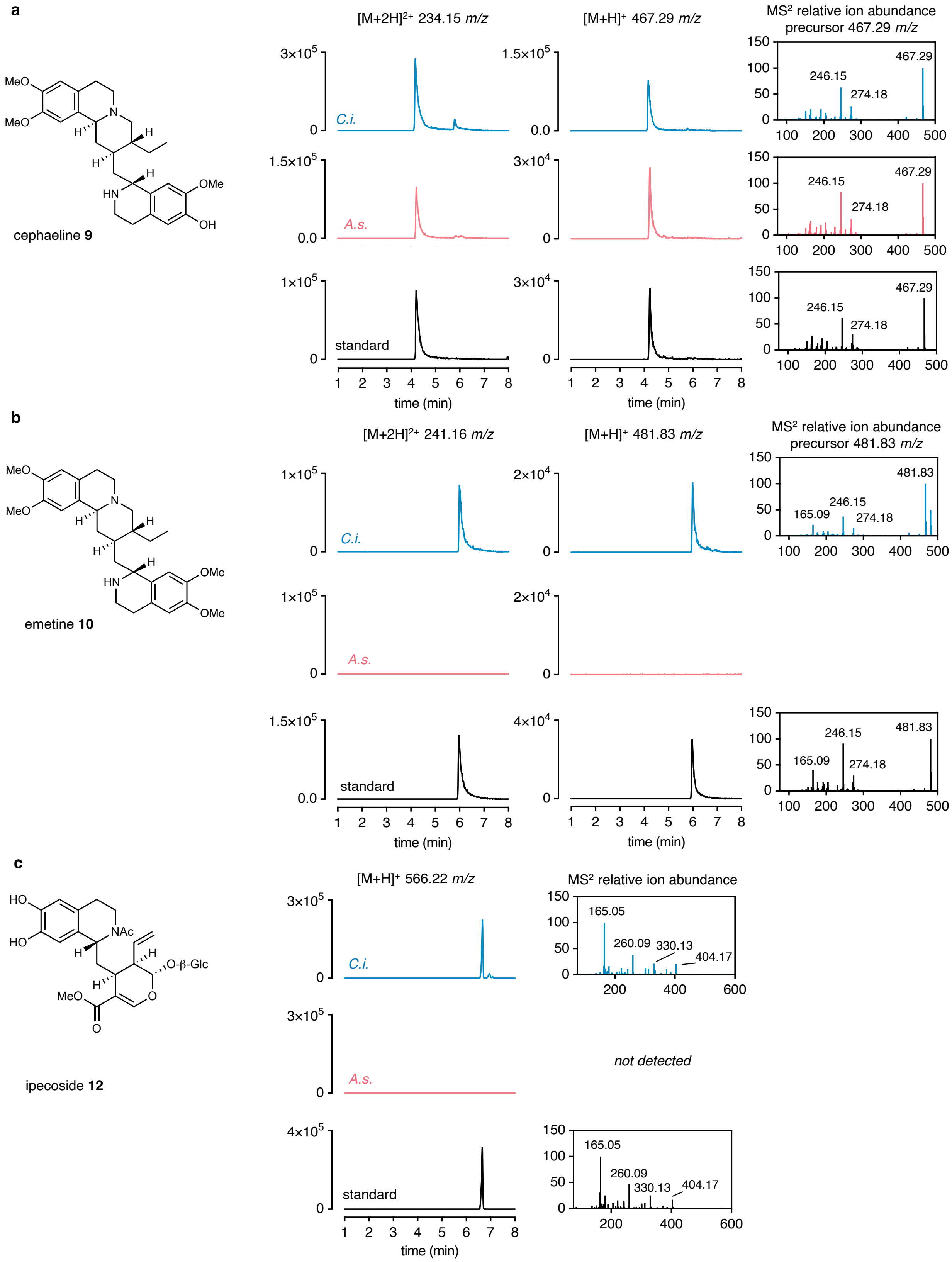
a
cephaeline 9
[M+2H]2+ 234.15 m/z
[M+H]+ 467.29 m/z
MS2 relative ion abundance precursor 467.29 m/z
C.i.
A.s.
standard
467.29
246.15
274.18
time (min)
b
emetine 10
[M+2H]2+ 241.16 m/z
[M+H]+ 481.83 m/z
MS2 relative ion abundance precursor 481.83 m/z
481.83
165.09
246.15
274.18
c
ipecoside 12
[M+H]+ 566.22 m/z
MS2 relative ion abundance
165.05
260.09
330.13
404.17
not detected

**Supplementary Fig. 3. Extracted ion chromatograms (EIC) and MS$^2$ data of cephaeline, emetine and ipecoside of A. salviifolium roots and C. ipecacuanha rhizome compared to purchased authentic standards. a,** cephaeline, **b,** emetine, **c**, ipecoside. All chromatograms are extracted from the same LC-MS experiment, which is the same as shown in main text Fig. 2a. EIC for *C. ipecacuanha* extract is shown in blue and labelled with *C.i.* as abbreviation; for *A. salviifolium* in magenta, abbreviated with *A.s.*; and for the respective standards in black. For cephaeline and emetine $[M+2H]^{2+}$ and $[M+H]^{+}$ ions are detected. MS$^2$ data is shown as relative abundance of ions with *m/z* values of the most abundant fragment ions indicated. Ipecoside and emetine were not detected in *A. salviifolium* extracts.

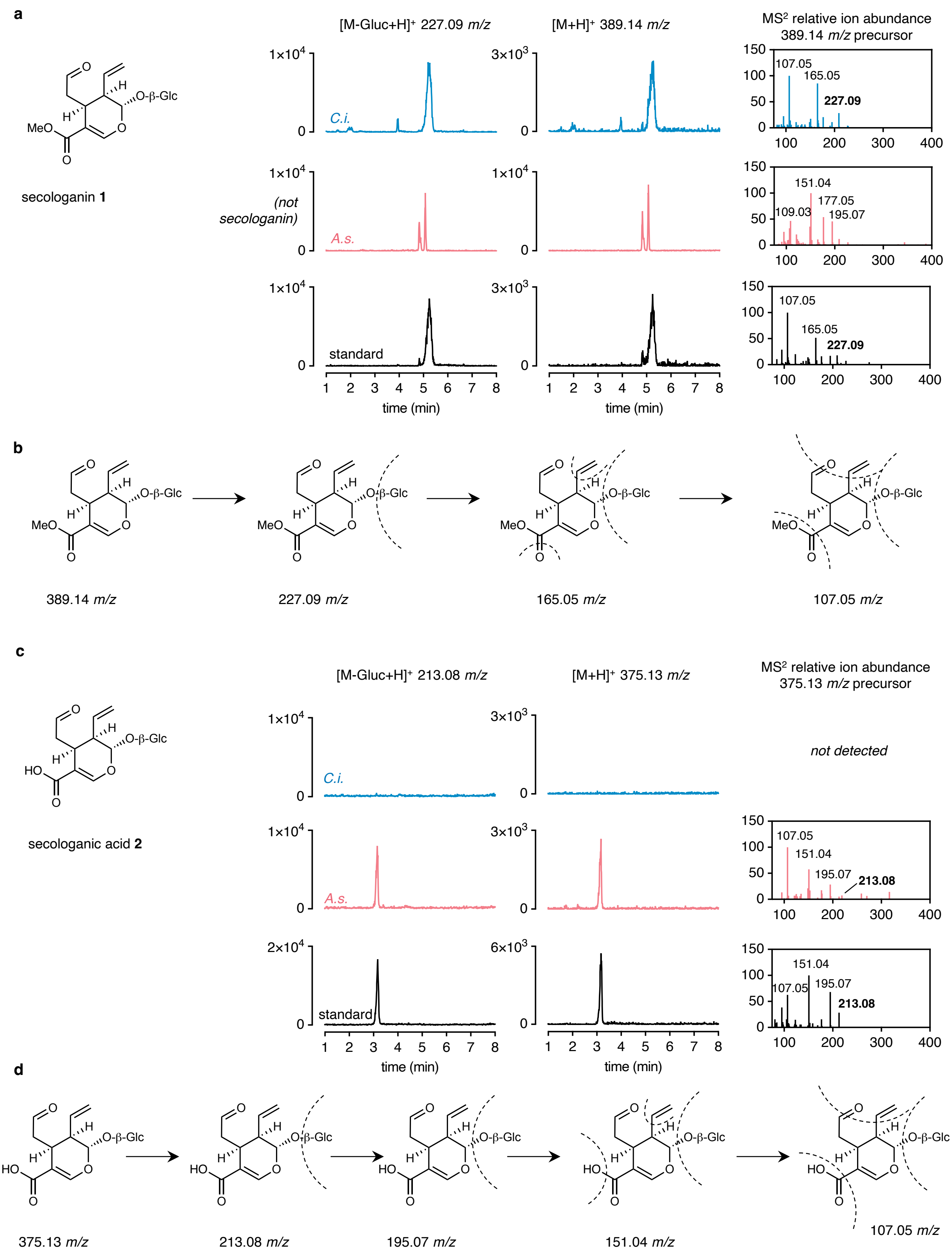

a
secologanin 1
[M-Gluc+H]+ 227.09 m/z
[M+H]+ 389.14 m/z
MS2 relative ion abundance
389.14 m/z precursor
C.i.
(not secologanin)
A.s.
standard
107.05
165.05
227.09
151.04
177.05
109.03
195.07
time (min)
b
389.14 m/z
227.09 m/z
165.05 m/z
107.05 m/z
c
secologanic acid 2
[M-Gluc+H]+ 213.08 m/z
[M+H]+ 375.13 m/z
MS2 relative ion abundance
375.13 m/z precursor
not detected
213.08
d
375.13 m/z
213.08 m/z
195.07 m/z
151.04 m/z
107.05 m/z

**Supplementary Fig. 4. Extracted ion chromatograms and MS$^2$ data of secologanin and secologanic acid of *A. salviifolium* roots and *C. ipecacuanha* rhizome compared to standards. a**, data for secologanin. Authentic standard is commercially available. An in source fragment ion (chromatogram on the left), likely secologanin aglycone, is detetected in addition to the $[M+H]^+$ ion (chromatogram on the right). An *m/z* identical to the MS$^1$ of secologanin is detected in *A. salviifolium* extracts but peak shape, retention time and MS$^2$ fragmentation pattern indicate that this compound is not secologanin. These compounds are likely vogeloside and epivogeloside that have previously been described to form from secologanic acid (Yeon *et al*, 2018). We have also observed these peaks when secologanic acid standard is diluted in methanol prior to LC-MS analysis. **b,** proposed fragmentation of secologanin. **c**, data for secologanic acid. An in source fragment ion (chromatogram on the left), presumably secologanic acid aglycone, is detected in addition to the $[M+H]^+$ ion (chromatogram on the right). **d**, Proposed MS$^2$ fragmentation pattern for secologanic acid. Secologanic acid standard was produced from purchased secologanin through alkaline hydrolysis (see methods). LC-MS and MS$^2$ data of the confirmed the correct identity of secologanic acid as the resulting product. *C. ipecacuanha* data is shown in blue and labelled with *C.i.* as abbreviation; *A. salviifolium* data in magenta, abbreviated with *A.s.*; and data for the respective standards in black. MS$^2$ data is shown as relative abundance of ions with *m/z* values of the most abundant fragment ions indicated.

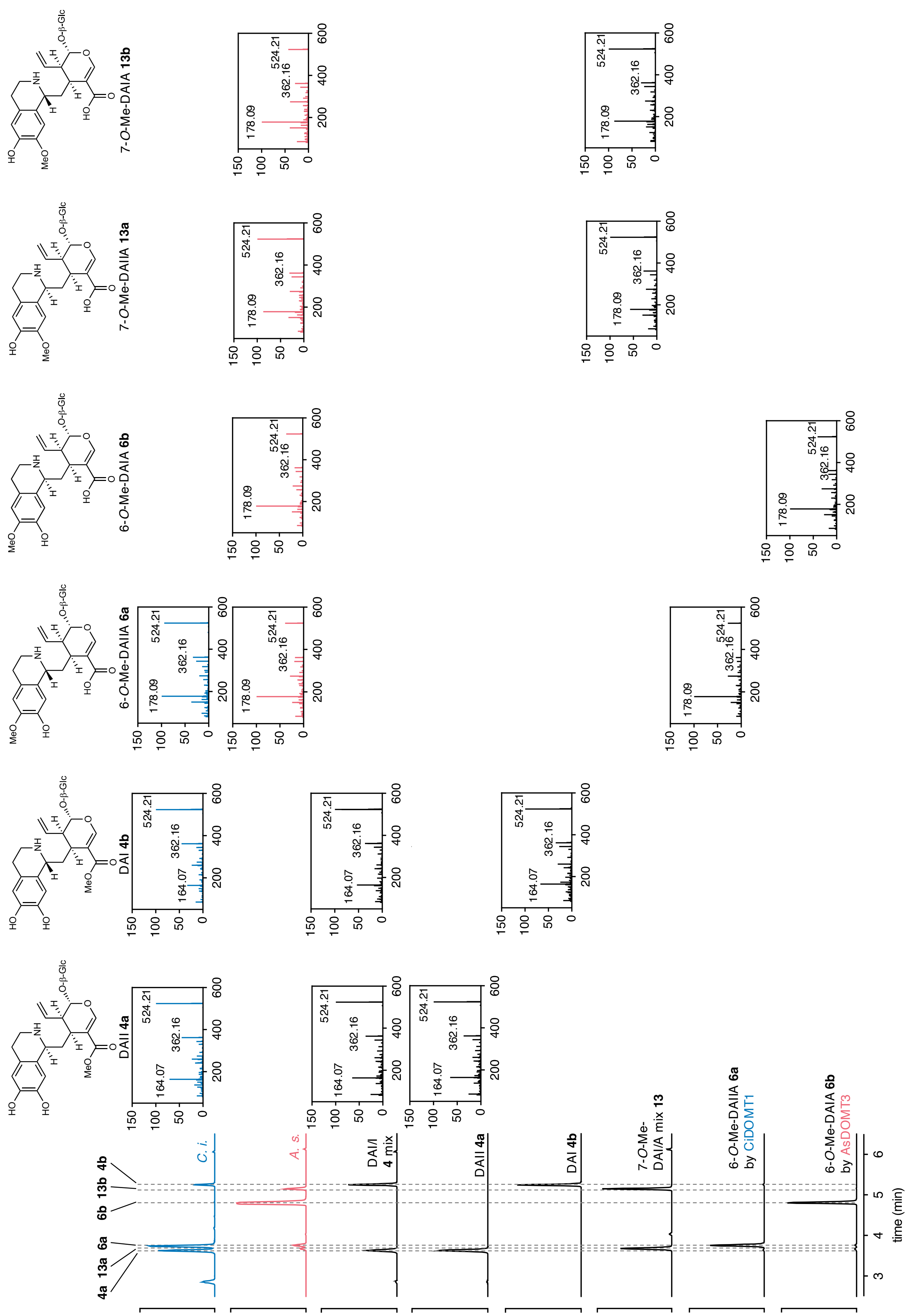
DAII 4a
DAI 4b
6-O-Me-DAIIA 6a
6-O-Me-DAIA 6b
7-O-Me-DAIIA 13a
7-O-Me-DAIA 13b
O-β-Glc
164.07
178.09
362.16
524.21
C. i.
A. s.
DAI/I 4 mix
7-O-Me-DAI/A mix 13
6-O-Me-DAIIA 6a by CiDOMT1
6-O-Me-DAIA 6b by AsDOMT3
time (min)

**Supplementary Fig. 5. Extracted ion chromatograms and $MS^2$ data of compounds with same m/z DA(I)I epimers and 6-, or 7-O-methylated DA(I)A epimers of *A. salviifolium* roots and *C. ipecacuanha* rhizome.** Left: EICs for 524.22 *m/z*. From top to bottom: *C. ipecacuanha* data (*C.i.*, blue), *A. salviifolium* (*A.s.*, magenta), DAII/DAI **4a/4b** epimer mix, purified DAII **4a** (stereochemistry confirmed by NMR), purified DAI **4b** (stereochemistry confirmed by NMR), 7-*O*-Me-DAI/IA **13a/b** epimer mixture, tentative 6-*O*-Me-DAIIA **6a** produced by recombinant CiDOMT1, and tentative 6-*O*-Me-DAIA **6b** produced by recombinant AsDOMT3 (see methods). The mixtures of epimers were produced by Pictet-Spengler reaction of dopamine with secologanin in case of DAI/I or 4-*O*-Me-dopamine with secologanic acid for DAIA/DAIIA (see methods). The reactions yield *S* and *R* epimers at a ratio of approximately 40:60. Minor peaks with the same *m/z* values are consistently observed when performing these reactions and could correspond to “neo” isomers which were previously described to be minor by products of the reaction of the similar DAII and DAI (Beke *et al.*, 2001). On top the molecular structure of each detected molecule is shown. $MS^2$ fragmentations are shown wherever each of these molecules were detected. The main distinction criterion between *O*-methylated acid epimers and DAI/I is a fragment derived from the dopamine moiety which is 178.09 *m/z* when this moiety is methylated and 164.07 *m/z* when not methylated. These signals are labelled in each $MS^2$ fragmentation. The difference in *m/z* is indicative of a methyl group (+14). $MS^2$ fragmentation for 7-*O*-Me-DAI(I)A epimers and tentative 6-*O*-Me-DAI(I)A is identical but the retention time is shifted. Therefore, it can be assumed that the indicated peaks are indeed the 6-*O*-methylated compounds. $MS^2$ data is shown as relative abundance of ions with *m/z* values of the most abundant fragment ions indicated.

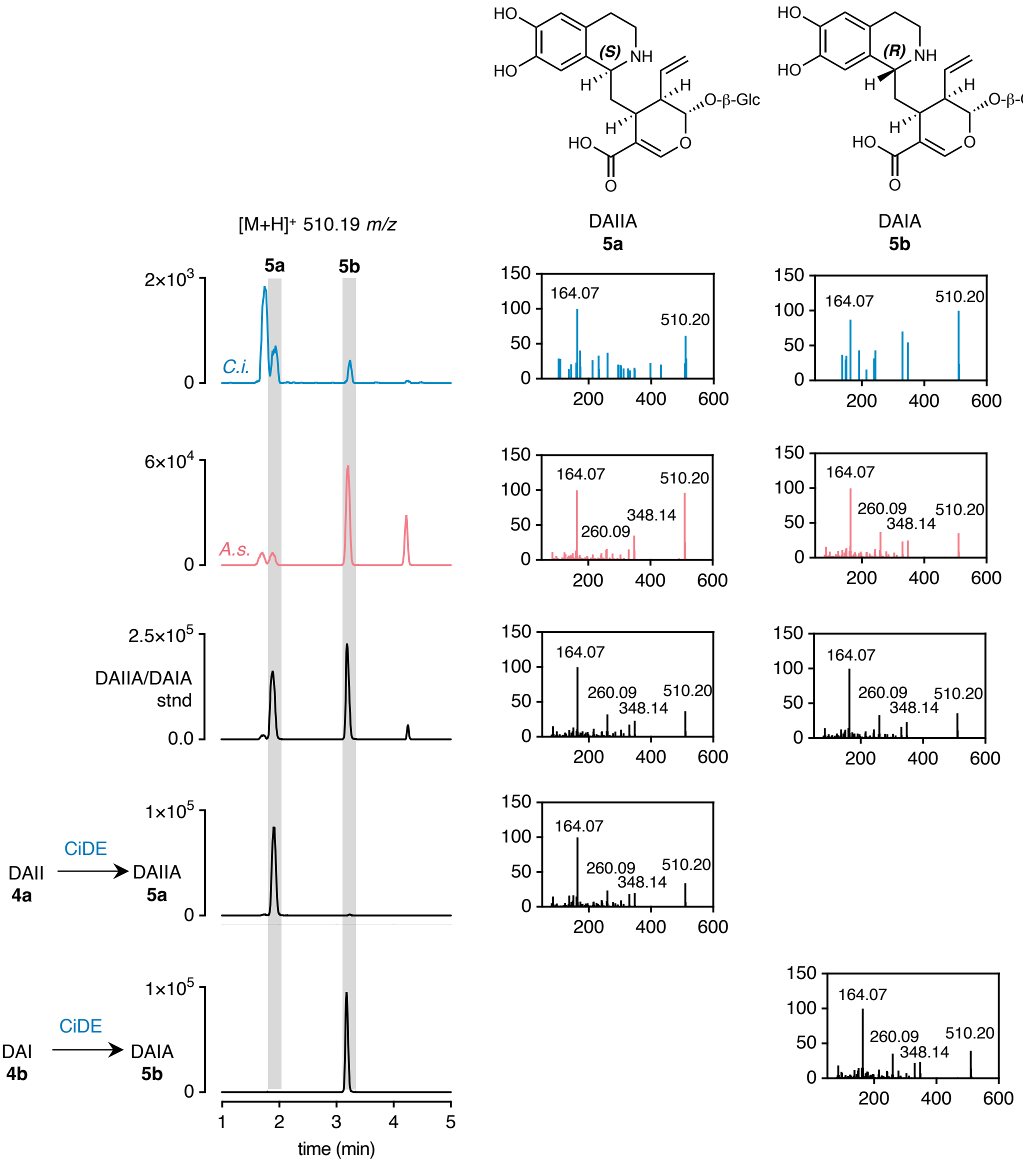

**Supplementary Fig. 6. Extracted ion chromatograms and MS$^2$ data of deacetylisoipecosidic acid (DAIIA, 5a) and deacetylipecosidic acid (DAIA, 5b) of *A. salviifolium* roots and *C. ipecacuanha* rhizome compared to standards.** The mixture of epimers was produced by Pictet-Spengler reaction of dopamine with secologanic acid (see methods). The reactions yield *S* and *R* epimers at a ratio of approximately 40:60. Minor peaks are with the same *m/z* values are consistently observed when performing these reactions and could correspond to "neo" isomers which were previously described (Beke *et al.*, 2001). To confirm epimer specific peak identity epimer pure DAII (**4a**) or DAI (**4b**) (for which stereochemistry was confirmed by NMR, see below), respectively, were deesterified using recombinant CiDE. This confirmed the first peak as *S* epimer and the second as the *R* epimer. DAIIA (**5a**) and DAIA (**5b**) are also observed in *C. ipecacuanha*, albeit in low amounts. *C. ipecacuanha* data is shown in blue and labelled with *C.i.* as abbreviation; *A. salviifolium* data in magenta, abbreviated with *A.s.*; and data for the respective standards in black as indicated. MS$^2$ data is shown as relative abundance of ions with *m/z* values of the most abundant fragment ions indicated.

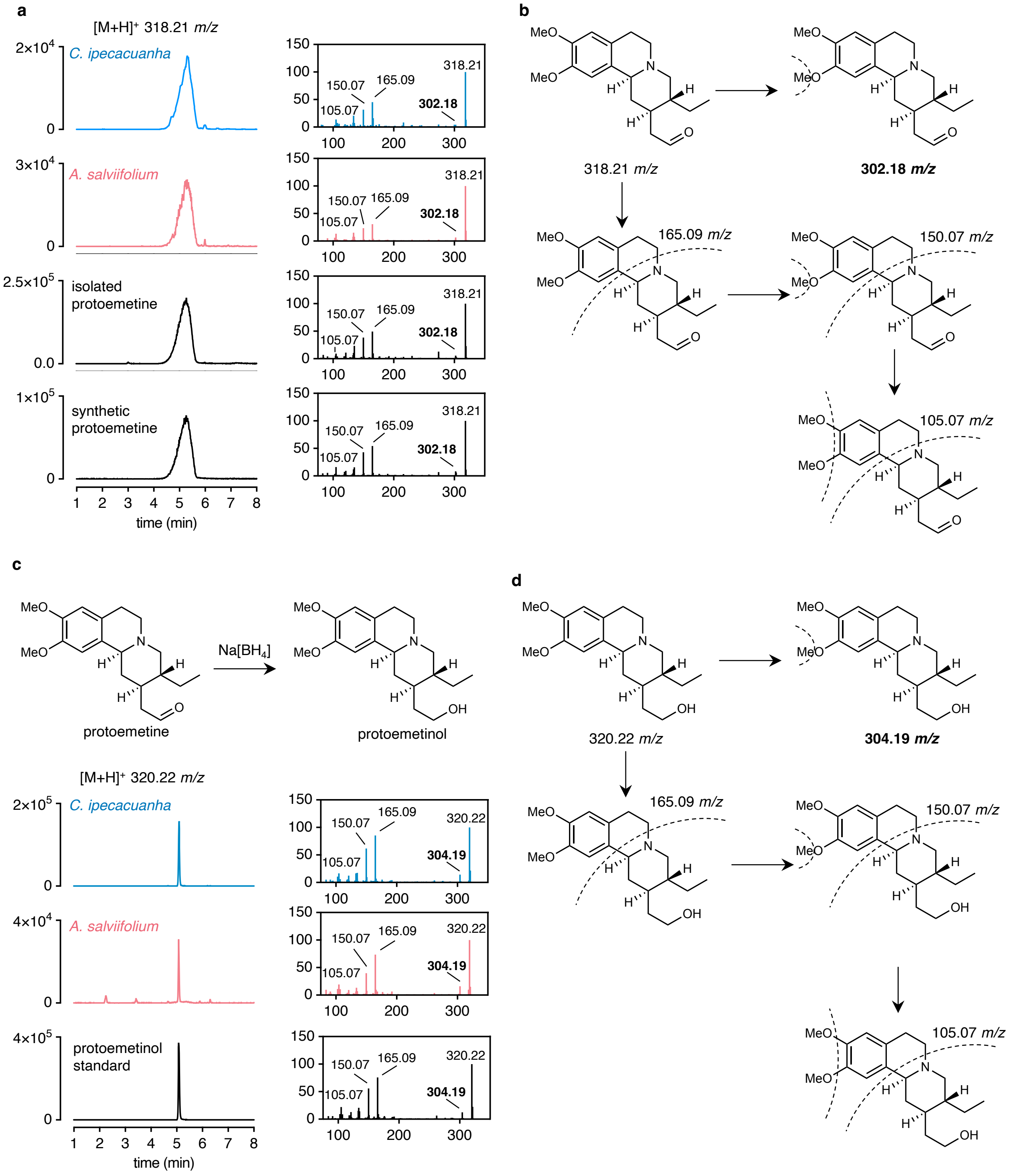

**Supplementary Fig. 7. Extracted ion chromatograms and MS$^2$ data of protoemetine and protoemetinol of *A. salviifolium* roots and *C. ipecacuanha* rhizome compared to standards. a**, data for protoemetine. Two chemically identical protoemetine standards were used throughout this study. A limited amount of protoemetine was isolated from *A. salviifolium* leaf buds and NMR data was in agreement with published data. An additional standard was obtained through chemical synthesis (see methods, full NMR spectra see Supplementary Fig. 24). **b**, proposed fragmentation of protoemetine. **c**, data for protoemetinol. Protoemetinol was produced through $NaBH_4$ reduction of protoemetine. MS data was consistent with values expected for protoemetinol. **d**, proposed MS$^2$ fragmentation of protoemetinol. *C. ipecacuanha* data is shown in blue and labelled with *C.i.* as abbreviation; *A. salviifolium* data in magenta, abbreviated with *A.s.*; and data for the respective standards in black as indicated. MS$^2$ data is shown as relative abundance of ions with *m/z* values of the most abundant fragment ions indicated and additionally a fragment that shows expected difference between protoemetine and protoemetinol.

**a**

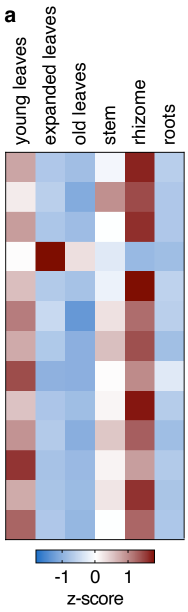

| Contig ID | Pearson | Name / annotation | % identity to bait | Name bait | Bait published/ enzyme class |
|---|---|---|---|---|---|
| NODE_49441_length_1590_cov_622.147552_g5127_i1 | 1 (bait) | **CiDOMT1** | 97.2 (DNA) | IpeOMT1 | Nomura et al. 2010 Class II OMT |
| NODE_48823_length_1616_cov_3020.873434_g2542_i2 | 0.85 | **CiDOMT2** | 98.7 (DNA) | IpeOMT2 | |
| NODE_52287_length_1477_cov_2155.816401_g14913_i1 | 0.99 | **CiDPOMT** | 98.5 (DNA) | IpeOMT3 | |
| NODE_39259_length_2017_cov_2392.595933_g810_i14 | -0.49 | **CiDGD** | 96.8 (DNA) | IpeGlu1 | Nomura et al. 2008 GH-1 |
| NODE_32402_length_2349_cov_4818.038222_g10846_i2 | 0.99 | **CiS6DGD** | 71.8 (P) | | |
| NODE_17451_length_3362_cov_3620.589335_g2352_i2 | 0.93 | **CiDE** | 42.1 (P) | CpDCE | Trenti et al. 2021 esterase |
| NODE_40827_length_1951_cov_351.045896_g560_i2 | 0.94 | esterase | 42.6 (P) | | |
| NODE_45313_length_1757_cov_1066.269602_g8912_i3 | 0.88 | esterase | 37.5 (P) | | |
| NODE_29146_length_2518_cov_1102.449773_g3581_i2 | 0.98 | **CiDR1** | 59.6 (P) | CpDCS | Trenti et al. 2021 MDR |
| NODE_60625_length_1162_cov_1650.389464_g21999_i1 | 0.94 | CiDR2 | 60.8 (P) | | |
| NODE_27176_length_2633_cov_1787.503552_g11761_i0 | 0.87 | **CiIpS** | | | BAHD |
| NODE_40057_length_1983_cov_3156.798301_g3796_i6 | 0.98 | SLS | | CrSLS | P450 |
| NODE_51992_length_1489_cov_1828.264029_g13669_i1 | 0.95 | TyrDC | | | decarboxylase |

**b**

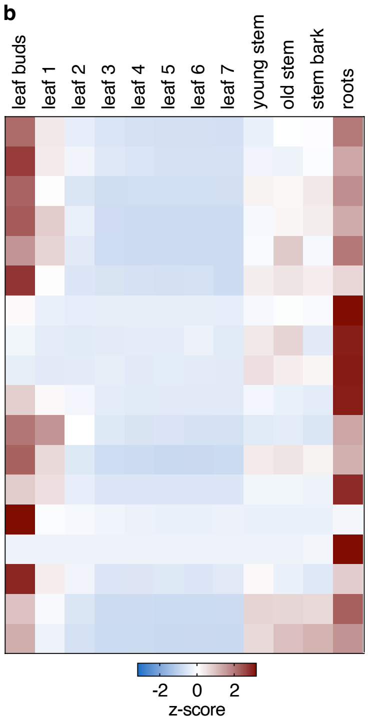

| Contig ID | Pearson | Name / annotation | Enzyme class |
|---|---|---|---|
| NODE_37865_length_2861_cov_2825.464156_g5654_i1 | 1 (bait) | TyrDC | Decarboxylase |
| NODE_29512_length_3407_cov_8690.483071_g9907_i5 | 0.96 | SLAS | P450 |
| NODE_19671_length_4252_cov_2535.745726_g6537_i3_orf2 | 0.97 | **AsDOMT3** | Class I OMT |
| NODE_58023_length_1865_cov_2571.295583_g11076_i4 | 0.96 | **AsDOMT4** | |
| NODE_86485_length_811_cov_4484.609551_g31936_i0 | 0.94 | **AsDOMT5** | |
| NODE_19671_length_4252_cov_2535.745726_g6537_i3_orf1 | 0.87 | **AsDOMT1** | |
| NODE_72219_length_1312_cov_4254.701566_g11513_i1 | 0.7 | **AsDOMT6** | |
| NODE_45199_length_2456_cov_116.731438_g12818_i3 | 0.6 | **AsDOMT7** | |
| NODE_66616_length_1523_cov_241.251404_g12818_i4 | 0.58 | **AsDOMT2** | |
| NODE_55469_length_1972_cov_6623.087026_g11371_i7 | 0.83 | **AsDPOMT1** | Class II OMT |
| NODE_56906_length_1913_cov_2483.830761_g11371_i8 | 0.89 | **AsDPOMT2** | |
| NODE_48307_length_2301_cov_2849.912352_g10896_i4 | 0.94 | GH-3 | GH-3 |
| NODE_18446_length_4378_cov_1156.998832_g3211_i3 | 0.88 | **AsS6DGD** | |
| NODE_48420_length_2294_cov_1341.643736_g14816_i1 | 0.69 | **AsDGD1** | GH-1 |
| NODE_50890_length_2176_cov_2234.009148_g11790_i2 | 0.61 | **AsDGD2** | |
| NODE_56935_length_1912_cov_549.413127_g11941_i1 | 0.88 | **AsDR1** | MDR |
| NODE_34732_length_3050_cov_6904.524229_g4904_i3 | 0.83 | **AsDR2** | |
| NODE_65175_length_1578_cov_5651.869506_g19320_i1 | 0.78 | dehydrogenase | |

**Supplementary Fig. 8. Selection of pathway gene candidates. a**, selection of *C. ipecacuanha* candidates by searching for genes similar to previously published active pathway enzymes (Nomura & Kutchan, 2010; Nomura *et al.*, 2008)). For each of the published sequences the contig with the highest sequencing homology (>96%) at cDNA level was considered to be identical to the published enzyme. *CiDOMT1* was used as a bait for Pearson correlation. Candidates for deesterification and reduction were found by homology-based search and coexpression with *CiDOMT1* (described in in the methods). A highly coexpressed gene predicted to encode for an acetyltransferase was picked as a candidate for ipecoside synthase. **b**, selection of *A. salviifolium* candidates was performed by coexpression analysis using orthologous and likely conserved sequences of enzymes supplying the direct precursor genes *TyrDC* and *SLS*, as baits, combined with filtering for predicted functional annotations for *O*-methyltransferases, glucosidases and dehydrogenases. Additionally, we considered candidates that showed either root or leaf bud specific expression (*AsDGDs*). Additional root specific *AsDOMT5-7* were also identified. For both species gene expression patterns are consistent with metabolite abundance (see Figure 2). Heatmaps depict z-scores of TMM normalized CPM values for each gene.

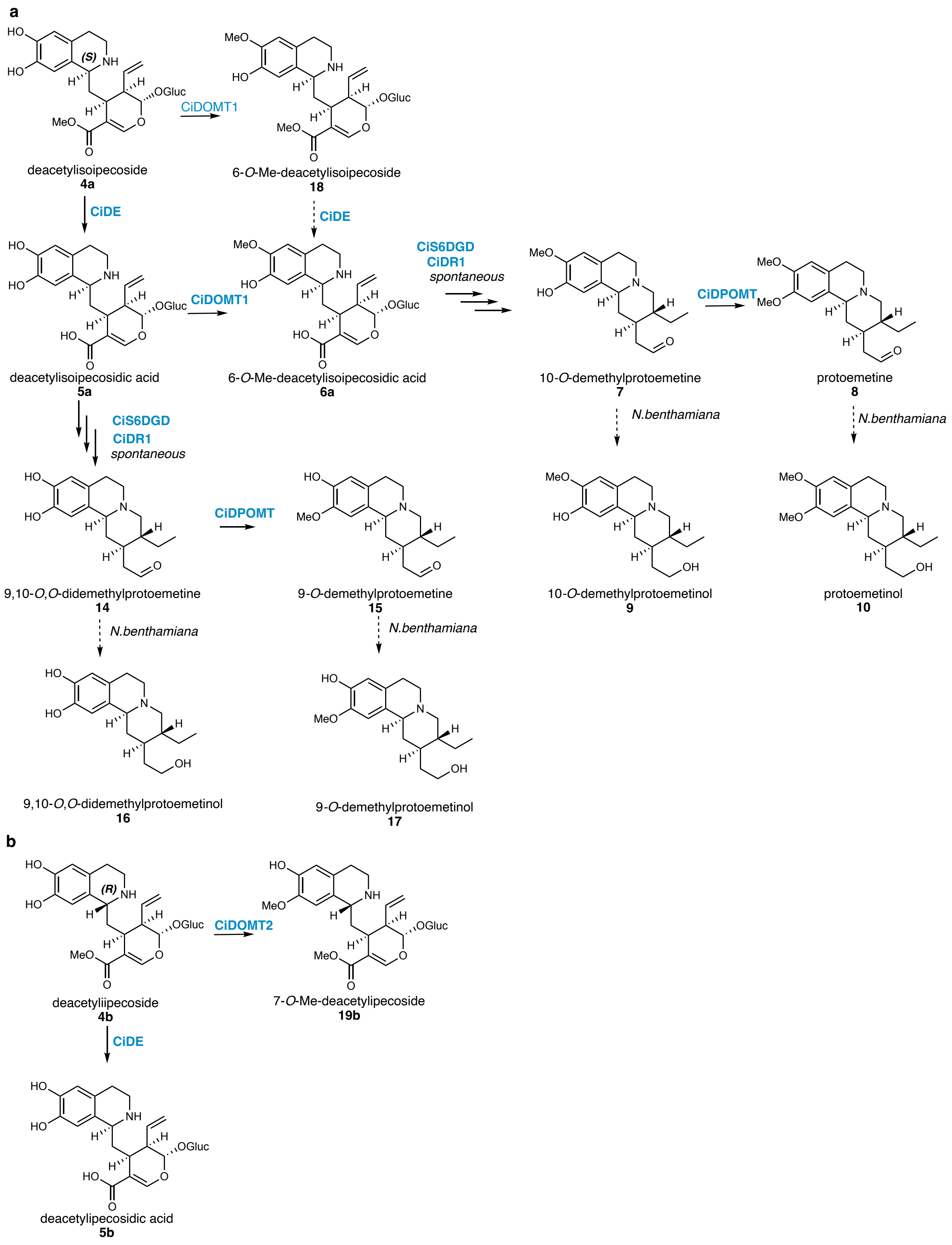

**Supplementary Fig. 9. *C. ipecauanha* protoemetine biosynthesis network including possible side branch products.** Activities were detected through expression of pathway genes shown in Supplementary Fig. 10.

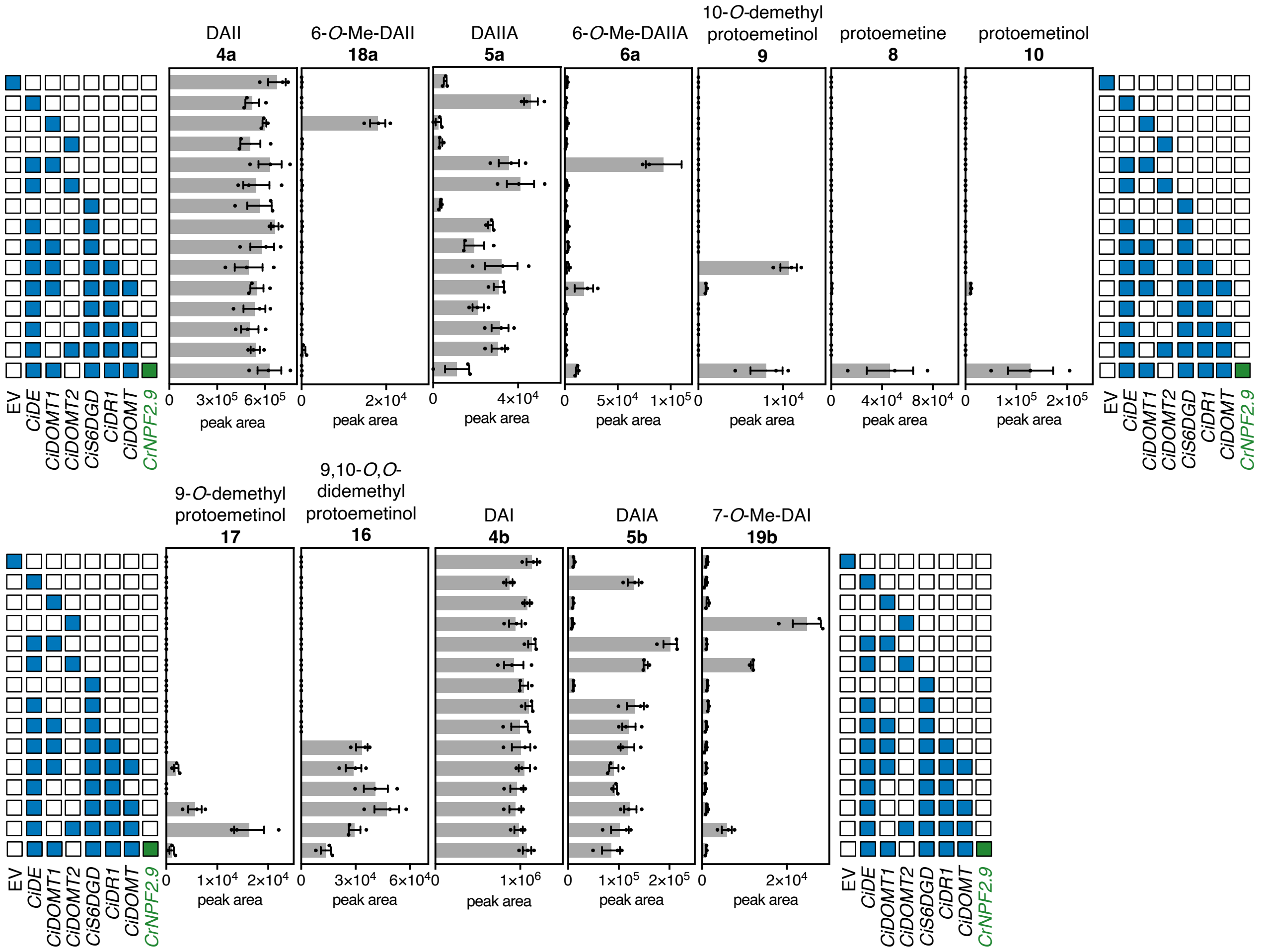

**Supplementary Fig. 10. Reconstitution of protoemetine biosynthesis network by expression of *C. ipecacuanha* pathway genes.** Data is from the same experiment shown in Fig. 3. This figure includes the data for Fig. 3 and additional gene combinations. Additionally, expression was combined with *Catharanthus roseus* strictosidine exporter CrNPF2.9 (Payne *et al*, 2017). LC-MS peak areas are shown as bars of the mean of three biological replicates, error bars are standard error of the mean.

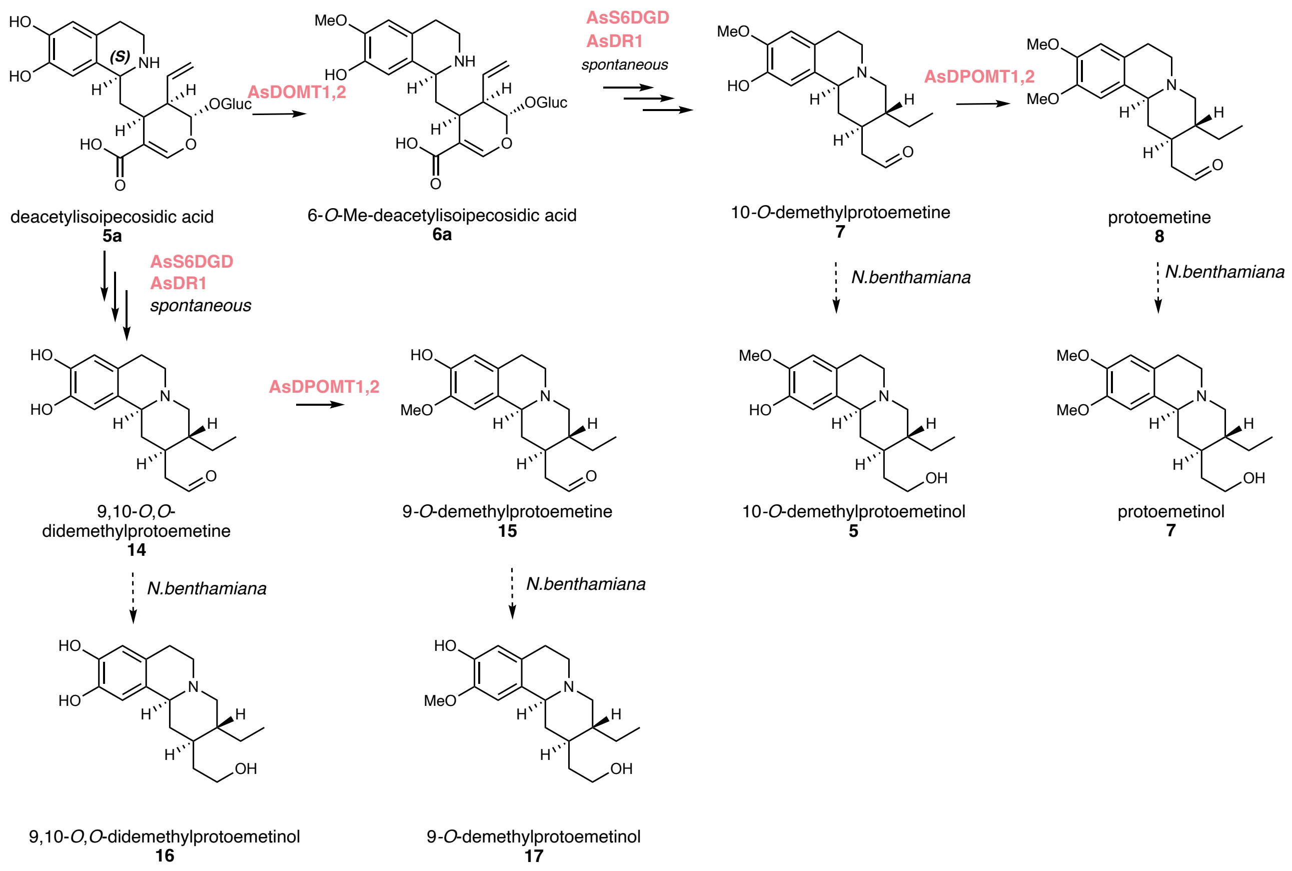

**Supplementary Fig. 11. *A. salviifolium* protoemetine biosynthesis network.** Activities were detected through expression of pathway genes shown in Supplementary Fig. 12.

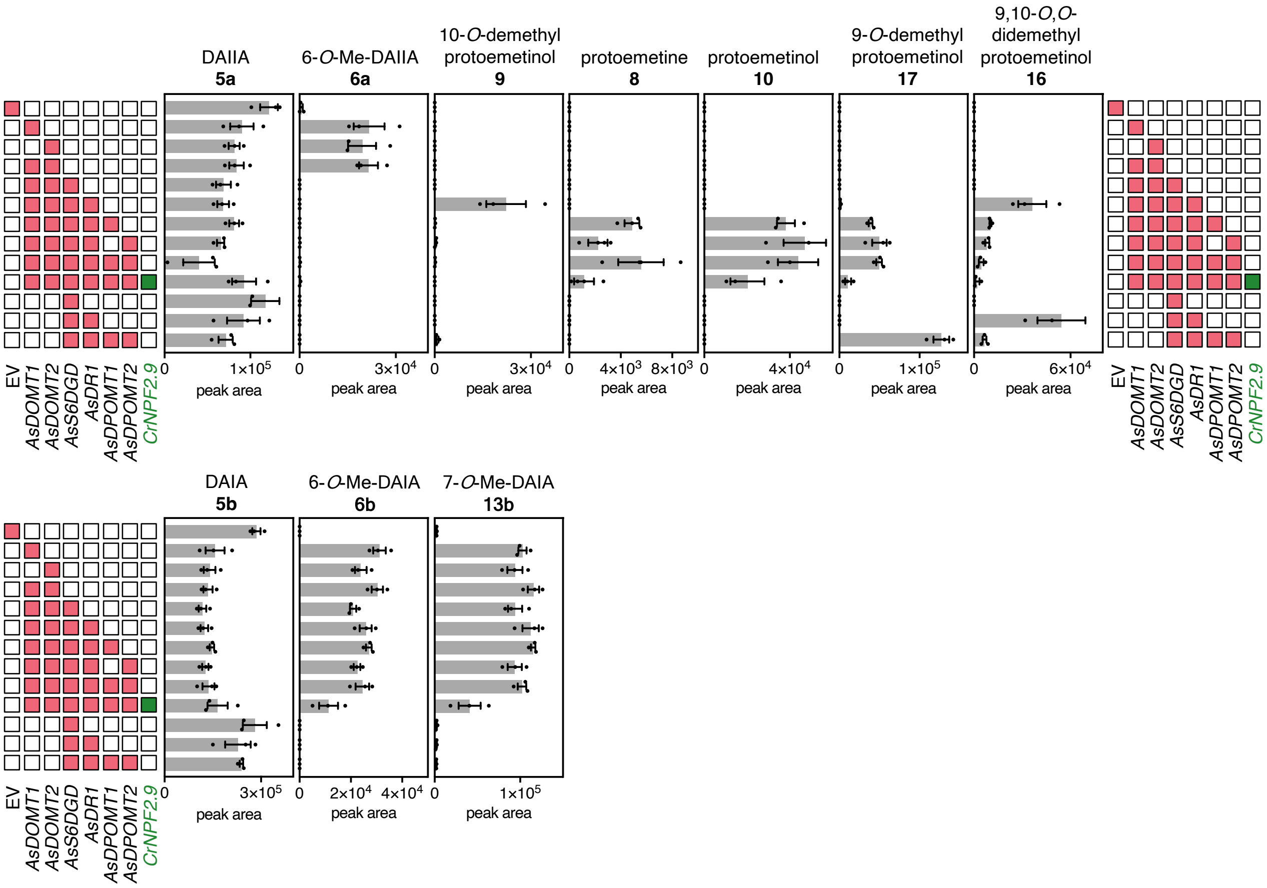

**Supplementary Fig. 12. Reconstitution of protoemetine biosynthesis network by expression of *A. salviifolium* pathway genes.** Data is from the same experiment shown in Fig. 3. This figure includes the data for Fig. 3 and additional gene combinations. Additionally, expression was combined with *Catharanthus roseus* strictosidine exporter CrNPF2.9 (Payne *et al.*, 2017). LC-MS peak areas are shown as bars of the mean of three biological replicates, error bars are standard error of the mean.

a

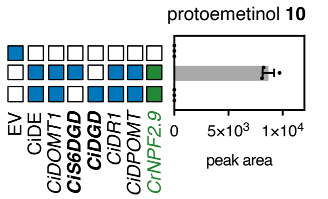

b

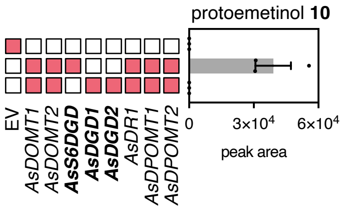

**Supplementary Fig. 13. Pathway reconstitution with CiDGD and AsDGD1,2 does not lead to protoemetine formation. a,** when *CiDGD* (Nomura et al. 2008) is expressed with the indicated pathway genes, no protoemetinol is formed, confirming that this glucosidase is not on protoemetine pathway. **b**, similarly, when *AsDGD1,2* are expressed with the indicated pathway genes no protoemetinol is formed. LC-MS peak areas are shown as bars of the mean of three biological replicates, error bars are standard error of the mean.

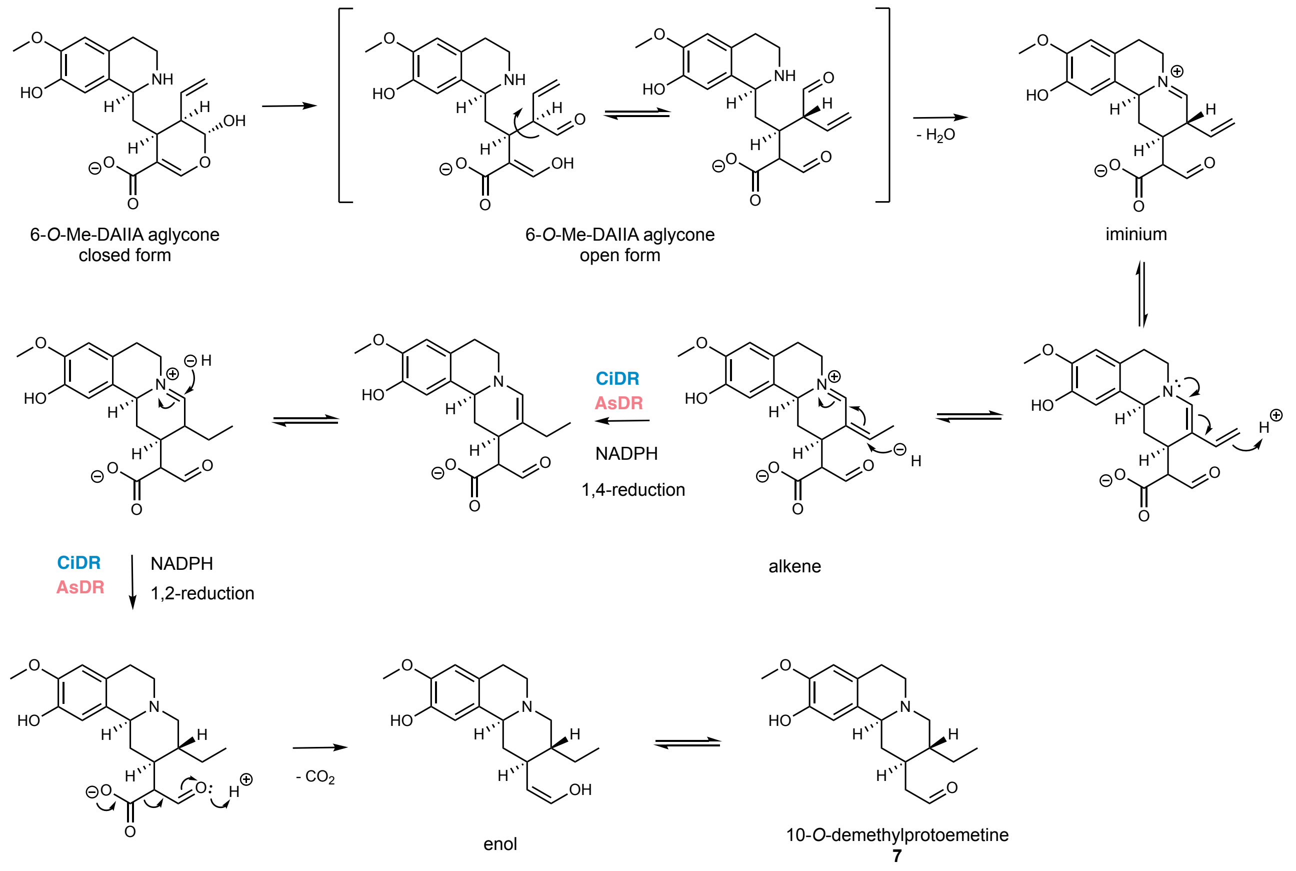

**Supplementary Fig. 14. Putative reduction mechanism of DRs and spontaneous decarboxylation.** 6-*O*-Me-DAIIA aglycone forms an iminium ion. After tautomerization, DR would first perform a 1,4 reduction followed by a 1,2 reduction. Spontaneous decarboxylation would then occur and lead to the enol which is in equilibrium with 10-*O*-demethylprotoemetine.

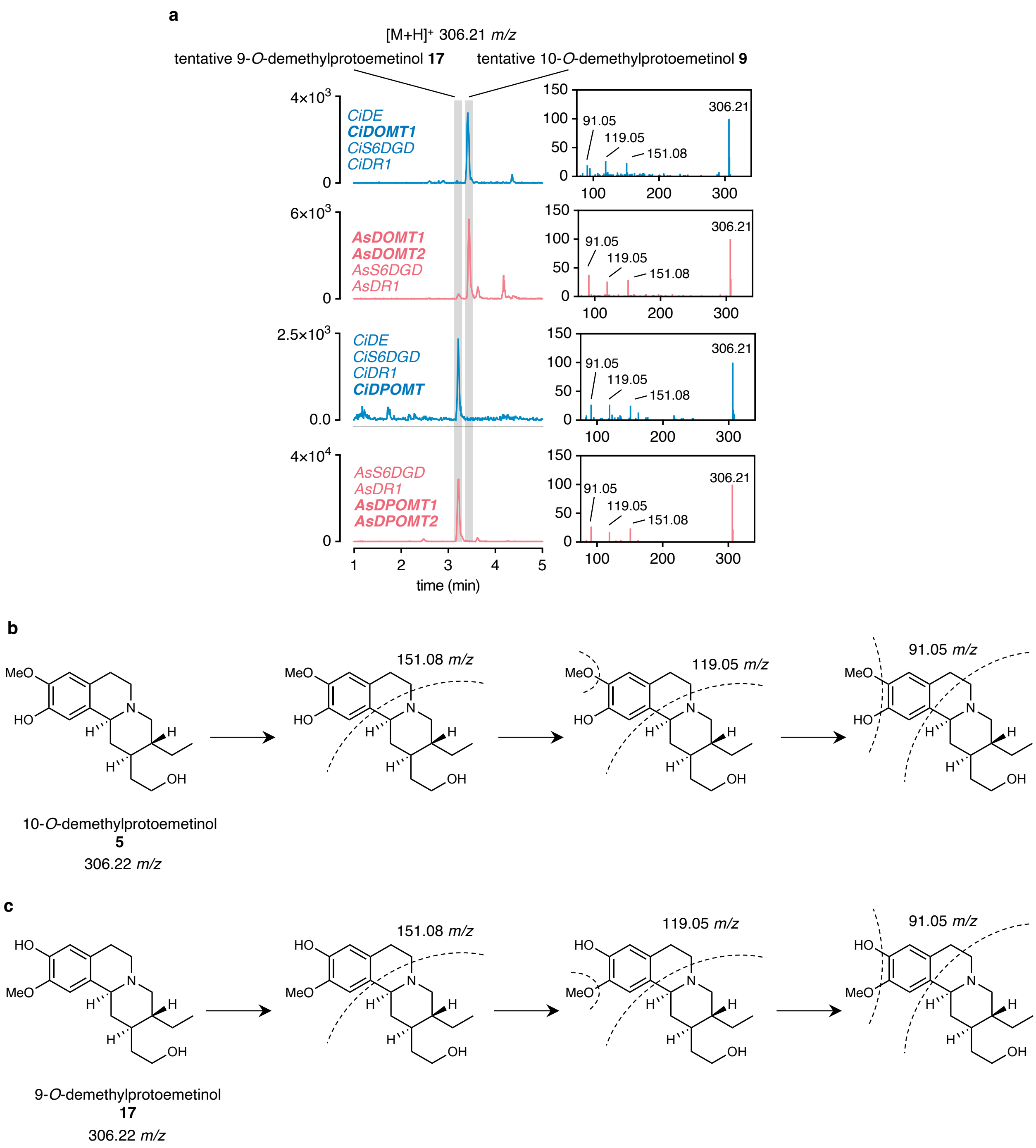

**Supplementary Fig. 15. Assignment of 10-*O*-demethylprotoemetinol and 9-*O*-demethylprotoemetinol. a**, LC-MS traces and MS2 data extracted from pathway reconstitution experiments shown in Fig. 3 and Supplementary Fig. 10 (*C. ipecacuanha*) and 12 (*A. salviifolium*). Two peaks with different retention times but identical $MS^1$ *m/z* and $MS^2$ fragments are detected when indicated pathway genes are expressed in combination. **b**, proposed fragmentation of 10-*O*-demethylprotoemetinol. **c**, proposed fragmentation of 9-*O*-demethylprotoemetinol. The peaks are proposed to correspond to 9-*O*-demethylprotoemetinol and 10-*O*-demethylprotoemetinolrespectively, based on the following observations: (a) $MS^1$ data are consistent with the theoretical *m/z*, (b) the observed $MS^2$ fragments are in agreement with predicted fragments, (c) $MS^2$ fragments show expected difference in *m/z* (-14) compared to protoemetinol fragments (*m/z* 165.09 in protoemetinol versus *m/z* 151.08 observed here), (d) different retention times depending on the specific combination of combinatorically expressed pathway genes are consistently observed and identical for both species. Data for *C. ipecacuanha* pathway gene expression is shown in blue and labelled with *C.i.* as abbreviation; data for *A. salviifolium* pathway gene expression in magenta, abbreviated with *A.s.*; and data for the respective standards in black as indicated. $MS^2$ data is shown as relative abundance of ions with *m/z* values of the most abundant fragment ions indicated. The specific pathway genes expressed were as indicated. In bold the pathway genes that differed in the combinations leading to peaks with different retention times.

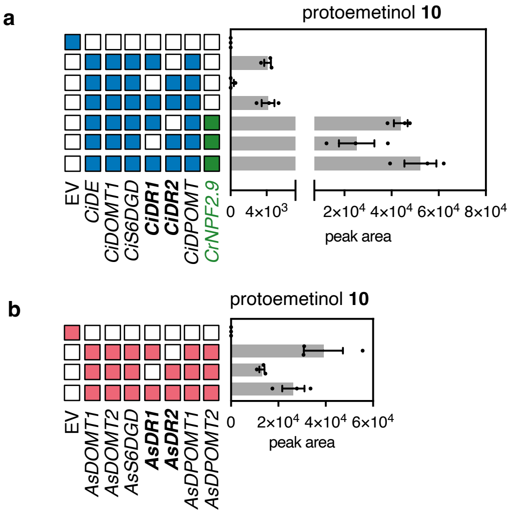

**Supplementary Fig. 16. Additional CiDR and AsDR paralogs have partial redundancy with DR1s. a,** *CiDR2* is a coexpressed *MDR* transcript with high sequence homology to *CiDR1* (see Supplementary Fig. 8a) and reconstitutes the pathway to protoemetinol, if the heterologous vacuolar exporter *CrNPF2.9* is co-overexpressed. However, in direct comparison to expression of *CiDR1*, less protoemetinol accumulates. **b**, *AsDR2* is a coexpressed *MDR* transcript with high sequence homology to *AsDR1* (see Supplementary Fig. 8b) and reconstitutes the pathway to protoemetinol, although less protoemetinol accumulates in direct comparison to expression of *AsDR1*. LC-MS peak areas are shown as bars of the mean of three biological replicates, error bars are standard error of the mean.

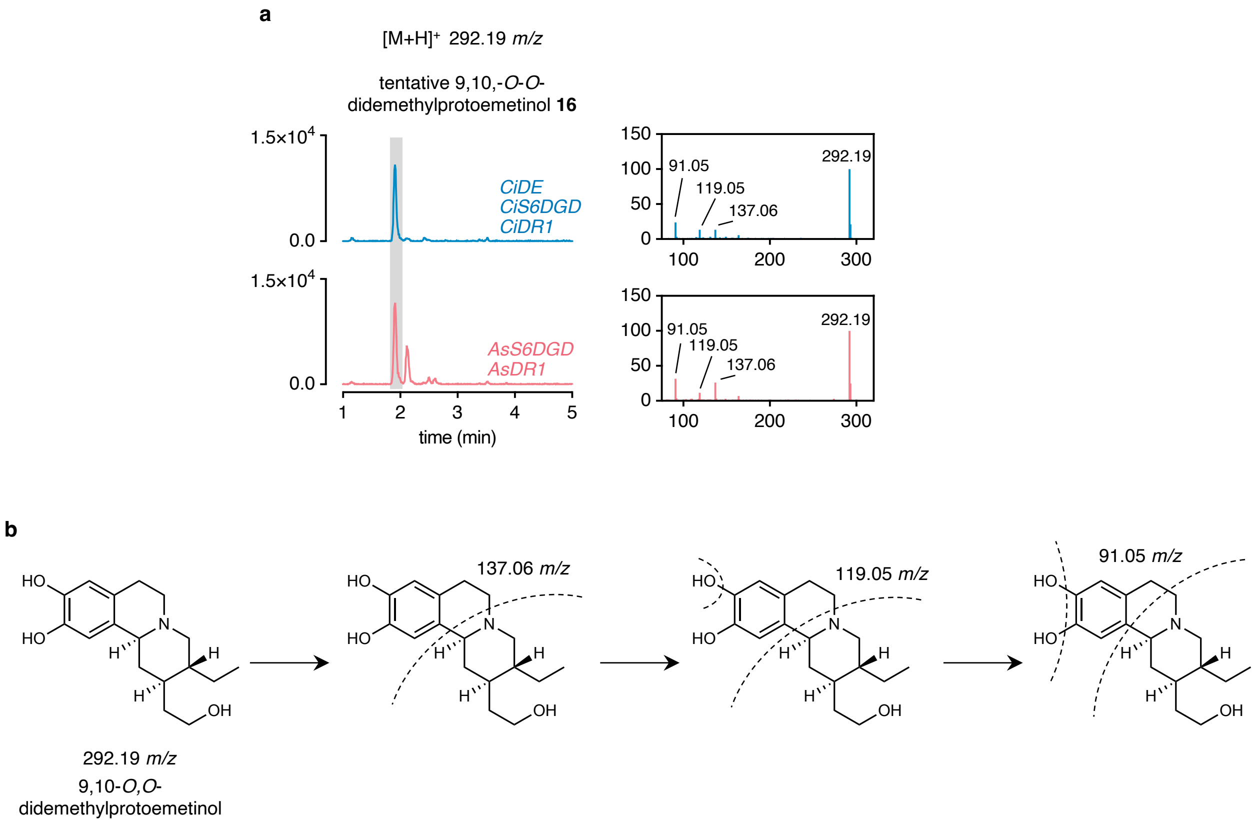

**Supplementary Fig. 17. Assignment of 9,10-*O*,*O*-didemethylprotoemetinol. a**, LC-MS traces and MS$^2$ data extracted from pathway reconstitution experiments shown in Fig. 3 and Supplementary Fig. 10 (*C. ipecacuanha*) and 12 (*A. salviifolium*). Upon expression of the indicated pathway genes, one peak with the same retention time and identical MS$^1$ *m/z* and MS$^2$ fragments was detected. Upon expression of *A. salviifolium* pathway genes only, an additional peak with different retention time but same *m/z* appeared which may be a different stereoisomer. **b**, fragmentation of 9,10-*O,O*-didemethylprotoemetinol. This peak is proposed to be 9,10-*O,O*-didemethylprotoemetinol based on the following observations: (a) MS$^1$ data are consistent with the theoretical *m/z*, (b) the observed MS$^2$ fragments are in agreement with predicted fragments, (c) MS$^2$ fragments show expected difference (-28 and -14, respectively) in *m/z* compared to protoemetinol and proposed 9- and 10-*O*-demethylprotoemetinol fragments (*m/z* 165.09 in protoemetinol versus *m/z* 137.06 observed here), (d) appearance of peaks in specific combinations of expressed pathway genes is as expected. *C. ipecacuanha* data is shown in blue and labelled with *C.i.* as abbreviation; *A. salviifolium* data in magenta, abbreviated with *A.s.*; and data for the respective standards in black as indicated. MS$^2$ data is shown as relative abundance of ions with *m/z* values of the most abundant fragment ions indicated.

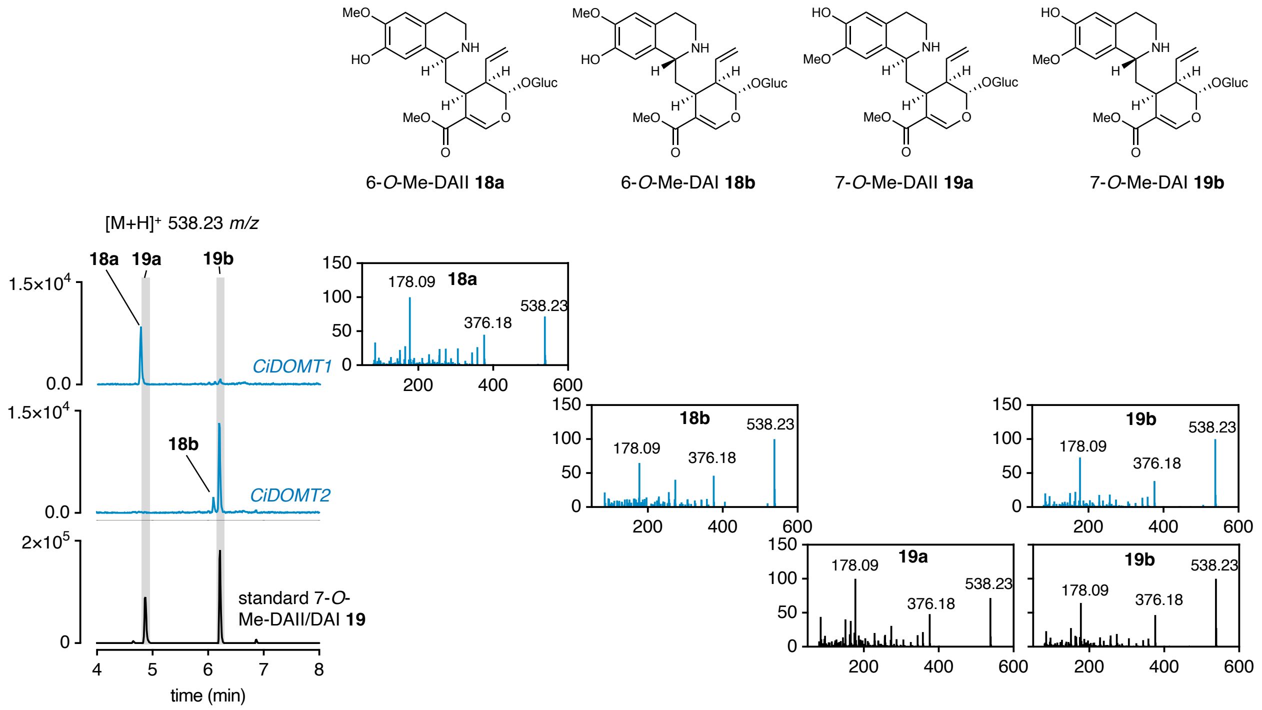

**Supplementary Fig. 18. Extracted ion chromatograms and MS$^2$ data of 7-*O*-Me-DAI/I and putative 6-*O*-Me-DAI/I from N. benthamiana agroinfiltrations and standards.** On the left EICs for 538.23 *m/z* of *N. benthamiana* expressing *CiDOMT1* (top), *CiDOMT2* (center) compared to 7-*O*-Me-DAI/I standard (bottom, black, produced through chemical Pictet-Spengler reaction of 4-*O*-Me-dopamine with secologanin). Based on results from derivatives and the observed ratio of peak intensity in the standard, it can be assumed that the first peak is the *S* epimer and the second one the *R* epimer. 6-*O*-Me-DAII/I are tentatively identified based on MS$^2$ fragmentation and peak shifts. Minor peaks in the standard with the same *m/z* values are consistently observed when performing these reactions and could correspond to “neo” isomers which were previously described (Beke *et al.*, 2001).

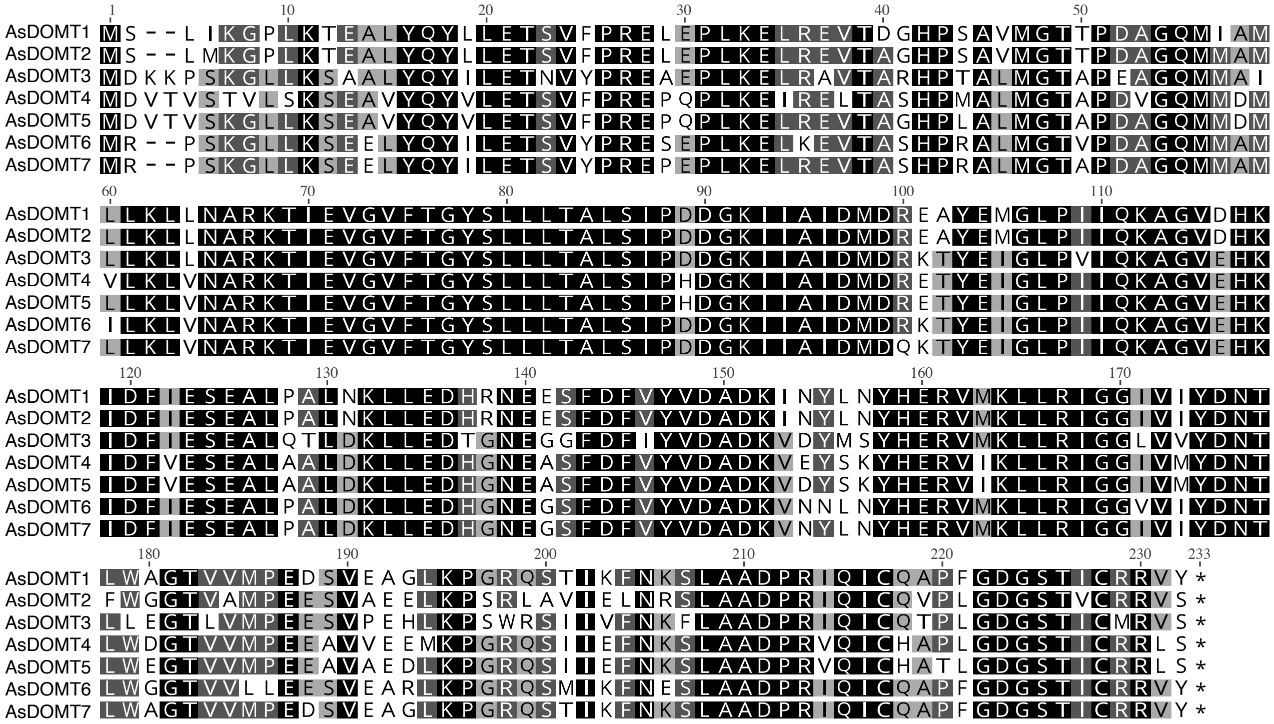

**Supplementary Fig. 19. AsDOMT amino acid alignment.** MUSCLE alignment. Colour code depicts identical amino acid residues in 100 % of the sequences (black), 80-100% (dark grey), 60-80% (light grey), and less than 60 % (white).

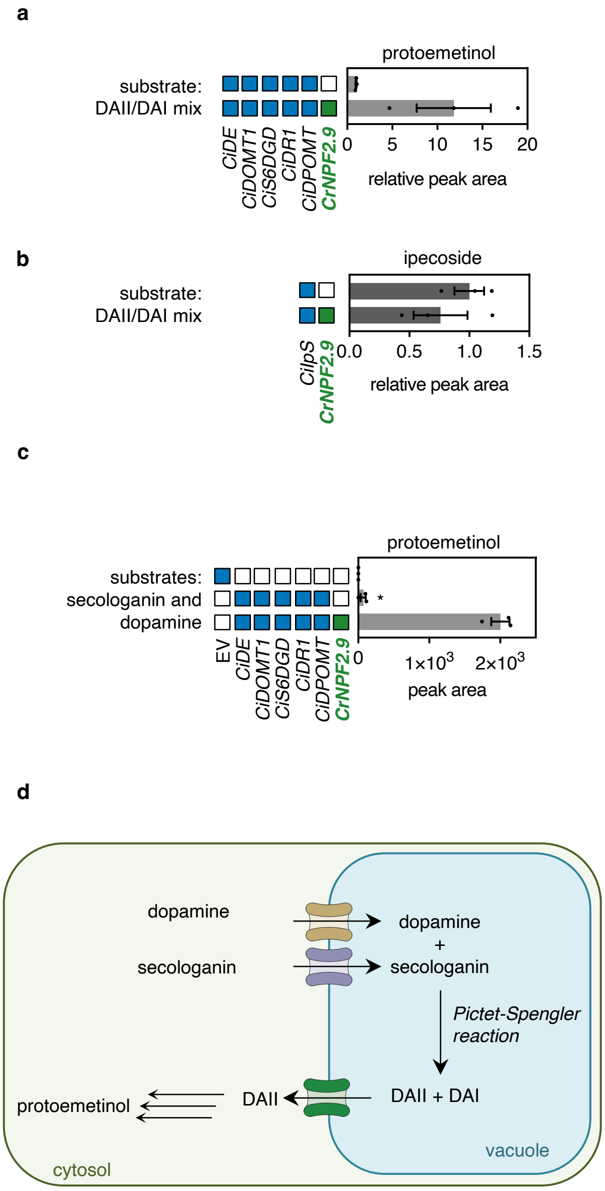

**Supplementary Fig. 20. Expression of the *C. roseus* strictosidine exporter gene *CrNPF2.9* boosts *C. ipecacuanha* protoemetine biosynthesis and enables biosynthesis from dopamine and secologanin directly. a**, comparison of protoemetinol accumulation upon expression of indicated *C. ipecacuanha* biosynthesis genes in *N. benthamiana* with or without co-expression of transporter *CrNPF2.9*. Exogenous DAI/DAII are infiltrated as substrate and allowed to react for 24 hours before the leaf was harvested. Data was extracted from experiment shown in Supplementary Fig. 10. Peak areas are shown here relative to areas from samples where *CrNPF2.9* was not expressed as bars of the mean of three replicates. **b**, expression of *CiIps* with or without co-overexpression of *CrNPF2.9* did not lead to enhanced accumulation of the *R*-epimer derivative ipecoside. **c**, when *C. ipecacuanha* protoemetine pathway genes are expressed and uncoupled secologanin and dopamine are fed for 48 hours, only traces of protoemetinol can be detected (indicated with an asterisk). When *CrNPF2.9* is co-overexpressed protoemetinol is formed to detectable levels. **d**, it is hypothesized that spontaneous *in planta* coupling occurs in the vacuole and the resulting formed DAII is only accessible to the cytosolic pathway enzymes if it is exported by a vacuolar exporter. CrNPF2.9 appears to export DAII. These results suggest the presence of a yet to be identified vacuolar exporter in *C. ipecacuanha* and *A. salviifolium*. LC-MS peak areas are shown as bars of the mean of three biological replicates, error bars are standard error of the mean.

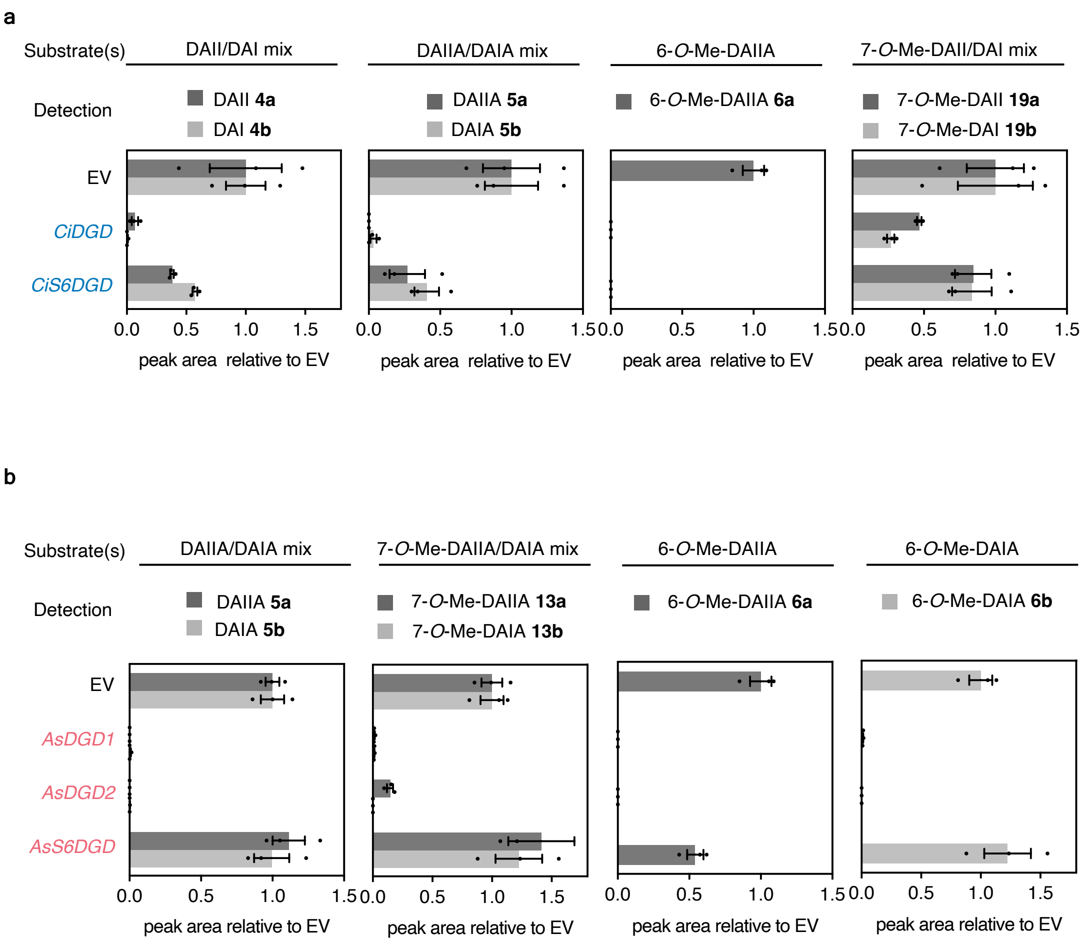

**Supplementary Fig. 21. Feeding ipecac alkaloid glucosides to leaf disks expressing *DGDs* or *S6DGDs*.** Consumption of glucosides was measured because the unstable aglycone products cannot be reliably detected. **a**, agroinfiltration of *N. benthamiana* with strains harbouring constructs for overexpression of *CiDGD* or *Ci6SDGD* or empty vector (EV) respectively. Leaf disks were cut and incubated separately with indicated substrates. Note that 7-*O*-Me-DAIA and 7-*O*-Me-DAIIA are not detected in native *C. ipecacuanha* plants. **b**, agroinfiltration of *N. benthamiana* with strains harbouring constructs for overexpression of *AsDGD1* or *2* or *As6S6DGD* or EV respectively. Leaf disks were cut out and incubated each with indicated substrates. AsDGDs deglycosylate all derivatives whereas AsS6DGD only shows activity towards 6-*O*-Me-DAIIA. 6-*O*-Me-DAIIA and 6-*O*-Me-DAIA cannot be obtained readily by Pictet-Spengler chemical reaction and were thus produced *in vitro* by recombinant CiDOMT1 and AsDOMT3, respectively (see methods). Bars are mean peak areas of three biological replicates relative to peak areas measured in EV, error bars are standard error of the mean SEM.

| Protein | Localizations | Cytoplasm | Nucleus | Extra cellular | Cell membrane | Mitochondrion | Plastid | Endoplasmic reticulum | Lysosome/ Vacuole | Golgi apparatus | Peroxisome | Peripheral | Trans membrane | Lipid anchor |
|---|---|---|---|---|---|---|---|---|---|---|---|---|---|---|
| CiDGD | **Nucleus** | 0.277 | **0.804** | 0.338 | 0.063 | 0.102 | 0.009 | 0.110 | 0.160 | 0.097 | 0.302 | 0.199 | 0.130 | 0.202 |
| CiS6DGD | **Nucleus** | 0.253 | **0.795** | 0.227 | 0.049 | 0.124 | 0.055 | 0.102 | 0.070 | 0.074 | 0.595 | 0.206 | 0.149 | 0.209 |
| AsDGD1 | **Nucleus** | 0.308 | **0.730** | 0.277 | 0.191 | 0.092 | 0.083 | 0.148 | 0.229 | 0.092 | 0.245 | 0.230 | 0.119 | 0.176 |
| AsDGD2 | **Nucleus** | 0.333 | **0.779** | 0.248 | 0.192 | 0.075 | 0.061 | 0.112 | 0.236 | 0.081 | 0.171 | 0.209 | 0.132 | 0.169 |
| AsS6DGD | **Cytoplasm** | **0.764** | 0.314 | 0.195 | 0.213 | 0.048 | 0.193 | 0.284 | 0.325 | 0.102 | 0.449 | 0.251 | 0.087 | 0.180 |

**Supplementary Fig. 22. Prediction of nuclear localization signal peptide by DeepLOC2.** The GH-1 type glucosidases AsDGD1 and 2 as well as CiS6DGD and CiDGD are all predicted to localize to the nucleus (and contain a bipartite NLS, not shown here) whereas the GH-3 type AsS6DGD is predicted to be cytosolic.

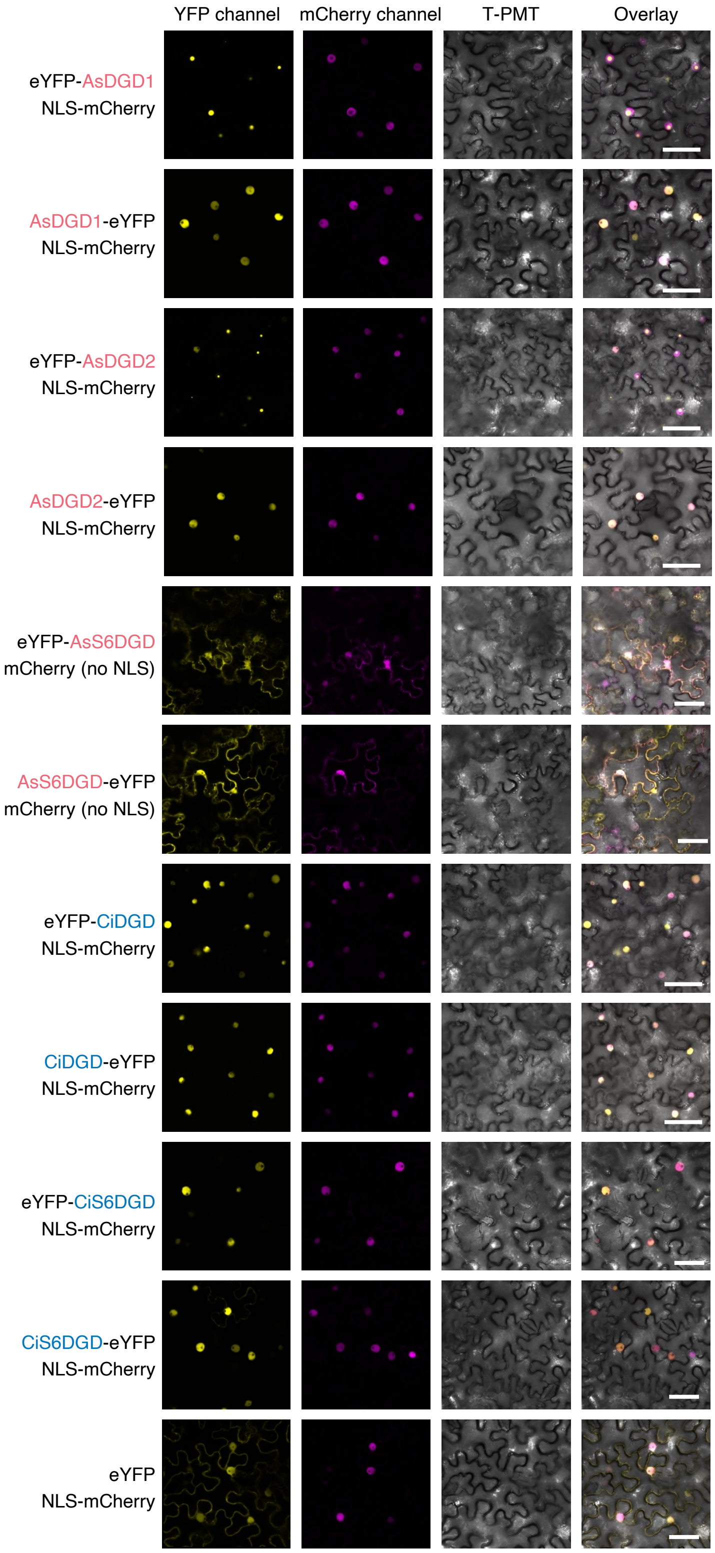

**Supplementary Fig. 23. Confocal laser scanning microscopy of glucosidases fused to eYFP.** AsDGD1,2 localize to the nucleus as *N*-terminal or *C*-terminal fusion proteins (row 1-4). When eYFP is fused to the *N*-terminus of AsDGDs the signal appears as a small particle within the nucleus (row 1 and 3 compared to 2 and 4). AsS6DGD shows localization signals in both nucleus and cytosol. Signals of CiDGD and CiS6DGD appear in the nucleus and appear similar when eYFP is fused to either the *N*- or *C*-terminus (row 7-10). mCherry is used either fused to NLS as nucleus marker or without tag as a marker for both cytosol and nucleus (only in combination with AsS6DGD). Scale bars indicate 50 μm.

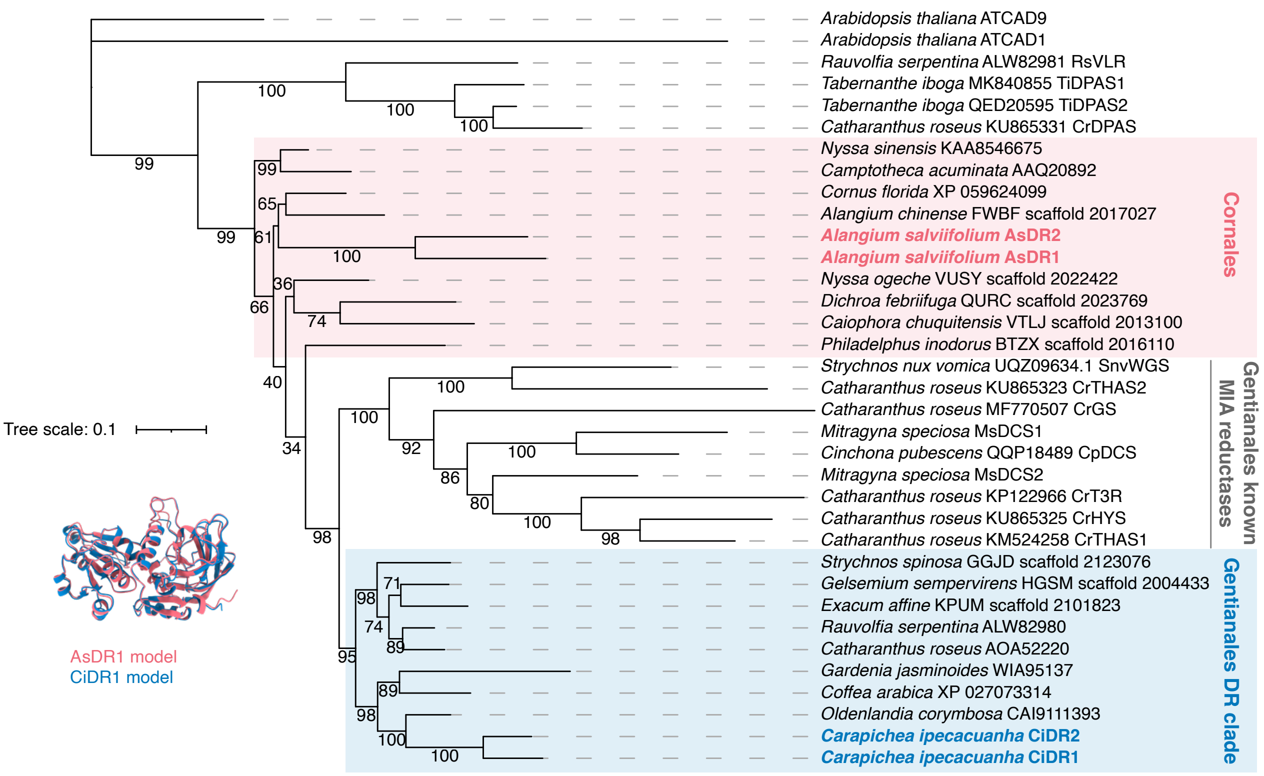

**Supplementary Fig. 24. Parallel evolution of *A. salviifolium* versus *C. ipecacuanha* DRs.** Maximum likelihood phylogenetic tree of DR amino acid sequences and homologs from other Cornales and Gentianales species. Alphafold3 models indicate same folds for all DRs.

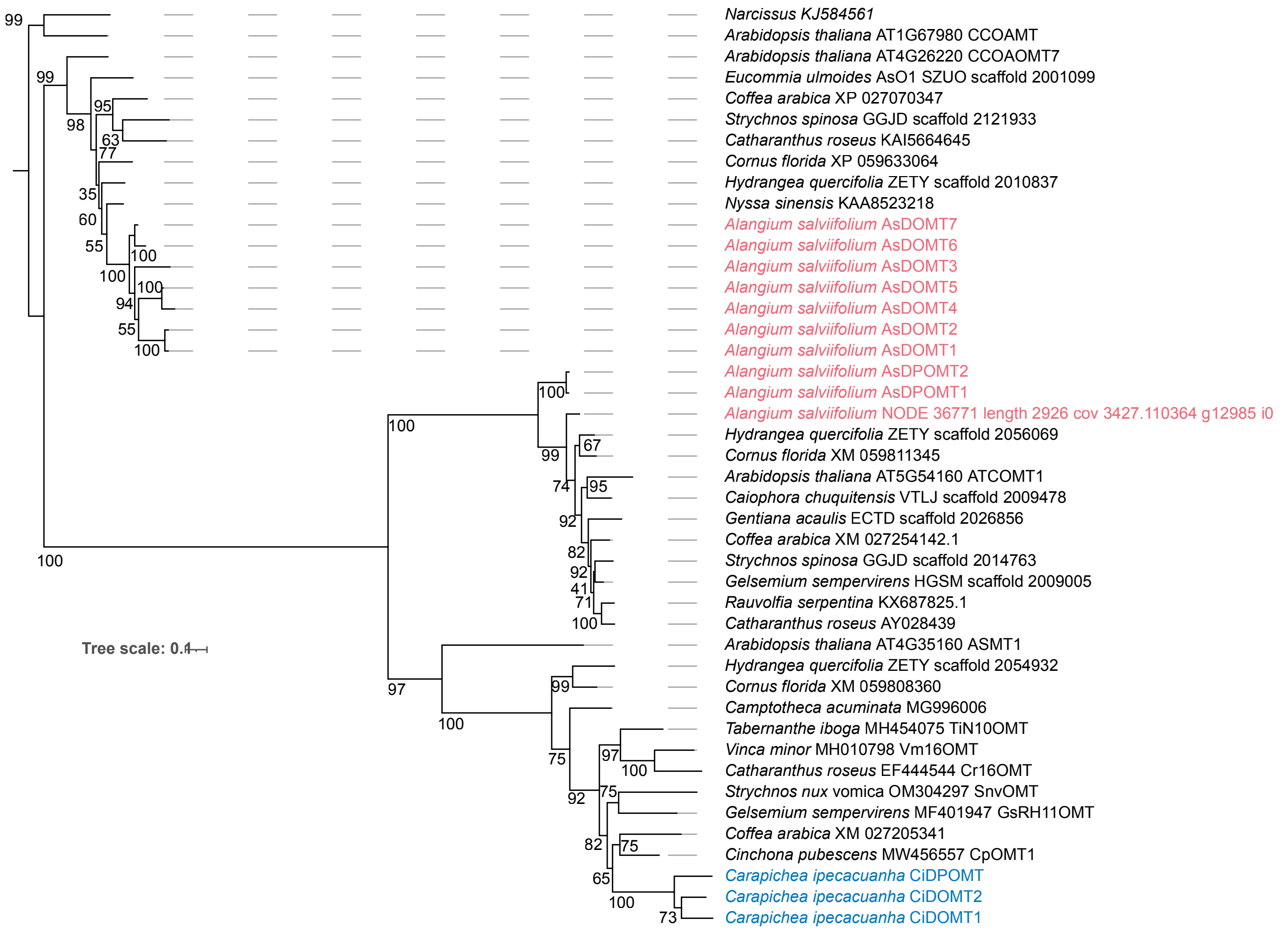

**Supplementary Fig. 25. Maximum likelihood tree of *A. salviifolium* versus *C. ipecacuanha* OMTs.** Tree from Fig. 6a shown here with all bootstrap values.

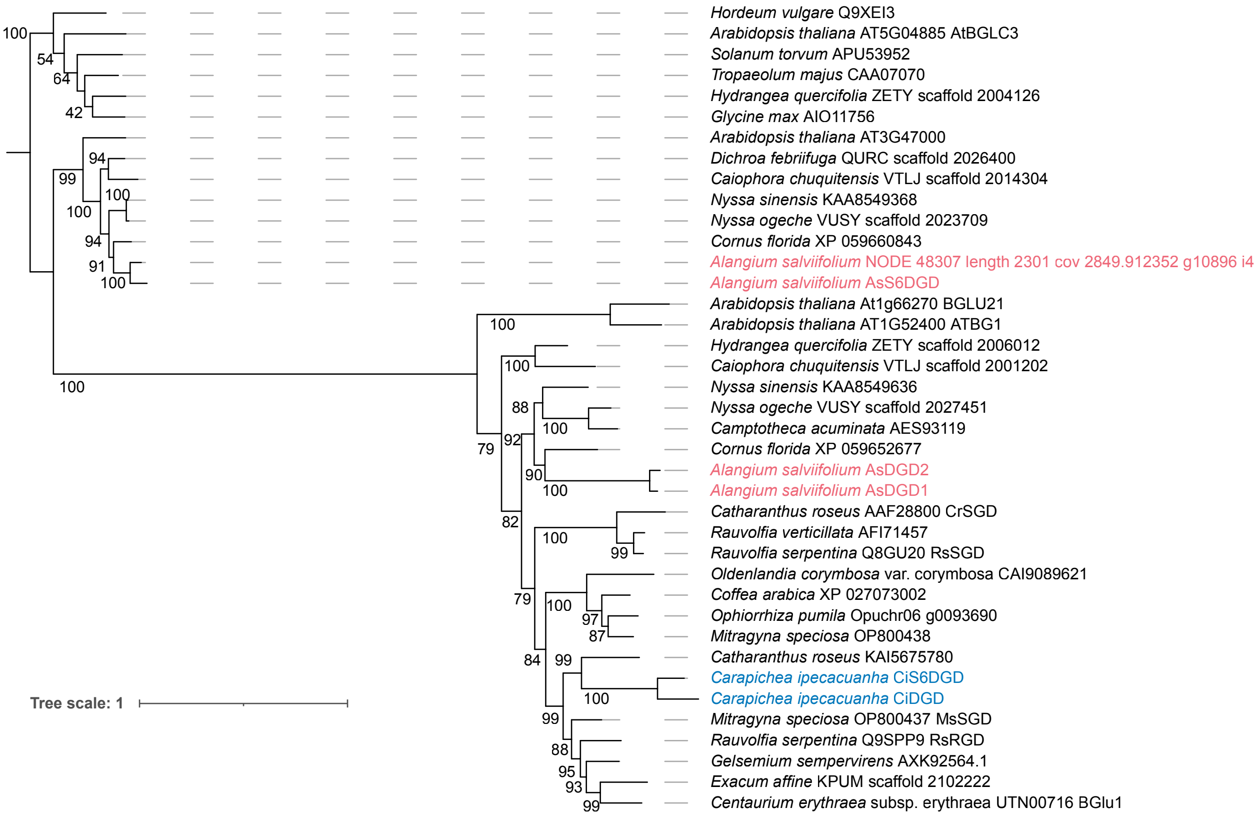

**Supplementary Fig. 26. Maximum likelihood tree of *A. salviifolium* versus *C. ipecacuanha* glucosidases.** Tree from Fig. 6b shown here with all bootstrap values

#### Supplementary Fig. 27. NMR data for synthetic protoemetine

500 MHz NMR in $CDCl_3$

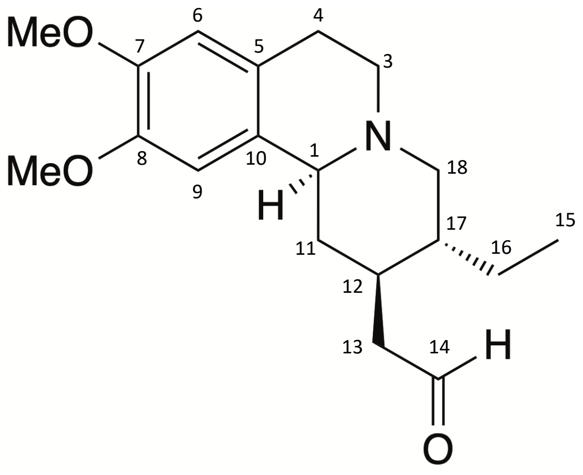

| pos. | $\delta_H$ | mult. | $J_{HH}$ | $\delta_C$ |
|---|---|---|---|---|
| 1α | 3.12 | *bd* | 11.8 | 62.6 |
| 3α | 2.97 | *ddd* | 11.2/5.9/1.4 | 52.6 |
| 3β | 2.50 | *ddd* | 11.5/11.2/4.0 | 52.6 |
| 4α | 2.63 | *m* | - | 29.3 |
| 4β | 3.10 | *m* | - | 29.3 |
| 5 | - | - | - | 126.8 |
| 6 | 6.56 | *s* | - | 111.6 |
| 7 | - | - | - | 147.6 |
| 8 | - | - | - | 147.3 |
| 9 | 6.63 | *s* | - | 108.2 |
| 10 | - | - | - | 129.7 |
| 11α | 2.33 | *m* | - | 38.3 |
| 11β | 1.28 | *ddd* | 12.3/11.8/11.8 | 38.3 |
| 12α | 1.95 | *dddd* | 15.4/12.3/8.3/3.8 | 36.0 |
| 13a | 2.72 | *ddd* | 17.1/3.8/1.3 | 48.3 |
| 13b | 2.33 | *ddd* | 17.1/8.3/2.3 | 48.3 |
| 14 | 9.87 | *dd* | 2.3/1.3 | 202.7 |
| 15 | 6.85 | *ddd* | 7.4/7.4/0.9 | 120.8 |
| 16a | 1.58 | *m* | - | 23.9 |
| 16b | 1.12 | *m* | - | 23.9 |
| 17 | 1.48 | *m* | - | 41.5 |
| 18α | 2.07 | *dd* | 11.4/11.2 | 61.2 |
| 18β | 3.09 | *dd* | 11.4/4.0 | 61.2 |
| 7-OMe | 3.84 | *s* | - | 56.0 |
| 8-OMe | 3.83 | *s* | - | 56.2 |

The chemical shifts agreed with the published data (Lin *et al.*, 2011).

**Supplementary Fig. 27** NMR spectra for protoemetine

$^1$H NMR full range in CDCl$_3$

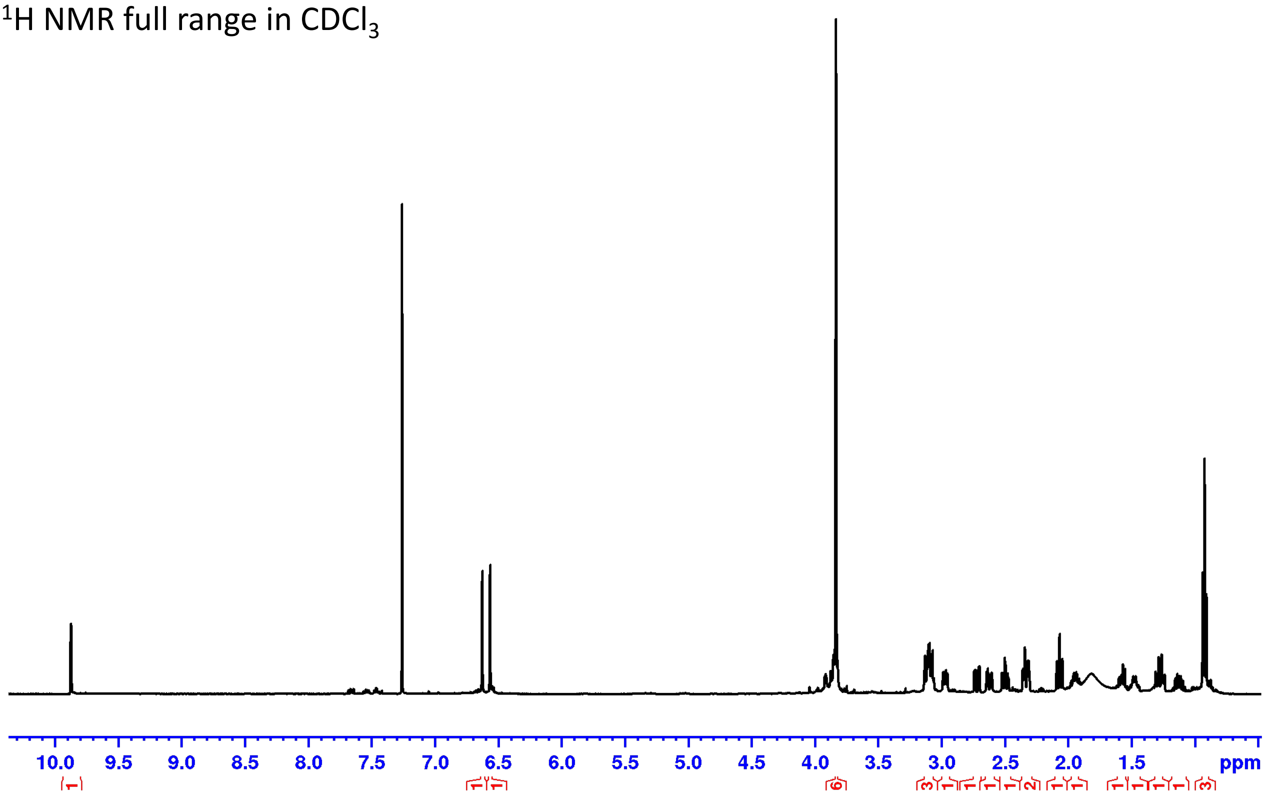

DEPTQ full range in CDCl$_3$

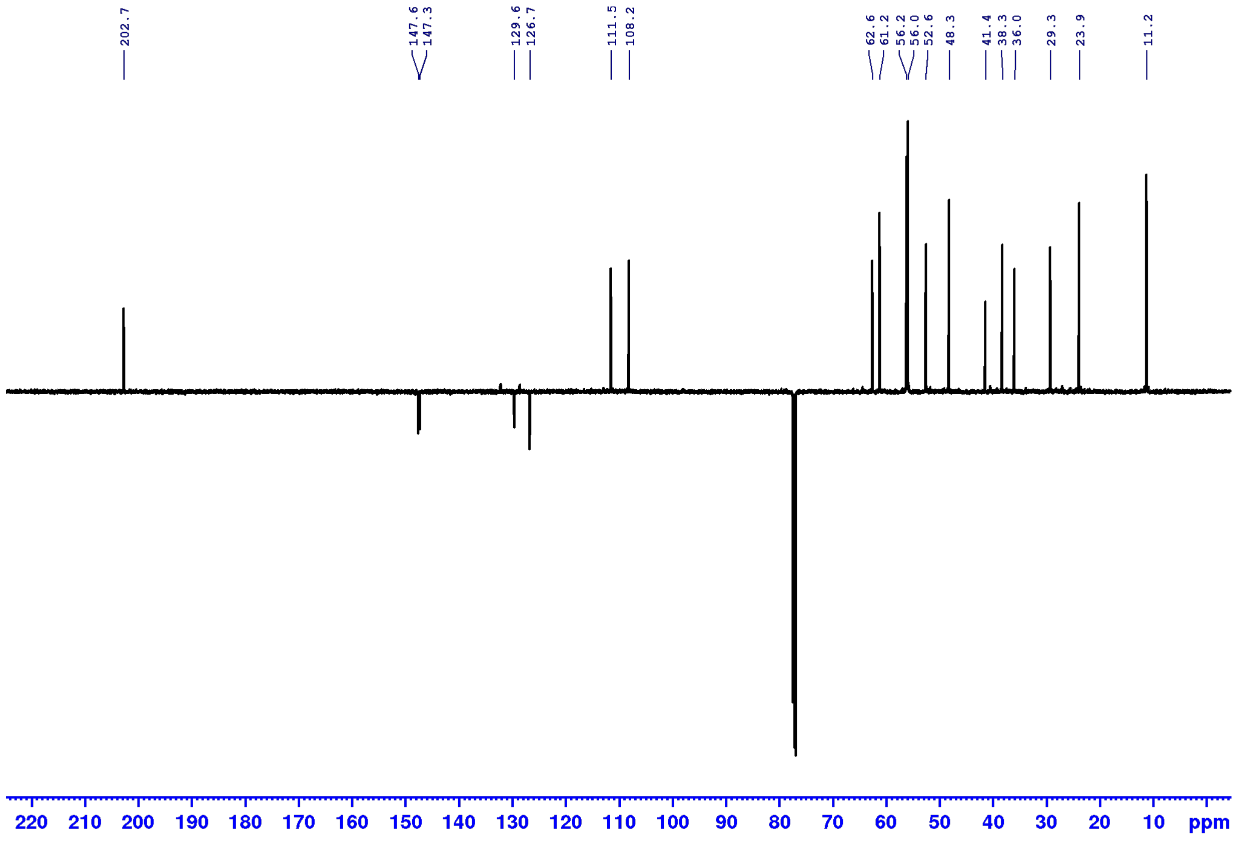

#### Supplementary Fig. 27 NMR spectra for protoemetine

Phase sensitive HSQC, full range in $CDCl_3$

red: $CH_2$, black: CH, $CH_3$

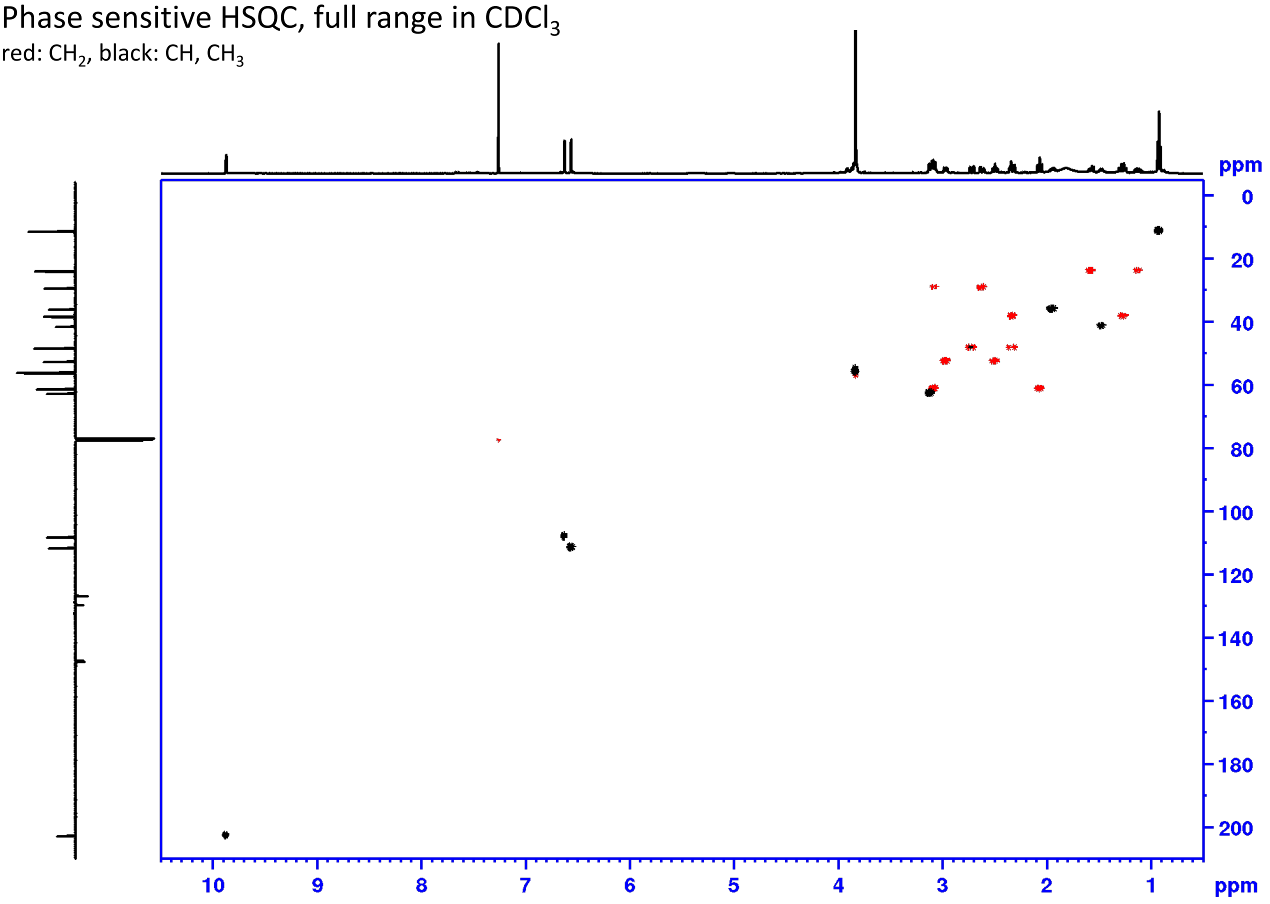

HMBC, full range in $CDCl_3$

Shaded areas mark impurity and solvent

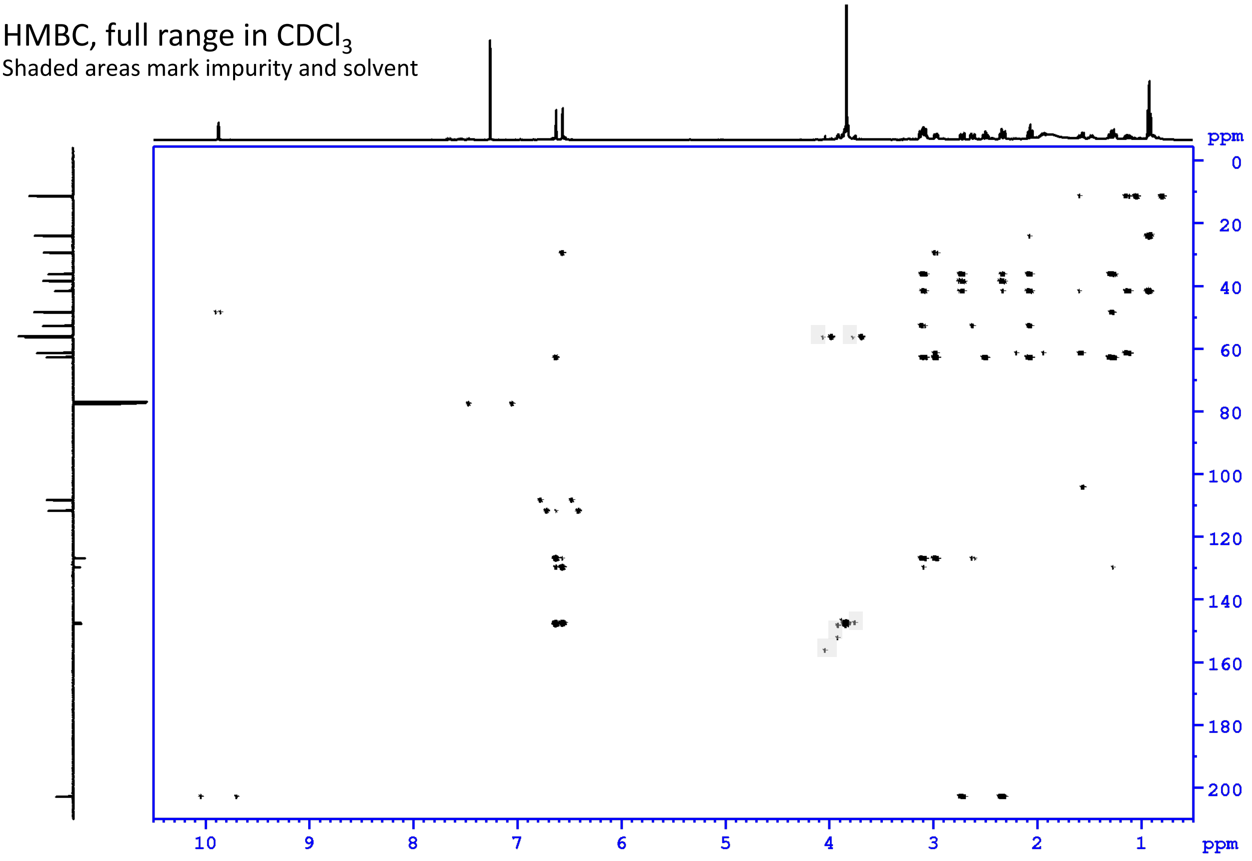

**Supplementary Fig. 27 NMR spectra for protoemetine**

COSY, full range in $CDCl_3$

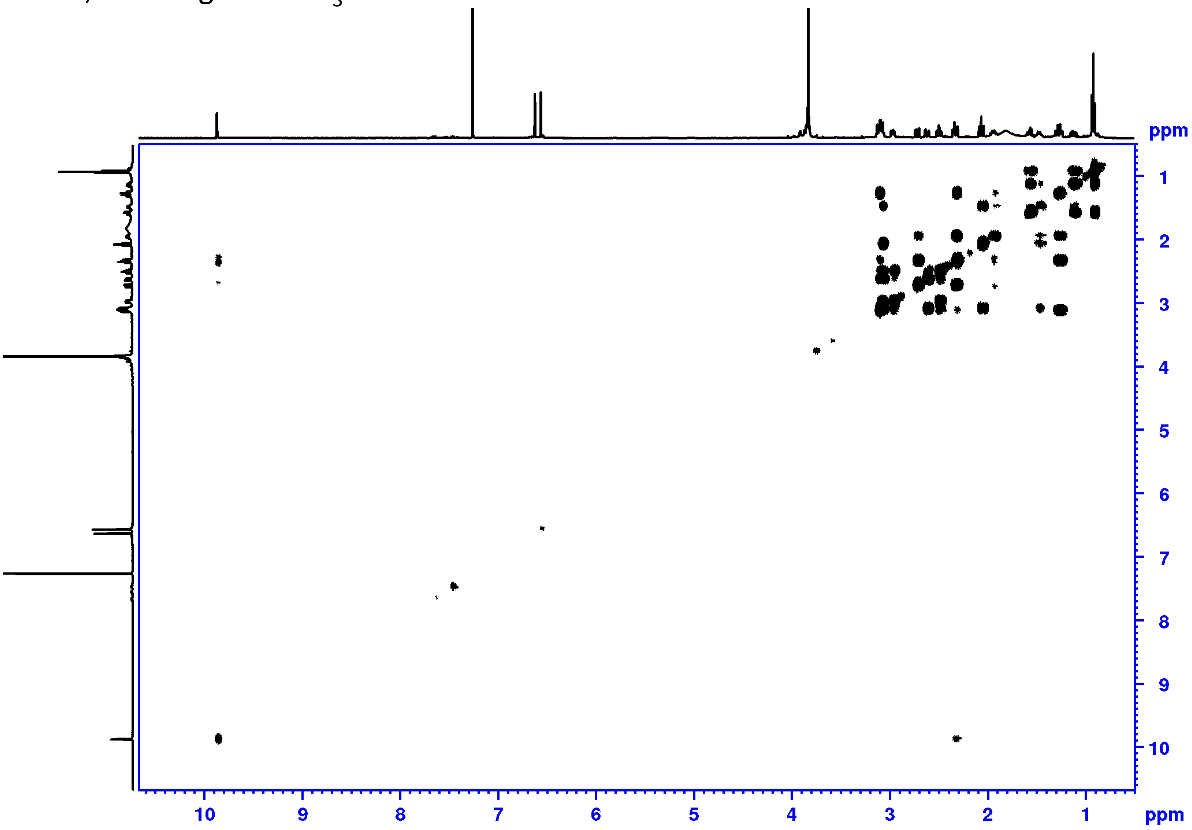

ROESY full range in $CDCl_3$

Shaded areas mark impurity and solvent

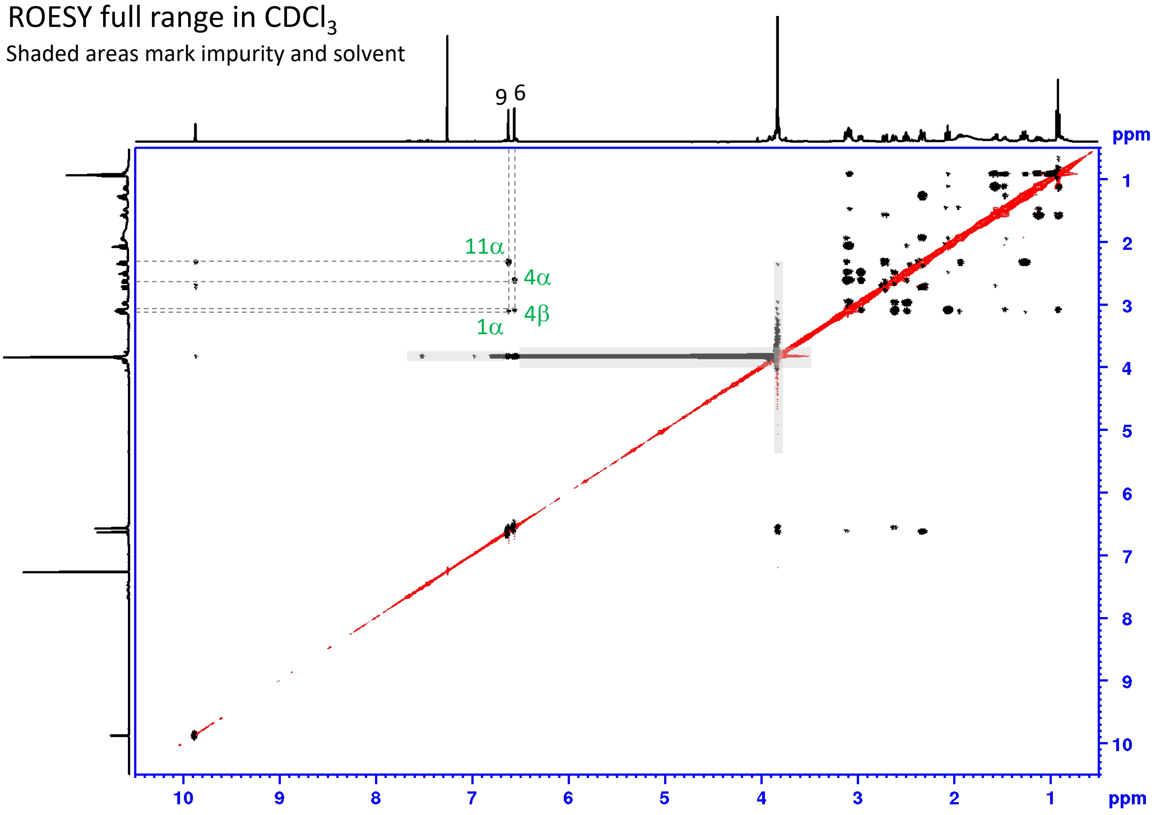

Important ROESY correlations are depicted in green.

#### Supplementary Fig. 27 NMR spectra for protoemetine

ROESY aliphatic range in $CDCl_3$

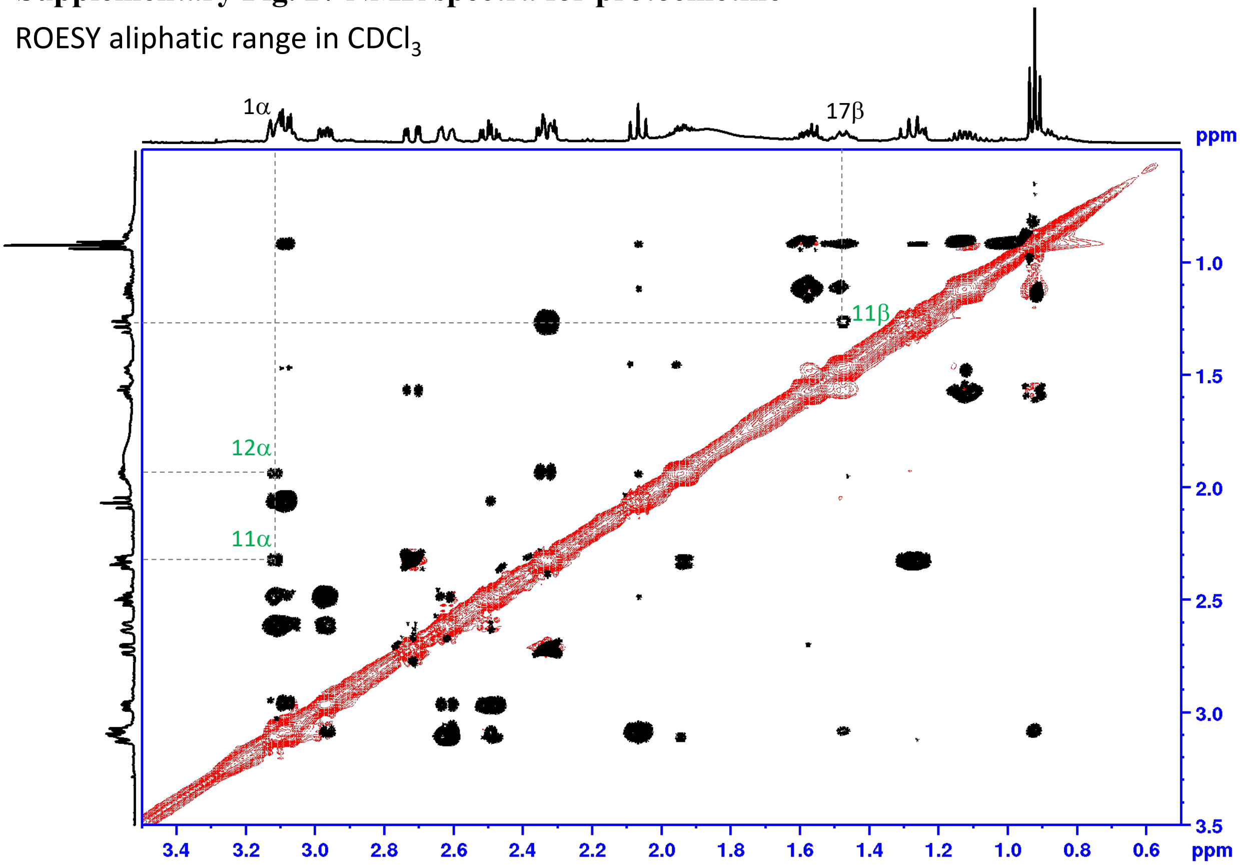

Important ROESY correlations are depicted in green.

#### Supplementary Fig. 27 ROESY correlations of protoemetine

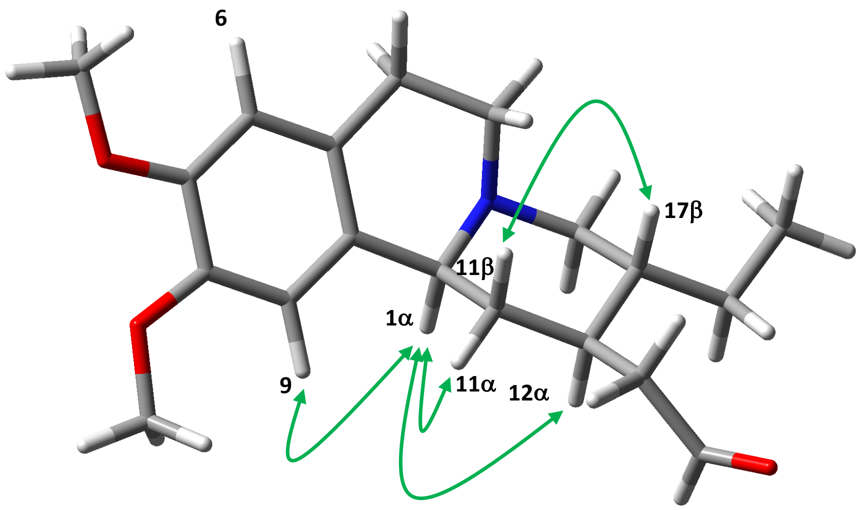

Optimized using Gaussian 16W (PM6, solvent $CDCl_3$).
Important ROESY correlations are depicted in green.

### Supplementary Fig. 28. NMR data for deacetylisoipecoside

500 MHz NMR in MeOH-$d_3$ + 0.1% formic acid

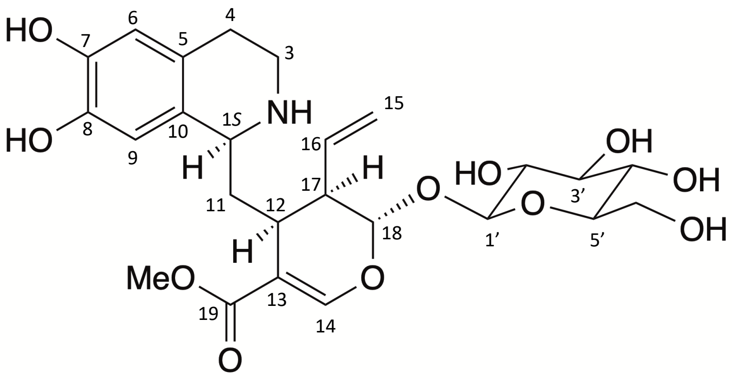

| pos. | $\delta_H$ | mult. | $J_{HH}$ | $\delta_C$ |
|---|---|---|---|---|
| 1α | 4.31 | *dd* | 11.1/3.2 | 54.4 |
| 3α | 3.35 | *m** | - | 39.9 |
| 3β | 3.41 | *m** | - | 39.9 |
| 4a | 2.95 | *m* | - | 25.5 |
| 4b | 2.95 | *m* | - | 25.5 |
| 5 | - | - | - | 123.2 |
| 6 | 6.58 | *s* | - | 116.1 |
| 7 | - | - | - | 146.9 |
| 8 | - | - | - | 146.0 |
| 9 | 6.56 | *s* | - | 114.1 |
| 10 | - | - | - | 124.2 |
| 11a | 1.99 | *ddd* | 14.8/11.7/3.2 | 36.5 |
| 11b | 2.18 | *ddd* | 14.8/11.1/3.5 | 36.5 |
| 12a | 2.98 | *m* | - | 32.1 |
| 13 | - | - | - | 109.3 |
| 14 | 7.74 | *s* | - | 156.2 |
| 15a | 5.34 | *ddd* | 17.4/1.2/1.2 | 119.7 |
| 15b | 5.28 | *ddd* | 10.5/1.2/1.1 | 119.7 |
| 16 | 5.84 | *ddd* | 17.4/10.5/7.7 | 135.6 |
| 17a | 2.69 | *m* | - | 45.3 |
| 18b | 5.74 | *d* | 8.5 | 97.3 |
| 19 | - | - | - | 170.7 |
| 19-OMe | 3.77 | *s* | - | 52.3 |
| 1' | 4.75 | *d* | 7.9 | 100.2 |
| 2' | 3.19 | *dd* | 9.3/7.9 | 74.7 |
| 3' | 3.38 | *dd* | 9.3/9.1 | 78.0 |
| 4' | 3.23 | *dd* | 9.7/9.1 | 71.6 |
| 5' | 3.32 | *m* | - | 78.5 |
| 6a' | 3.92 | *dd* | 11.8/2.1 | 62.8 |
| 6b' | 3.63 | *dd* | 11.8/6.7 | 62.8 |

* overlapped signals *J* unresolved

About position 11a to 18 b, please see the ROESY correlation figure.

**Supplementary Fig. 28 NMR spectra for deacetylisoipecoside**

presaturated $^1$H NMR full range in MeOH-$d_3$ + 0.1% formic acid (FA)

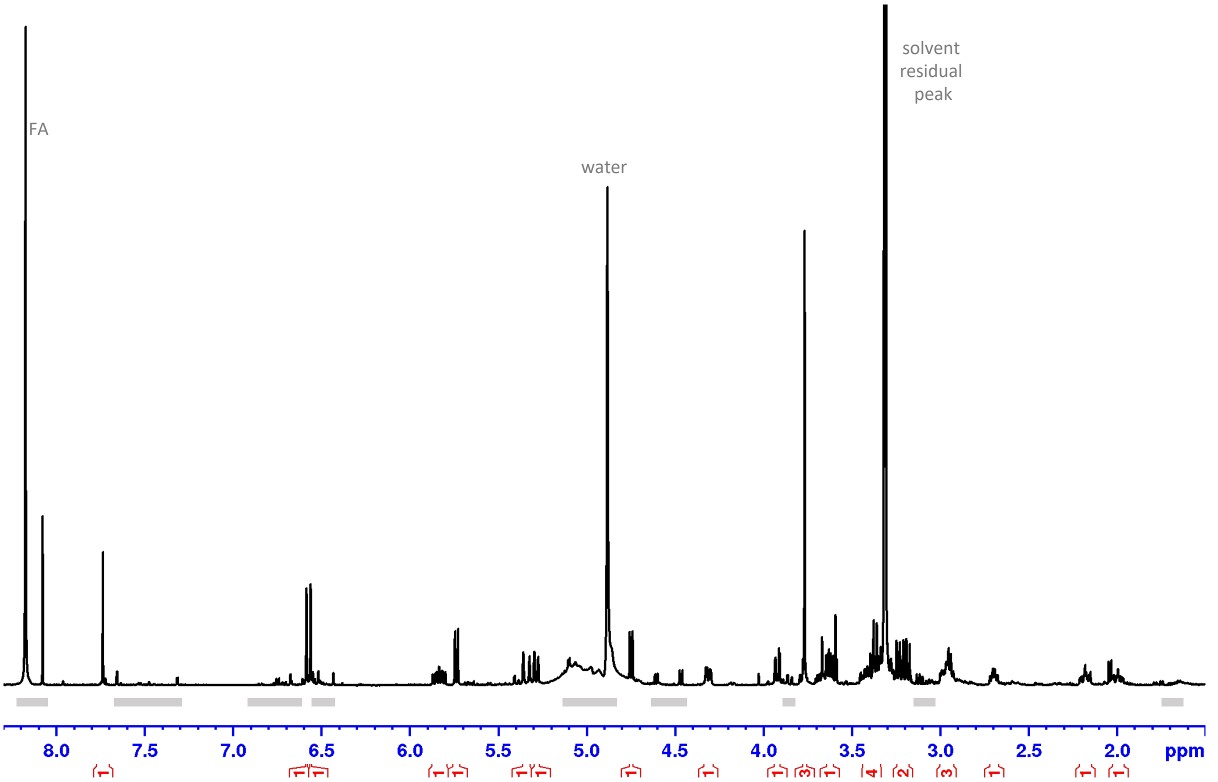

Shaded areas mark impurity and solvent

DEPTQ full range in MeOH-$d_3$ + 0.1% FA

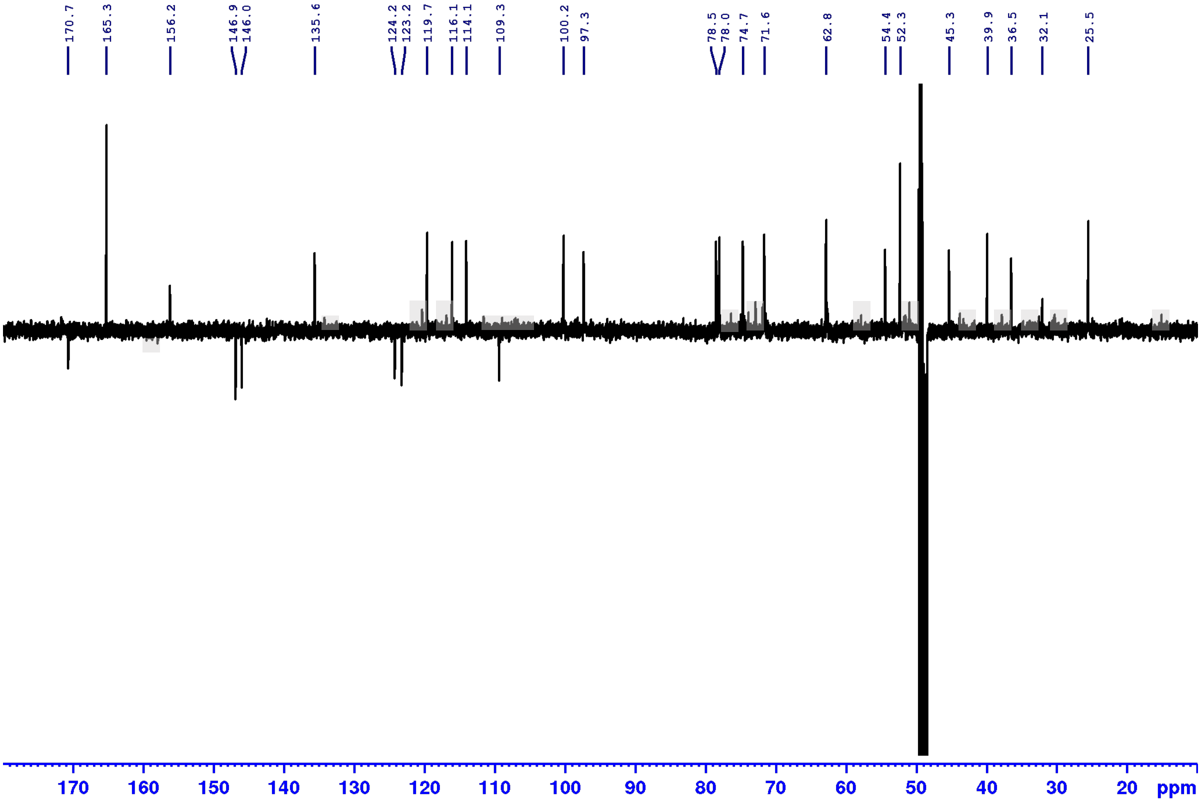

Shaded areas mark impurity and solvent

**Supplementary Fig. 28 NMR spectra for deacetylisoipecoside**

Phase sensitive HSQC, full range in MeOH-$d_3$ + 0.1% FA

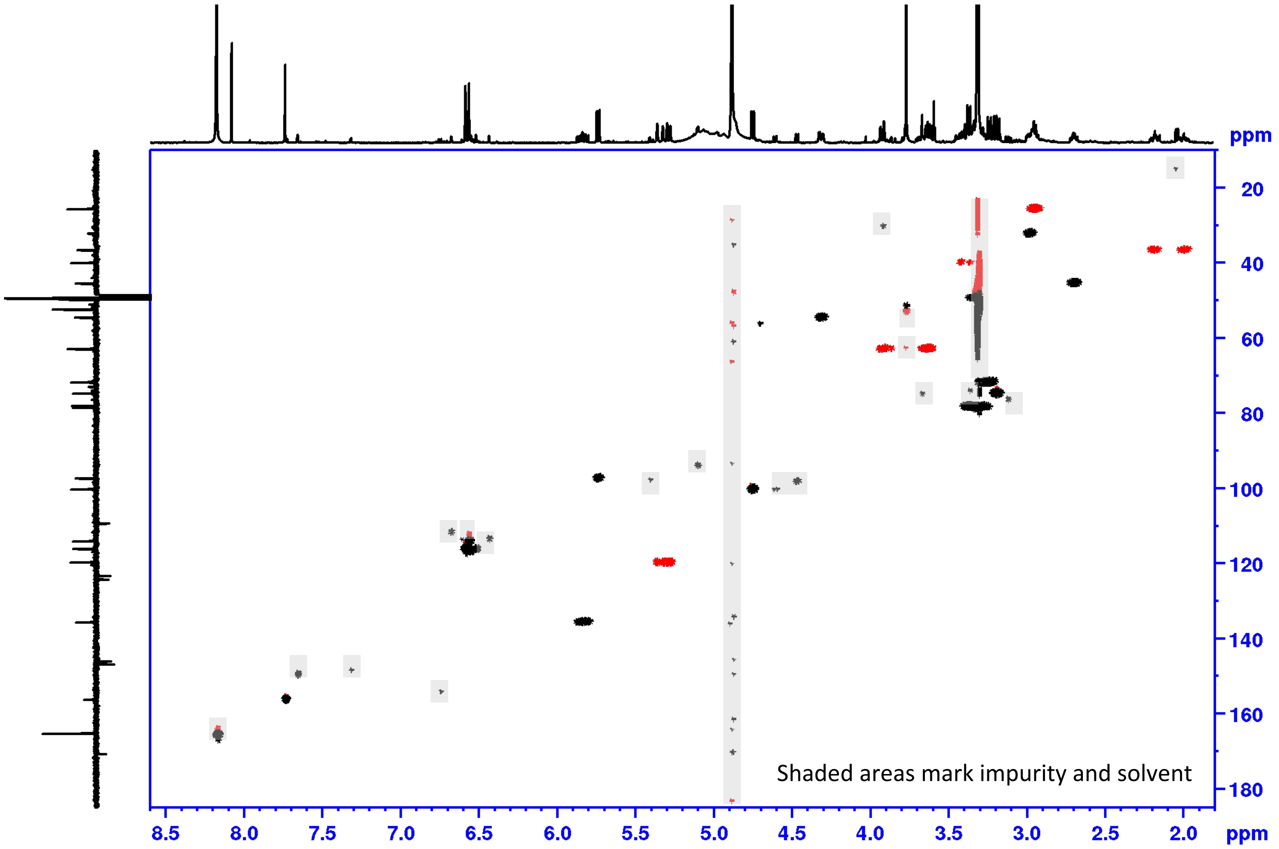

HMBC full range in MeOH-$d_3$ + 0.1% FA

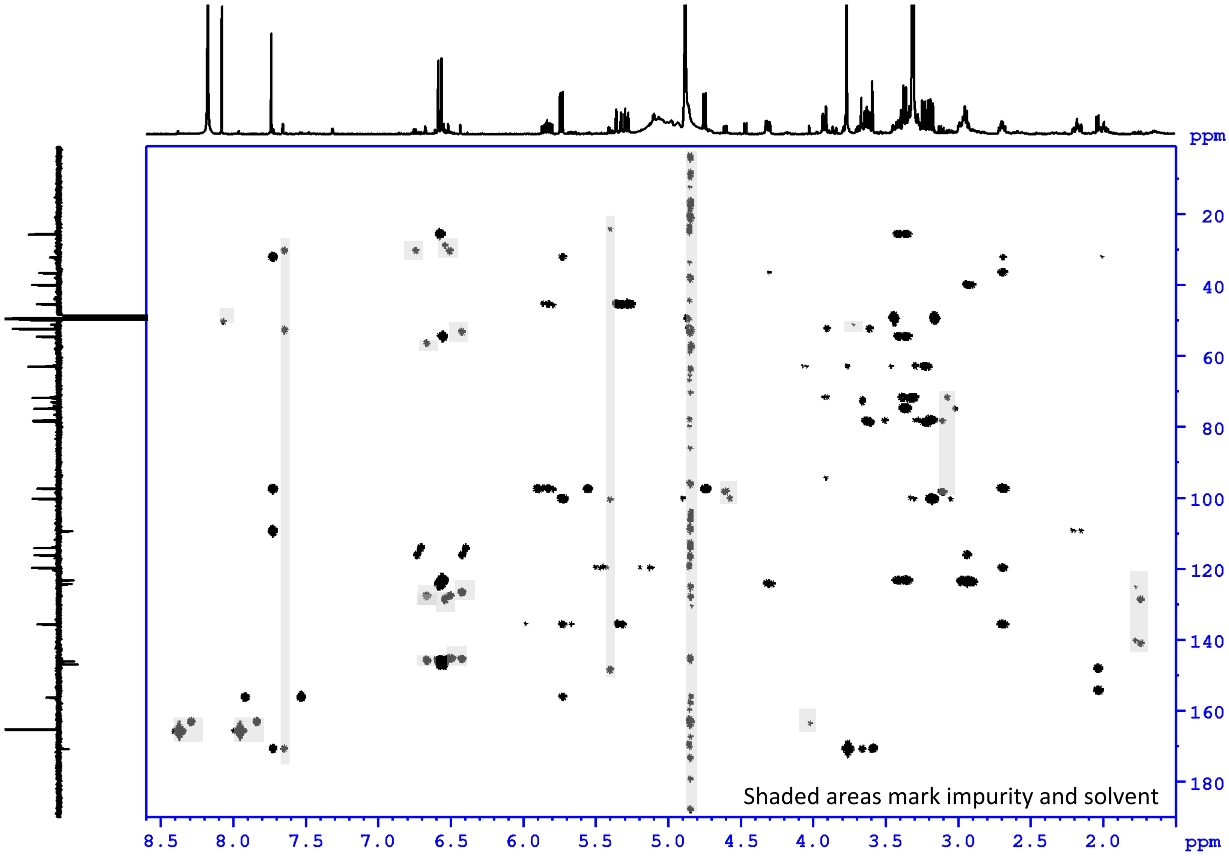

**Supplementary Fig. 28 NMR spectra for deacetylisoipecoside**

COSY, full range in MeOH-$d_3$ + 0.1% FA

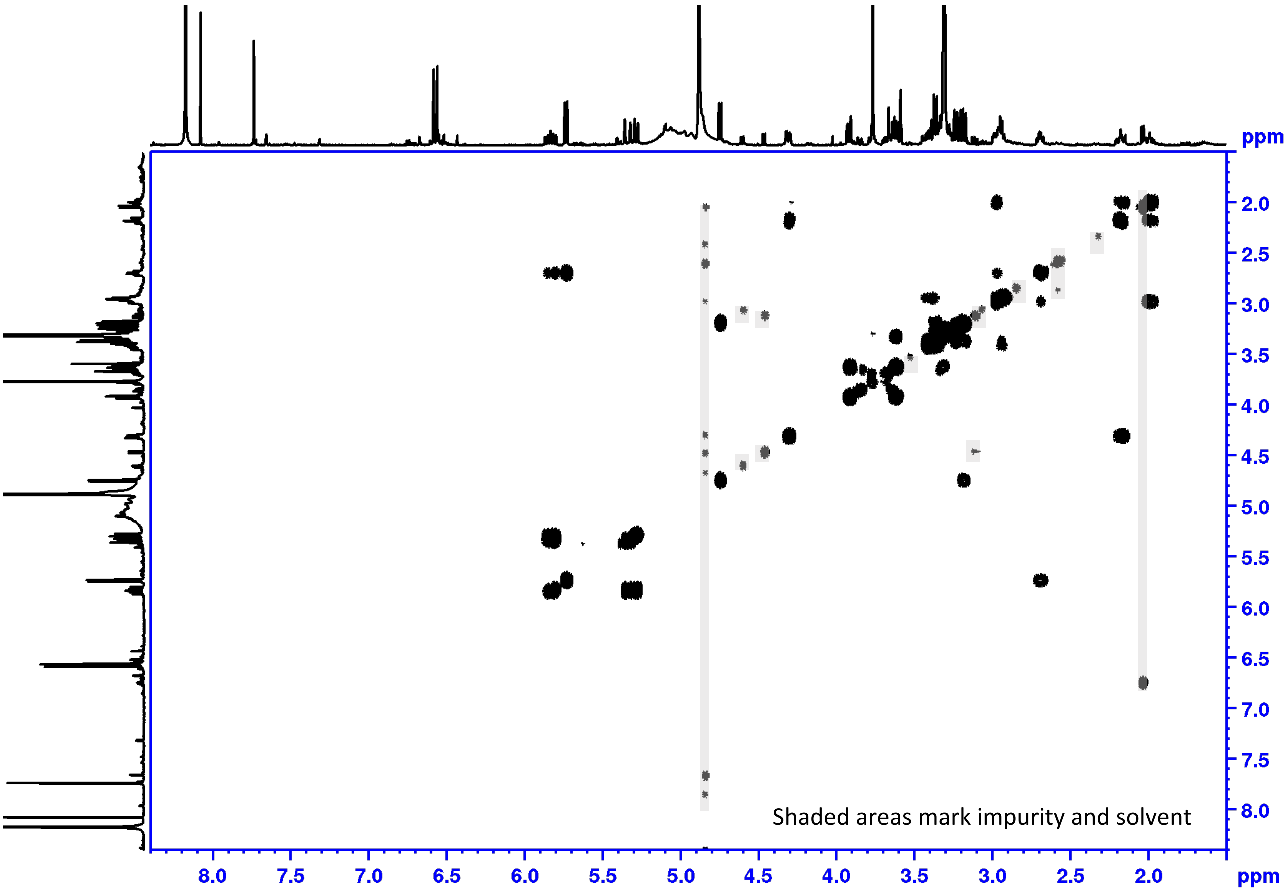

ROESY full range in MeOH-$d_3$ + 0.1% FA

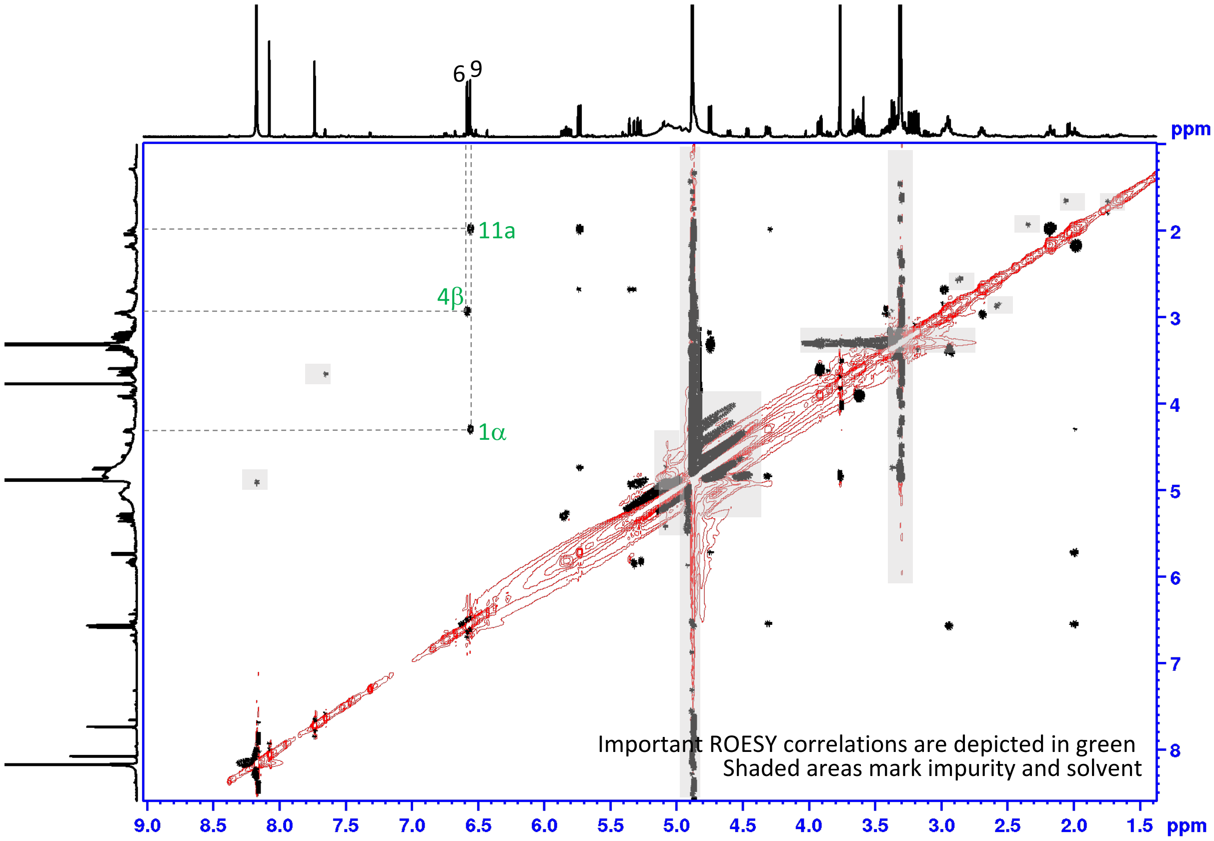

#### Supplementary Fig. 28 NMR spectra for deacetylisoipecoside

ROESY aliphatic range in MeOH-$d_3$ + 0.1% FA

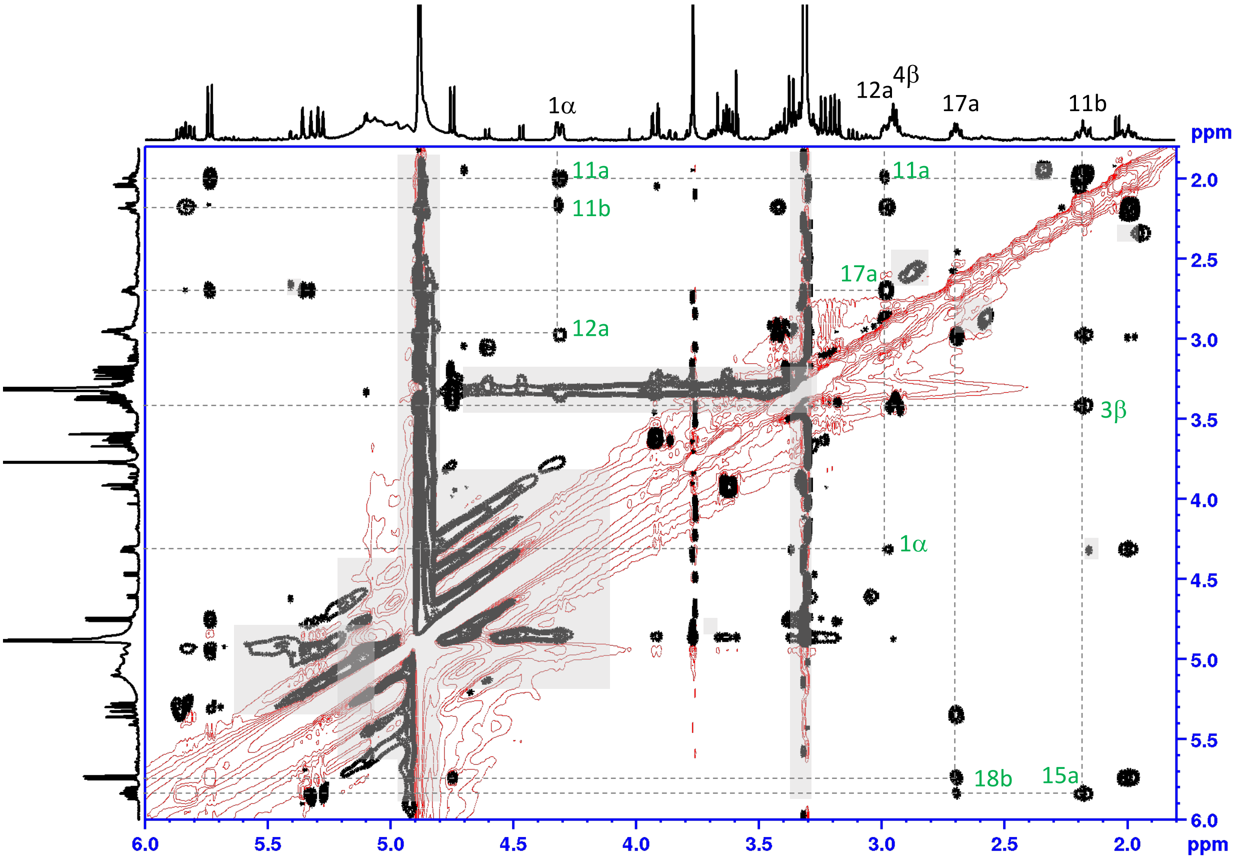

Important ROESY correlations are depicted in green.
Shaded areas mark impurity and solvent

#### Supplementary Fig. 28 ROESY correlations of deacetylisoipecoside (aglycon)

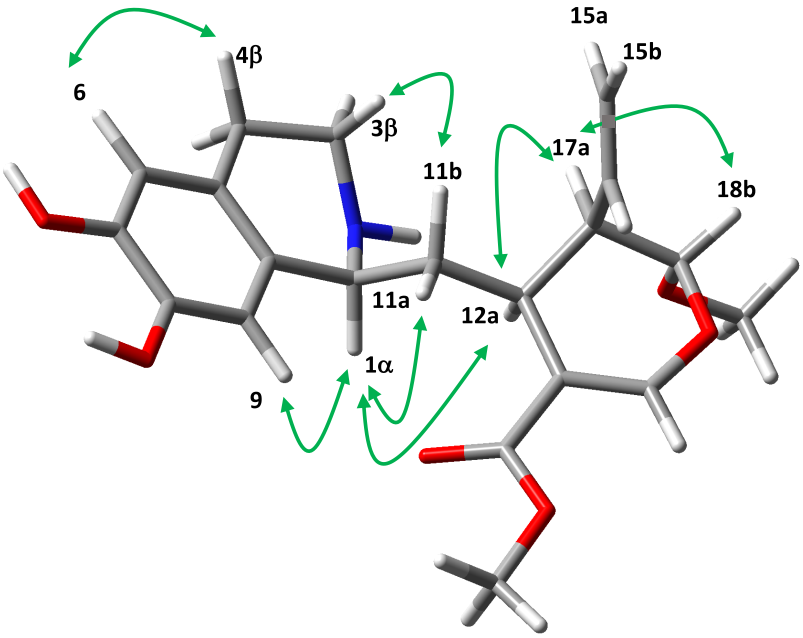

Glucose was replaced with OMe for the modeling.
Optimized using Gaussian 16W (B3LYP/6-31G(d), gas phase). Important ROESY correlations are depicted in green.

#### Supplementary Fig. 29. NMR data for demethylalangiside

#### Formed upon spontaneous lactamization of deacetylipecoside.

500 MHz NMR in MeOH-$d_3$

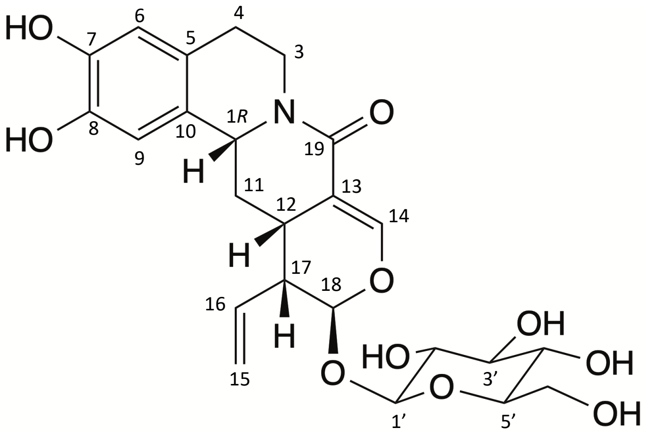

| pos. | $\delta_H$ | mult. | $J_{HH}$ | $\delta_C$ |
|---|---|---|---|---|
| 1β | 4.70 | *brdd* | 11.1/2.9 | 56.8 |
| 3α | 4.65 | *ddd* | 12.4/3.6/3.6 | 41.0 |
| 3β | 2.89 | *ddd* | 12.4/11.3/3.3 | 41.0 |
| 4α | 2.69 | *ddd* | 15.4/11.3/3.6 | 29.2 |
| 4β | 2.58 | *ddd* | 15.4/3.6/3.3 | 29.2 |
| 5 | - | - | - | 127.1 |
| 6 | 6.54 | *s* | - | 115.9 |
| 7 | - | - | - | 145.3 |
| 8 | - | - | - | 145.3 |
| 9 | 6.64 | *s* | - | 113.2 |
| 10 | - | - | - | 128.8 |
| 11α | 1.36 | *ddd* | 13.0/12.8/11.1 | 34.9 |
| 11β | 2.29 | *ddd* | 13.0/3.7/2.9 | 34.9 |
| 12β | 3.19 | *m* | - | 44.4 |
| 13 | - | - | - | 109.2 |
| 14 | 7.40 | *d* | 2.5 | 148.5 |
| 15a | 5.28 | *brdd* | 17.2/2.0 | 120.2 |
| 15b | 5.19 | *brdd* | 10.3/2.0 | 120.2 |
| 16 | 5.52 | *ddd* | 17.2/10.3/9.9 | 133.9 |
| 17β | 2.70 | *brdd* | 9.9/5.8 | 44.4 |
| 18α | 5.49 | *brd* | 1.9 | 97.3 |
| 19 | - | - | - | 165.9 |
| 1' | 4.68 | *d* | 7.8 | 99.4 |
| 2' | 3.19 | *m** | - | 74.8 |
| 3' | 3.37 | *m** | - | 78.0 |
| 4' | 3.29 | *m** | - | 71.6 |
| 5' | 3.31 | *m** | - | 78.2 |
| 6a' | 3.90 | *brd* | 11.4 | 62.7 |
| 6b' | 3.63 | *m* | - | 62.7 |

* overlapped signals *J* unresolved

About position 15a and 15 b, please see the ROESY correlation figure.

The chemical shifts agreed with the published data (Beke *et al.*, 2001).

**Supplementary Fig. 29 NMR spectra for demethylalangiside**

presaturated $^1$H NMR full range in MeOH-$d_3$

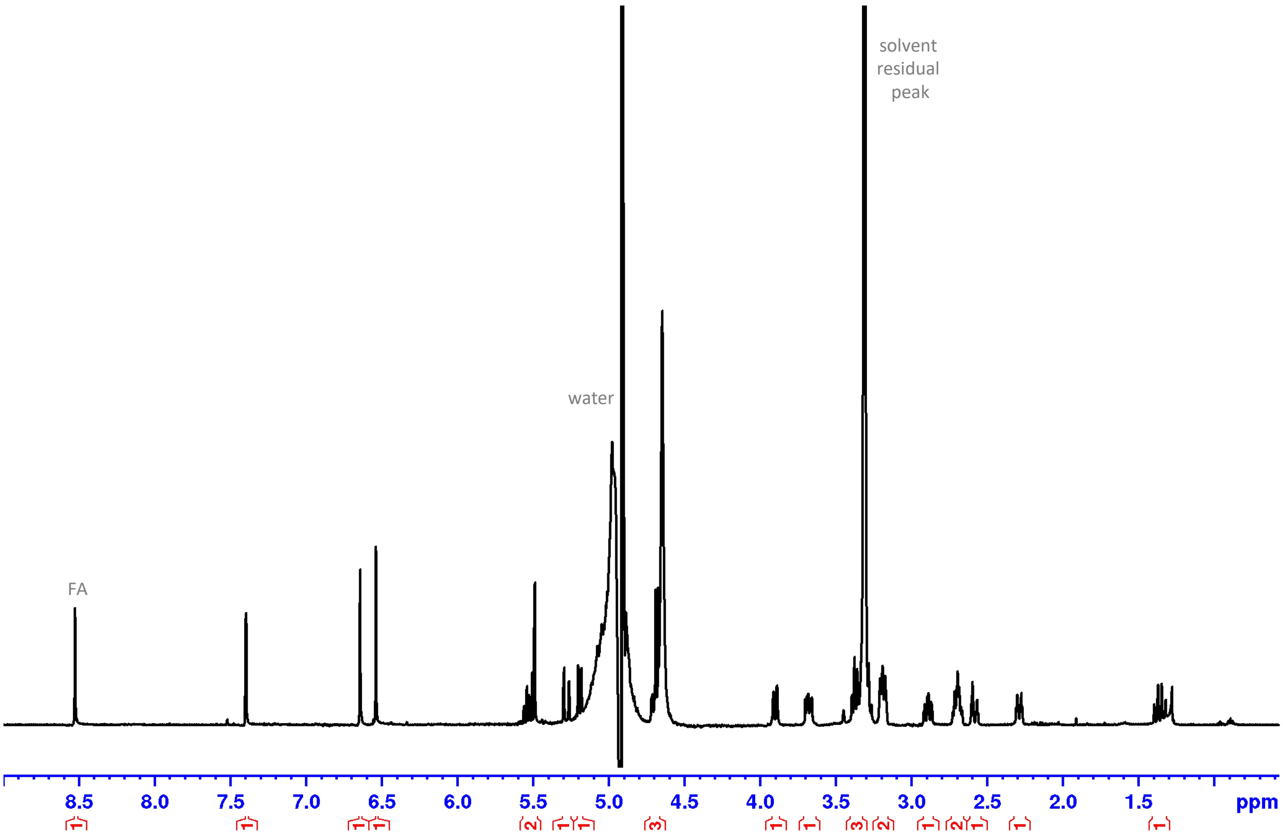

DEPTQ full range in MeOH-$d_3$

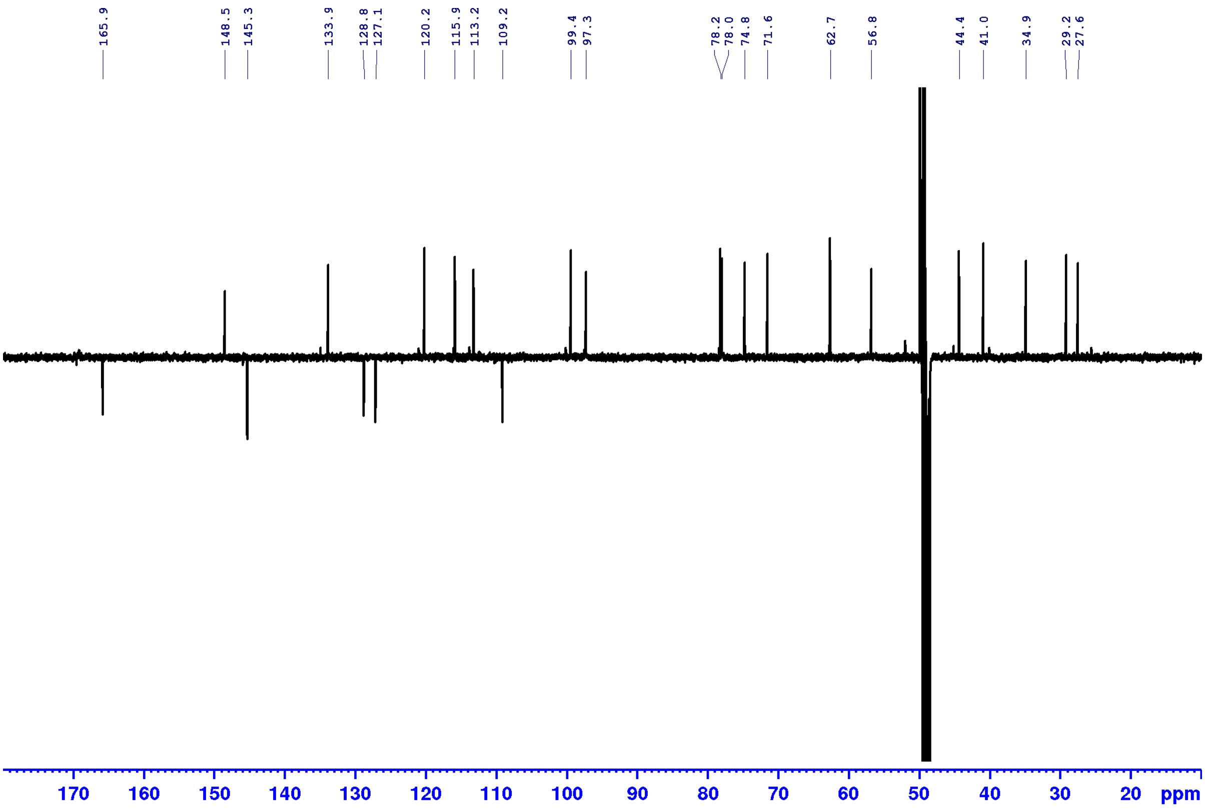

**Supplementary Fig. 29 NMR spectra for demethylalangiside**

Phase sensitive HSQC, full range in MeOH-$d_3$

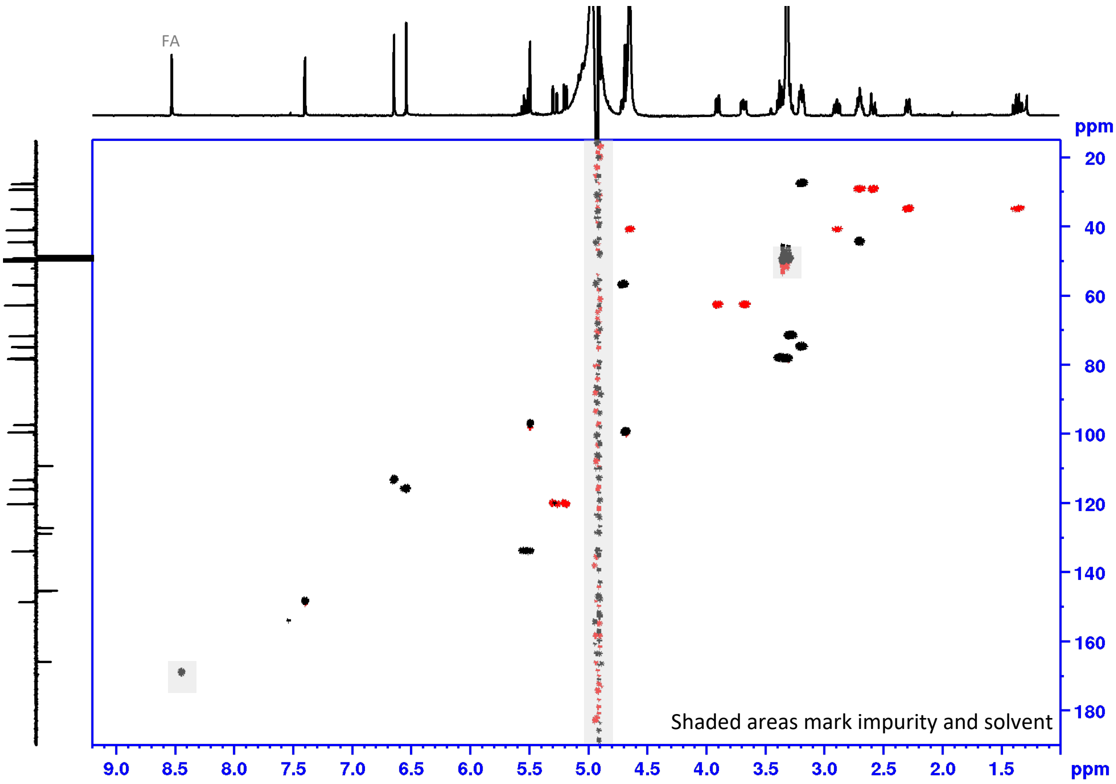

HMBC full range in MeOH-$d_3$

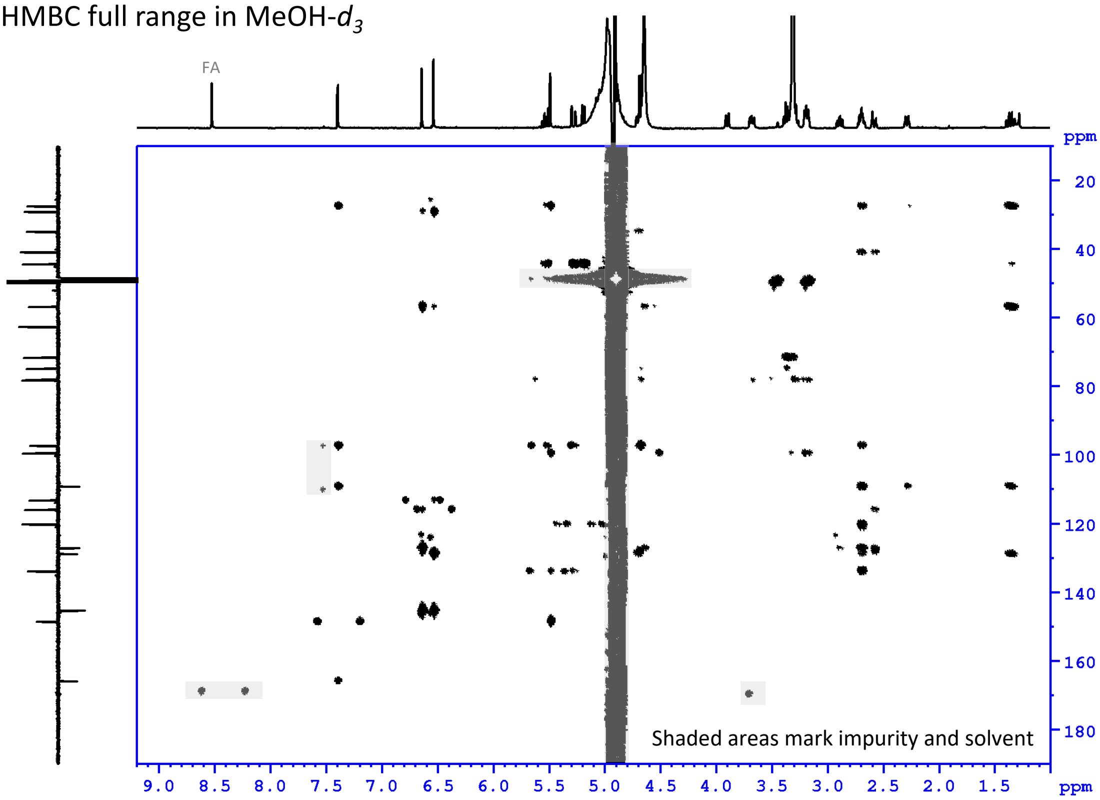

**Supplementary Fig. 29 NMR spectra for demethylalangiside**

COSY, full range in MeOH-$d_3$

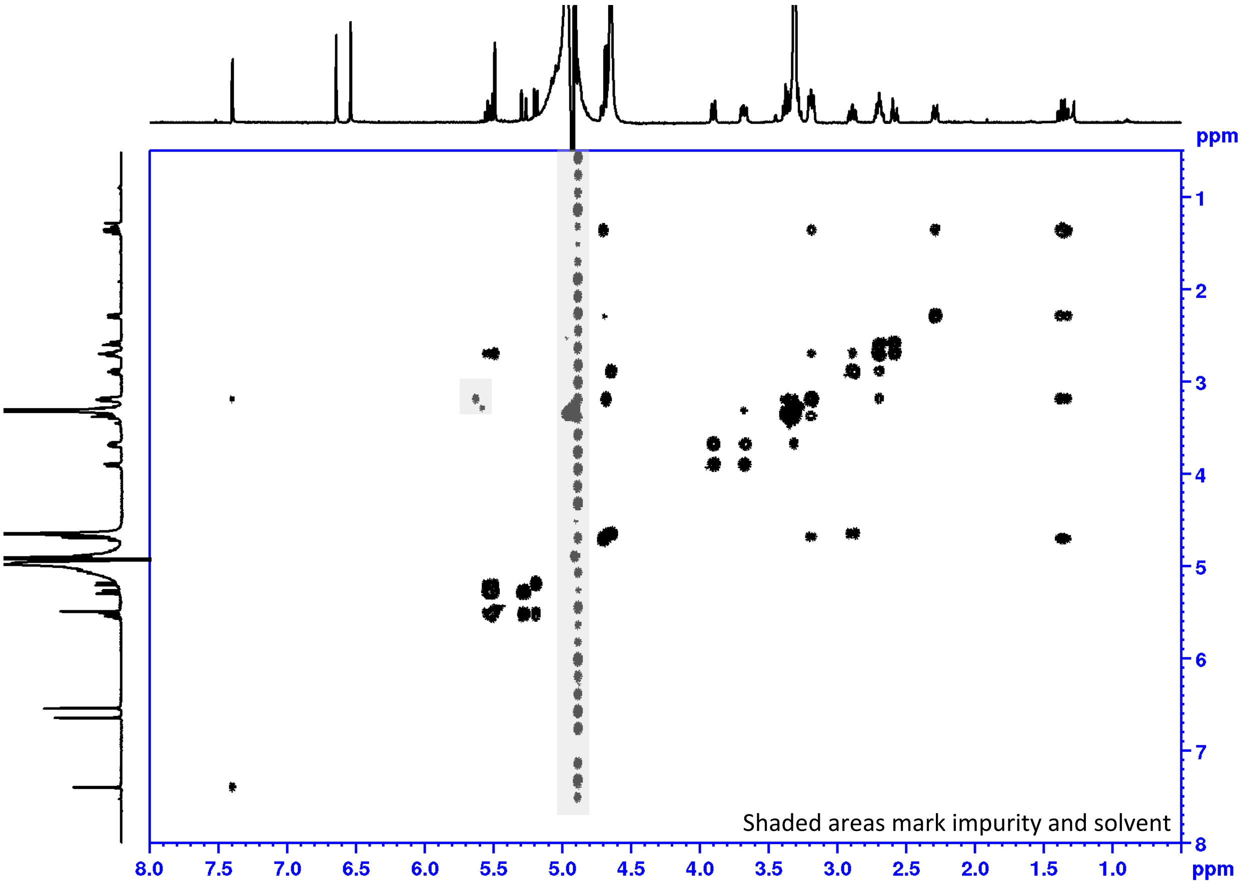

ROESY full range in MeOH-$d_3$

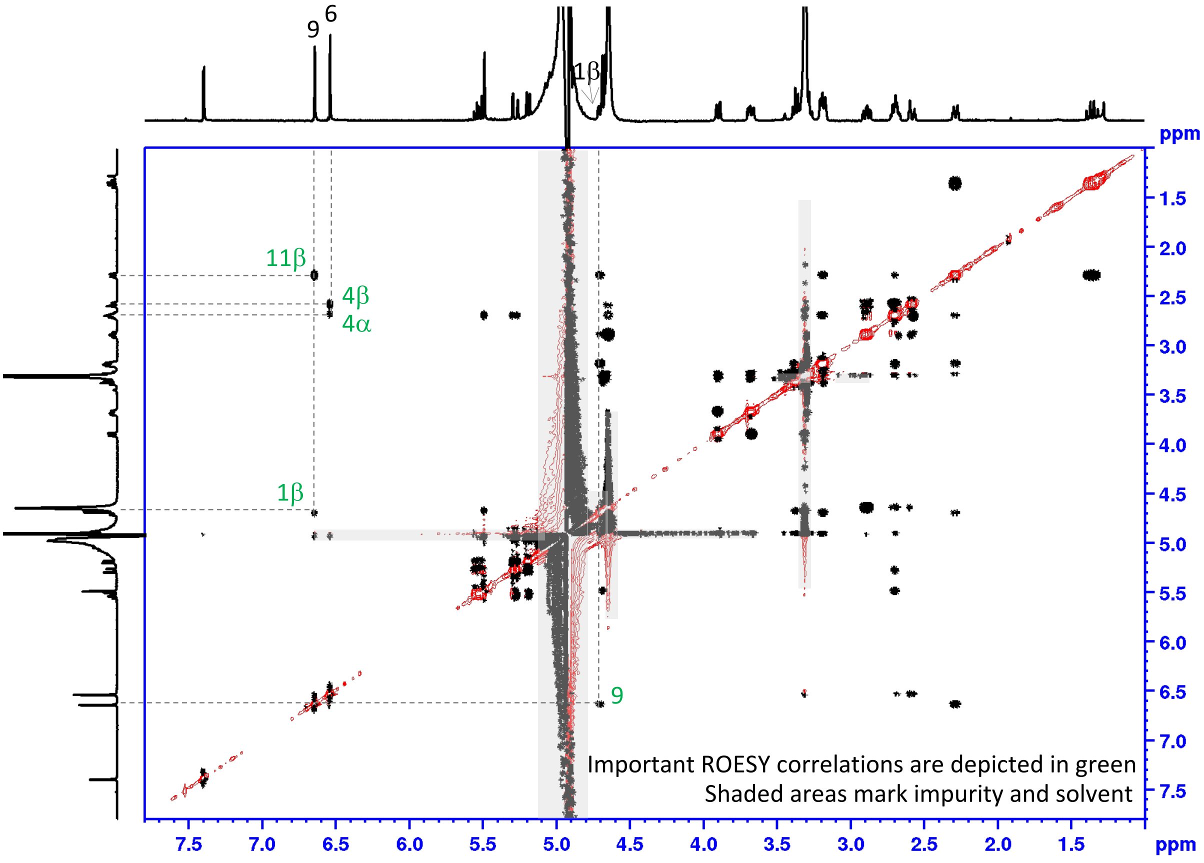

#### Supplementary Fig. 29 NMR spectra for demethylalangiside

ROESY aliphatic range in MeOH-$d_3$

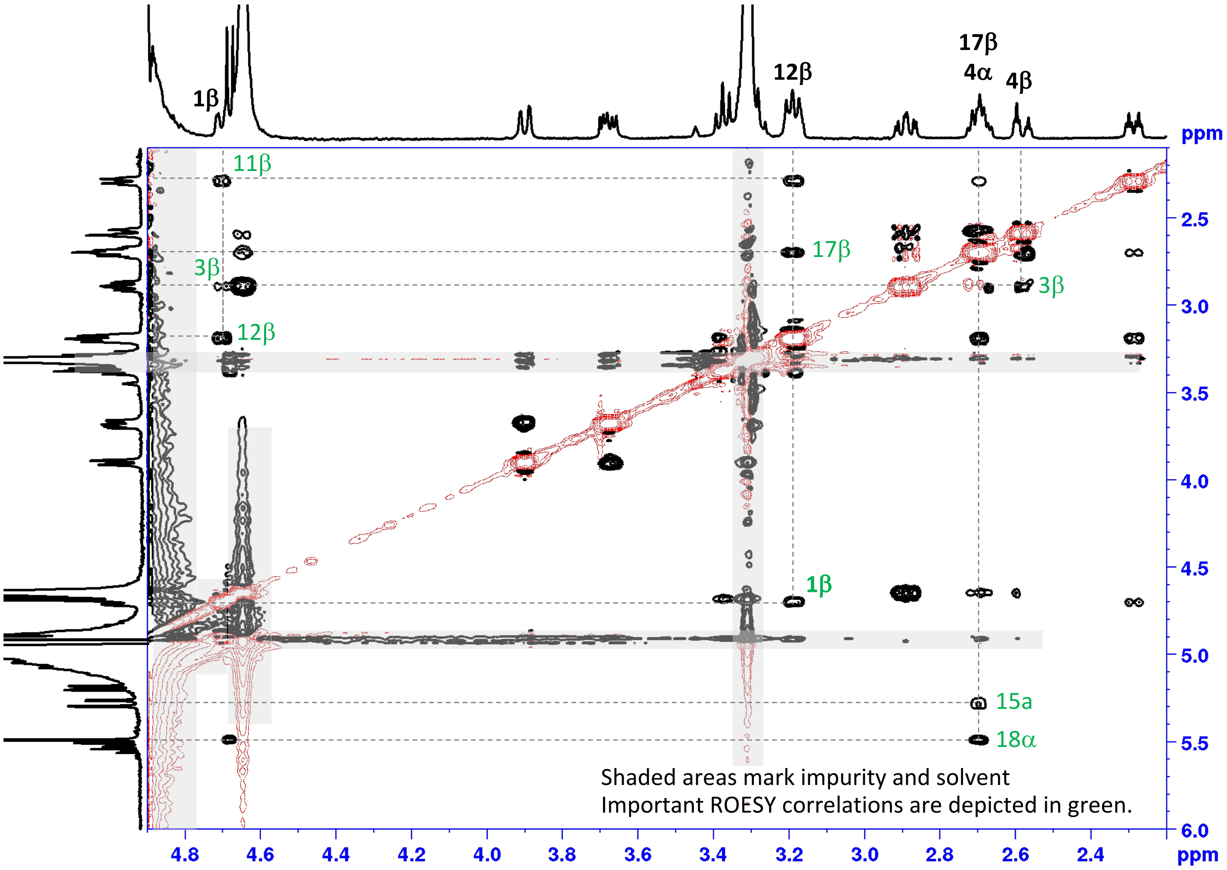

#### Supplementary Fig. 29 ROESY correlations of demethylalangiside (aglycon)

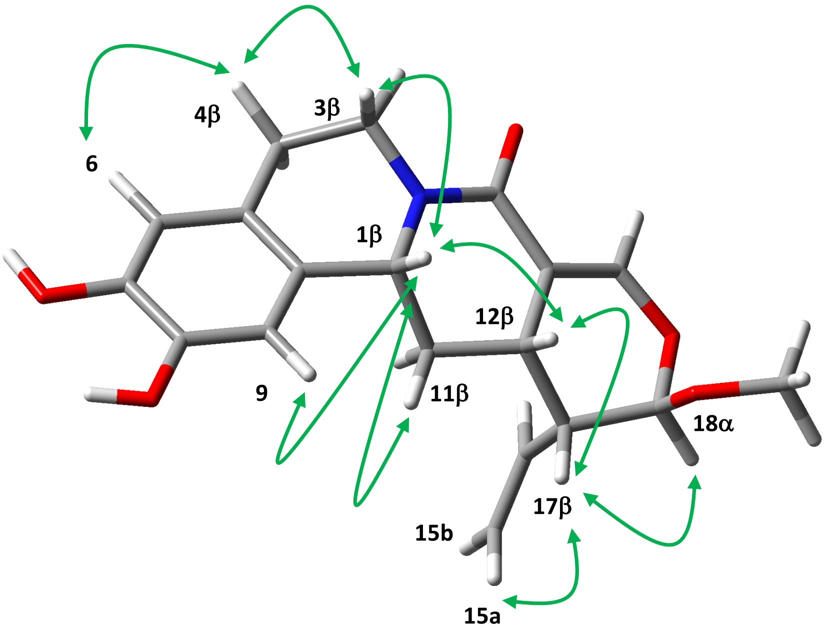

Glucose was replaced with OMe for the modeling.

Optimized using Gaussian 16W (B3LYP/6-31G(d), gas phase). Important ROESY correlations are depicted in green.

#### Supplementary Tables

#### Supplementary Table 1. List of primers used in this study.

| Gene name | Sequence | Vector |
|---|---|---|
| *CiDGD* (Nomura et al. 2008: I*peGlu1*) | TTTATGAATTTTGCAGCTCGATGTCTAGTGTTTTGCCTACCCC | Omega |
| | GACAACCACAACAAGCACCGTTAATACTTTCTTAACCTTTTCCTGGG | |
| *CiDOMT1* (Nomura and Kutchan 2010: *IpeOMT1*) | TTTATGAATTTTGCAGCTCGATGGAAACTGTCGAGAGTTCTTCC | |
| | GACAACCACAACAAGCACCGTCAAGGAGAAAGCTCCATGATACA | |
| *CiDOMT2* (Nomura and Kutchan 2010: *IpeOMT2*) | TTTATGAATTTTGCAGCTCGATGGAAACTGTTGAGAGTTCTTC | |
| | GACAACCACAACAAGCACCGTCAAGGAGAAAGCTCGATGATACA | |
| *CiDPOMT* (Nomura and Kutchan 2010: *IpeOMT3*) | TTTATGAATTTTGCAGCTCGATGGAAACTGTTGAGAGTTCTTCC | |
| | GACAACCACAACAAGCACCGTTAGTTATAAAAATTAACCTCAATGAG | |
| *CiS6DGD* | TTTATGAATTTTGCAGCTCGATGGCTACTGTTTTGGCTACCC | |
| | GACAACCACAACAAGCACCGTTACCTTCTTAATCTTTTCCTGGAGG | |
| *CiDE* | TTTATGAATTTTGCAGCTCGATGGTGGACACCACTGCAAA | |
| | GACAACCACAACAAGCACCGTTAATTGGAGTTTAGTGTCCGAGC | |
| *CiDR1* | TTTATGAATTTTGCAGCTCGATGGCACAATCACCAGAGACG | |
| | GACAACCACAACAAGCACCGTTAAGGTGATTTCAATGAGCTCATGT | |
| *CiDR2* | TTTATGAATTTTGCAGCTCGATGGCAAAATCACCGGAGACG | |
| | GACAACCACAACAAGCACCGTTAAGATGACTTCAATGAGCTCATGTC | |
| *CiIpS* | TTTATGAATTTTGCAGCTCGATGGAGATATCATCAAAGGAGTTGATT | |
| | GACAACCACAACAAGCACCGTCAGAAATAGGCAGTTGTCTCAAG | |
| *AsDOMT1* | TTTATGAATTTTGCAGCTCGATGAGTTTAATCAAAGGACCATTAAAG | |
| | GACAACCACAACAAGCACCGTCAGTAGACTCGCCTGCAGAT | |
| *AsDOMT2* | TTTATGAATTTTGCAGCTCGATGAGTTTAATGAAAGGACC | |
| | GACAACCACAACAAGCACCGTCAGGAGACTCGTCTGCAGAC | |
| *AsDOMT3* | TTTATGAATTTTGCAGCTCGATGGATAAGAAGCCAAGCAAAGGGT | |
| | GACAACCACAACAAGCACCGTCAGGAGACACGCATGCAGATA | |
| *AsDOMT4* | TTTATGAATTTTGCAGCTCGATGGACGTGACTGTCAGCA | |
| | GACAACCACAACAAGCACCGTCAGGAGAGTCGCCTGCAGAT | |
| *AsDOMT5* | TTTATGAATTTTGCAGCTCGATGGACGTGACTGTTAGCAAGG | |
| | GACAACCACAACAAGCACCGTCAGGAGAGTCGCCTGCAGAT | |
| *AsDOMT6* | TTTATGAATTTTGCAGCTCGATGAGGCCTAGCAAAGGATTGT | |
| | GACAACCACAACAAGCACCGTCAGTAGACTCGCCTGCAGA | |
| *AsDOMT7* | TTTATGAATTTTGCAGCTCGATGAGGCCTAGCAAAGGATTGT | |
| | GACAACCACAACAAGCACCGTCAGTAGACTCGCCTGCAGA | |
| *AsS6DGD* | TTTATGAATTTTGCAGCTCGATGAGCATGGACTGCGTTTAC | |
| | GACAACCACAACAAGCACCGTCAATTGGGAAGCCCCTCTT | |
| *AsDGD1* | TTTATGAATTTTGCAGCTCGATGGCGAAGACCCCATCG | |
| | GACAACCACAACAAGCACCGTTAAATACCTACTCTAGAGGCTCTTCT | |
| *AsDGD2* | TTTATGAATTTTGCAGCTCGATGGCGAAGACCCCATCG | |
| | GACAACCACAACAAGCACCGTTAAACACCCACTCTAGAGGCTC | |
| *AsDR1* | TTTATGAATTTTGCAGCTCGATGGCGAAAGCACCGGAGA | |

| | | |
|---|---|---|
| | GACAACCACAACAAGCACCGCTAGTAGGGGTTTTCAAGGTGCT | |
| *AsDR2* | TTTATGAATTTTGCAGCTCGATGGCGAAGTCGCCAGAGAC | |
| | GACAACCACAACAAGCACCGTTAGGGGGTCTTCAAGGTGT | |
| *AsDPOMT1* | TTTATGAATTTTGCAGCTCGATGAGTTGGGCCGGCGAGAT | |
| | GACAACCACAACAAGCACCGTCAATTCTTCTTCAGATATTCCATAAT | |
| *AsDPOMT2* | TTTATGAATTTTGCAGCTCGATGAGTTGGGCCGGCGAGATAA | |
| | GACAACCACAACAAGCACCGTCAATTCTTCTTCAAATATTCCA | |
| *CiDOMT1* | AAGTTCTGTTTCAGGGCCCGGAAACTGTCGAGAGTTCTTCCTC | pOPINF |
| | ATGGTCTAGAAAGCTTTATCAAGGAGAAAGCACCATGA | |
| *CiDE* | AAGTTCTGTTTCAGGGCCCGGTGGACACCACTGCAAAACA | |
| | ATGGTCTAGAAAGCTTTATTAATTGGAGTTTAGTGTCCGAGC | |
| *AsDOMT3* | AAGTTCTGTTTCAGGGCCCGGATAAGAAGCCAAGCAAAGGGT | |
| | ATGGTCTAGAAAGCTTTATCAGGAGACACGCATGCAG | |
| *AsDGD1* | CCTGACCTAGGTCTCCAATGGCGAAGACCCCATCG | alpha |
| | AAGCCTGGTCTCTGCTTAATACCTACTCTAGAGGCTCTTCTCT | |
| | CCTGACCTAGGTCTCTAAGCTGGCGAAGACCCCATCGCTC | |
| | TAGCCTGGTCTCTAAGCTTAAATACCTACTCTAGAGGCTCTTCTCT | |
| *AsDGD2* | CCTGACCTAGGTCTCCAATGGCGAAGACCCCATCG | |
| | AAGCCTGGTCTCTGCTTAACACCCACTCTAGAGGCTC | |
| | CCTGACCTAGGTCTCTAAGCTGGCGAAGACCCCATCGCTC | |
| | TAGCCTGGTCTCTAAGCTTAAACACCCACTCTAGAGGCTC | |
| *CiDGD* | CCTGACCTAGGTCTCCAATGTCTAGTGTTTTGCCTACCCC | |
| | AAGCCTGGTCTCTGCTTATACTTTCTTAACCTTTTCCTGGGG | |
| | CCTGACCTAGGTCTCTAAGCTGTCTAGTGTTTTGCCTACCCCTG | |
| | TAGCCTGGTCTCTAAGCTTAATACTTTCTTAACCTTTTCCTGGG | |
| *CiS6DGD* | CCTGACCTAGGTCTCCAATGGCTACTGTTTTGGCTACCC | |
| | AAGCCTGGTCTCTGCTTCCTTCTTAATCTTTTCCTGGAGG | |
| | CCTGACCTAGGTCTCTAAGCTGGCTACTGTTTTGGCTACCCCT | |
| | TAGCCTGGTCTCTAAGCTTACCTTCTTAATCTTTTCCTGGAGG | |
| *AsS6DGD* | CCTGACCTAGGTCTCTAAGCTGAGCATGGACTGCGTTTACAGA | |
| | TAGCCTGGTCTCTAAGCTTATCAATTGGGAAGCCCCTCTT | |
| | CCTGACCTAGGTCTCCAATGAGCATGGACTGCGTTTAC | |
| | AAGCCTGGTCTCTGCTTATTGGGAAGCCCCTCTTTGT | |
| | TTCGGTCATCAGGGACCTCTC | |
| | GAGAGGTCCCTGATGACCGAA | |
| *EYFP* | CCTGACCTAGGTCTCCAATGGTGAGCAAGGGCGAG | |
| | AAGCCTGGTCTCTGCTTCTTGTACAGCTCGTCCATGC | |
| | CCTGACCTAGGTCTCTAAGCTGGTGAGCAAGGGCGAGGAG | |
| | TAGCCTGGTCTCTAAGCTTACTTGTACAGCTCGTCCATGC | |

**Supplementary Table 2. Accession numbers of genes described in this study.**

| Gene name | Accession number |
|---|---|
| *CiDGD* | To be announced |
| *CiDOMT1* | To be announced |
| *CiDOMT2* | To be announced |
| *CiDE* | To be announced |
| *CiDR1* | To be announced |
| *CiDR2* | To be announced |
| *CiDPOMT* | To be announced |
| *CiIpS* | To be announced |
| *AsDOMT1* | To be announced |
| *AsDOMT2* | To be announced |
| *AsDOMT3* | To be announced |
| *AsDOMT4* | To be announced |
| *AsDOMT5* | To be announced |
| *AsDOMT6* | To be announced |
| *AsDOMT7* | To be announced |
| *AsS6DGD* | To be announced |
| *AsDGD1* | To be announced |
| *AsDGD2* | To be announced |
| *AsDR1* | To be announced |
| *AsDR2* | To be announced |
| *AsDPOMT1* | To be announced |
| *AsDPOMT2* | To be announced |
